# Milk-derived extracellular vesicles mitigate NF-κB pathway and NLRP3 inflammasome formation in Long Evans neonates

**DOI:** 10.1101/2025.03.30.645467

**Authors:** Jueqin Lu, Jasmyne A. Storm, Mon Francis Obtial, Sanoji Wijenayake

## Abstract

Exposure to a maternal high fat diet (HFD) during perinatal (prenatal and postnatal combined) life increases offspring’s risk of developing metabolic diseases (obesity, type II diabetes and hypertension), impairs immunity, behaviour, and neurodevelopment. Exclusive breast/chest milk feeding is a potential solution to reduce the negative developmental effects of HFD, mainly chronic systemic pro-inflammation. This study focuses on analyzing anti-inflammatory effects of a group of biological nanovesicles found in human milk, entitled milk-derived extracellular vesicles (MEVs). Specifically, we characterized the modulation of the nuclear factor κB (NF-κB) signaling pathway and NLR family pyrin domain containing 3 (NLRP3) inflammasome formation by MEVs in male and female neonatal rats with perinatal HFD exposure in the liver and hypothalamus. Female Long Evans dams were placed on a HFD or a control diet (CHD), with matching sucrose levels, 4 weeks before breeding and remained on the diets through gestation and lactation. HFD and CHD offspring received human MEVs through oral gavage twice a day from postnatal day (PND) 4 to 11. Transcript and protein abundance of candidate targets in the NF-κB signaling pathway and NLRP3 inflammasome were measured by quantitative reverse transcription polymerase chain reaction (RT-qPCR) and western immunoblotting, respectively. Our results indicate that MEV treatment attenuates the activation of NF-κB pathway and NLRP3 inflammasome formation at critical checkpoints, in males and females with perinatal HFD exposure in liver and the hypothalamus. Taken together, our data suggests that MEVs may elicit anti-inflammatory benefits postnatally that mitigates gestational HFD exposure.

## Introduction

Approximately 43.8 % of women worldwide are diagnosed with overweight (Body Mass Index (BMI) ≥ 25.0 – 29.9 kg/m^2^) or type I-III obesity (BMI ≥ 30 kg/m^2^) at the time of conception (1). Excessive dietary fat intake is strongly linked to increased risk of obesity (2). Perinatal (combined prenatal and postnatal periods) high fat diet (HFD) consumption leads to pre-eclampsia and gestational diabetes, and retention of adiposity and metabolic dysfunction post pregnancy (3). Perinatal HFD exposure also impacts the metabolic (4), immune, behavioural, and physiological health of offspring (5). Offspring with perinatal HFD have a higher tendency to develop childhood obesity, type II diabetes (6), and cardiovascular diseases (7). As described by the developmental origins of health and disease (DOHaD) hypothesis, early life malnutrition (described as intake of hyper and/or hypo caloric diets) programs offspring development, metabolism, immunity, and behaviour acutely during the exposure and chronically later in life (8).

Recent studies suggest that exclusive breast/chest feeding positively influence metabolic health of maternal and offspring systems. As the principal diets of mammalian newborns, milk contains the necessary nutrients, including lipids (9), proteins (10), and carbohydrates (11) that supports growth and development. Non-nutritive bioactive components that help establish neonatal immunity (12), gut microbiome (13, 14), and support anti-inflammatory (15) and pro-survival outcomes (16) are also present in milk (17). However, the impact of HFD-induced maternal obesity (HFD-MO) on shaping the non-nutritive components of breast/chest milk is not clear. Previous studies have reported that in milk collected from overweight females (pregnancy BMI ≥ 25.0 – 29.9 kg/m^2^) contains higher levels of saturated fatty acids that may lead to increased adiposity in offspring (18, 19). Maternal overweight status is also associated with increased abundance of candidate pro-inflammatory cytokines in milk during select lactational days (20). However, due to the plethora of beneficial bioactive components that is transferred from parents to offspring during lactation, exclusive breast/chest milk feeding is highly recommended, irrespective of the parent’s metabolic status (21). In fact, exclusive breast/chest milk feeding may revert the adverse developmental effects of gestational HFD exposure. Specifically, a group of bioactive nanovesicles enriched in human milk, entitled milk-derived extracellular vesicles (MEVs), may have the potential to combat the developmental consequences associated with perinatal diet stress (22) and promote cytoprotection (23) in offspring.

MEVs are biological nanovesicles, 30-150 nm in size, that transport lipids, peptides, proteins, deoxyribonucleic acid (DNA), messenger RNA (mRNA), and small non-coding RNA from parents to their offspring during lactation. MEVs survive intestinal degradation and cross complex biological barriers, including the intestinal epithelium and the blood-brain barrier (24). *In vitro* and *in vivo* studies have shown that MEVs readily localize in human microglia within 6-12 hours of administration and localize in the brain of adult mice within 18 hours post oral gavage and/or intravenous injections (25, 26). MEVs exert anti-inflammatory and pro-survival benefits to recipient cells and systems. Studies have shown that MEVs alleviate pro-inflammation and increase cell proliferation in intestinal systems (27, 28), lungs (29) and liver (30). MEV-microRNA (miRNA) alleviate intestinal inflammation and promotes cell proliferation (31) and mediate post-transcriptional regulation against pro-inflammatory cytokine production in intestines (32) and murine microglia (33). Thus, neuroinflammatory regulation by MEVs in rodent and human systems is starting to gain traction. However, we are yet to investigate if and how MEVs regulate nuclear factor κB (NF-κB) signaling cascade and downstream pro-inflammatory factors in female and male neonates with perinatal HFD exposure.

NF-κB is a central pro-inflammatory signaling pathway that leads to the formation of NLR family pyrin domain containing 3 (NLRP3) inflammasome and elevated production of pro-inflammatory cytokines (34). NLRP3 inflammasome is a cellular complex that modifies pro-inflammatory precursors into their functional cleaved forms and guide cells on an irreversible route to apoptosis (35, 36). NLRP3 inflammasome consists of NLRP3, apoptosis-associated speck-like protein containing a CARD (ASC) and pro-caspase-1. After NLRP3 binds with ASC, ASC recruits pro-caspase-1 and forms an activated oligomerization that leads to the self-cleavage of pro-caspase-1 into its final form, caspase-1 (37). HFD-induced obesity has been shown to promote chronic pro-inflammation through the activation of NF-κB signaling pathway (38, 39). Generally, hypercaloric intake induces metabolic inflammation by triggering the activation of the innate immune system. The activation of the innate immune responses leads to enhanced release of cytokine and chemokine signaling molecules that activates toll-like receptors (TLRs) in recipient cells, tissues, and organs (40). Multiple rodent studies have shown degrees of neuroinflammation stemming from HFD with increased expression of interleukin-1β (IL-1β) and increased apoptosis in the hypothalamus (38, 41). Neuroinflammation can lead to insulin and leptin resistance and affect metabolic homeostasis. In an HFD rat model, saturated fats in the diet were associated with increase induction of TLR pathway and autoimmune responses (42).

In this study HFD-MO is used as an early life stress to activate NF-κB signaling and NLRP3 inflammasome formation in male and female neonates to investigate if MEV treatment during critical periods of postnatal life could attenuate systemic pro-inflammation and provide pro-survival benefits. The **objective** of this study is to investigate if MEV treatment during postnatal life may rescue HFD-MO induced pro-inflammation in two metabolic tissues, liver, and hypothalamus, in male and female neonates during the stress hyporesponsive period at postnatal day (PND) 11. We **hypothesize** that MEVs will reduce the activation of NF-κB signaling and NLRP3 inflammasome formation in pups with perinatal HFD-MO and the degree of attenuation will be tissue and sex-specific.

## Maternal and methods

### Animal care

All experimental protocols were approved by the Local Animal Care Committee at The University of Winnipeg (AE18072) and falls within the guidelines of the Canadian Council on Animal Care. The same cohort of adult rats and offspring used in this study were used in a previous study (43). 7-week-old female (n = 12) and male (n = 6) Long Evans rats (strain code: 006) were purchased from Charles River (Kingston, NY, USA). Rats were housed in same-sex pairs with matching bodyweights (g). The rats were kept under a 12:12 hour light/dark cycle with *ad libitum* access to food and water.

After one week of acclimation in the housing room, a subset of females (n = 6) was randomly chosen and placed on control diet (CHD) containing 3.82 g/kCal and 10 % fat (Research diet Inc., D12450J). The remaining females (n=6) were placed on a HFD containing 5.24 g/kCal and 60 % saturated fats (Research Diet Inc. D12492) for 4 weeks (**Figure 1**) (44). The CHD has matching sucrose content to HFD. The females remained on the respective diets during a week of mating, 3 weeks of gestation, and lactation. Dam weight and caloric intake were recorded weekly. During breeding, females were housed in pairs with one control male for seven days, after which females were housed individually. Pregnancy was confirmed by monitoring maternal body weight as the presence of vaginal plugs does not always guarantee pregnancy (45). Day of parturition was considered PND 0 and the day before was considered the last day of gestation. All litters were culled to 12 pups/litter (6 males and 6 females, when possible) at PND 2 to standardize maternal care differences across litters. Note: Maternal care differences resulting from consuming this specific HFD is reported to be negligible (45, 46).

**Figure 1:**
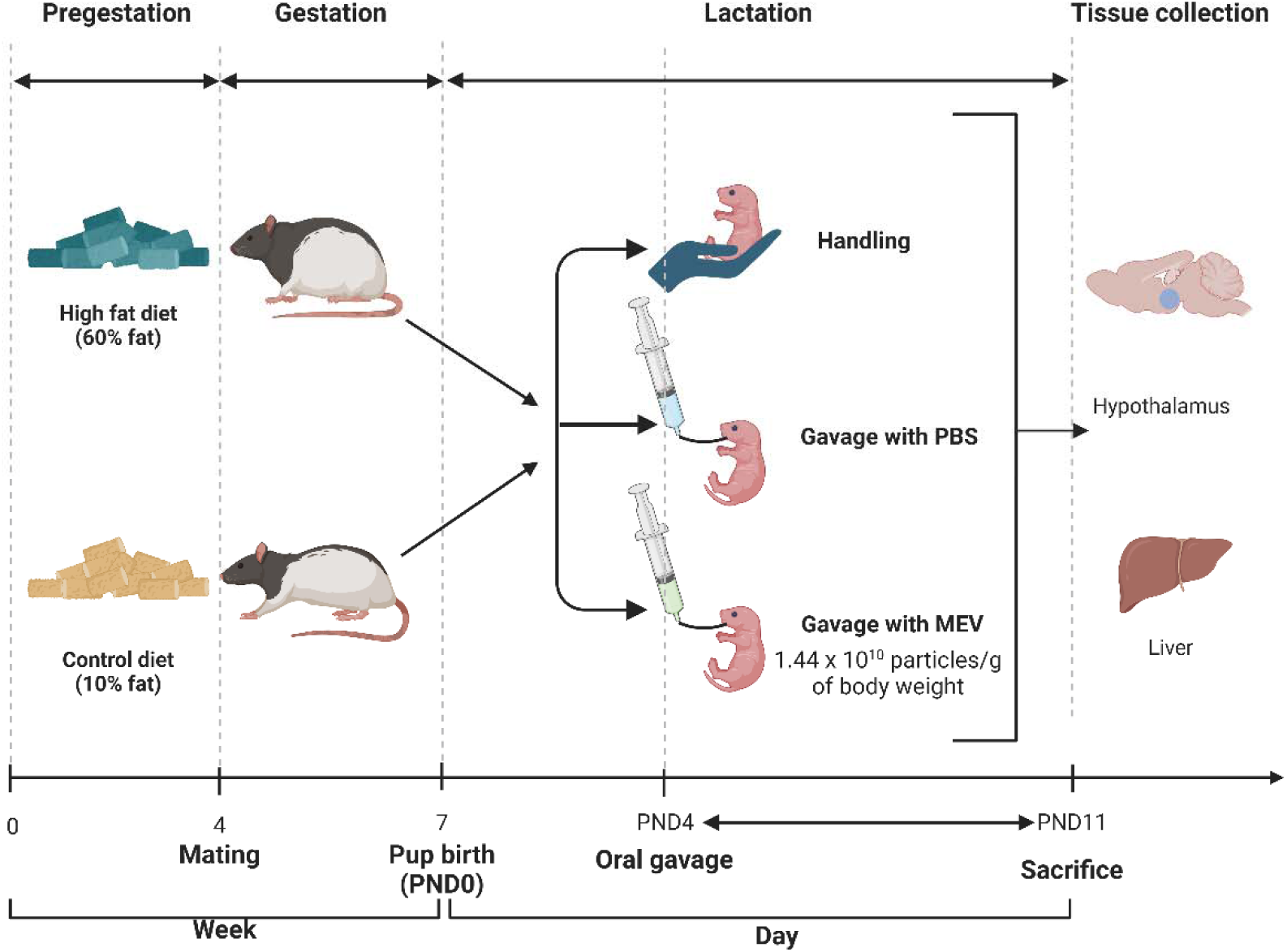
Animal care and treatment. Female Long Evans rats were placed on a control diet (CHD) consisting of 10 % saturated fat (n=6 biological replicates) or a high fat diet (HFD) consisting of 60 % saturated fat (n=6 biological replicates) for 4 weeks of pre-gestation, 3 weeks of gestation, and 11 days of lactation. Male and female offspring were culled to n=12 pups/litter with matching sex ratios at postnatal day (PND) 2. Starting at PND 4, a subset of offspring received oral gavage of MEVs, or sham, or were handled, twice a day, 6 hours apart, until PND 11 (n=4/sex/diet/treatment). Milk-derived extracellular vesicles (MEVs) dosage: 1.44×10^10^ particles per grams of body weight. Litters were sacrificed at PND 11 and liver and hypothalamus were collected for downstream analysis.

Offspring body weight and nose-anal length were recorded daily.

At PND 4, neonates were randomly selected into three treatment groups (n = 4 biological replicates/sex/diet/treatment). Group 1: neonates who received oral gavage of MEVs at a dose of 1×10^12^ per gram of body weight (termed CHD-MEV or HFD-MEV). Group 2: neonates who received a sham gavage of 1X filtered phosphate-buffered saline (PBS), at a dose of 50 µL per gram of body weight (termed CHD-PBS or HFD-PBS). And group 3: neonates who were separated from the nest and handled for the same duration of time as the neonates who received MEVs and sham (termed CHD or HFD). The MEV dosage is based on previously published literature (25, 43). Gavage experiments spanned from PND 2 to PND 11, twice a day, 6 hours apart, during the light phase to not disrupt maternal care provisions during the active phase in rodents. Each oral gavage treatment took 15 minutes from time of maternal separation to returning the offspring to the nest. This was done to ensure limited stress responses from maternal separation is induced in the neonates during treatment (47). As per previously published works, significant increases in plasmas corticosterone in neonates are observed after 2 hours of maternal separation (48, 49). Offspring were sacrificed at PND 11 via swift decapitation. Liver and hypothalamus were flash frozen and stored at −80 °C for later use. In total, 48 female and male neonates at PND 11 from n = 12 separate litters (n = 6 CHD and n = 6 HFD) were used for this study. N = 2 females and n = 2 male offspring were used per litter for analysis.

Offspring’s Lee index was calculated using the following formula (50):

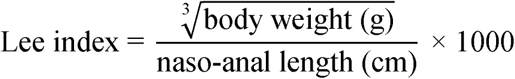

### MEV characterization

MEVs were characterized in accordance with the MISEV 2023 guidelines (51). Nanoparticle tracking analysis (NTA) (Malvern analytical, NS300), was used to measure particle concentration, particle size, and distribution at the Structural & Biophysics Core Facility at the Peter Gilgan Centre for Research and Learning (Toronto, ON, CA) as per Wijenayake et al., (2021). Samples were diluted in 1X filtered PBS (1:300, v/v). A standard curve ranging from 1:100 to 1:700 (v/v, 1X PBS) was generated to determine the optimal dilution prior to quantifications. Parameters used for the analysis: detection threshold of 10, camera level of 15, 3 replicates of 30 s capture and a green laser 532 nm.

MEV morphology and integrity was characterized using transmission electron microscopy (TEM: FEI Talos F200x) as per Wijenayake et al., (2021) at the Manitoba Institute of Materials (Winnipeg, MB, CA). Briefly, MEVs were diluted in 1X filtered PBS (1:3, v/v) and crosslinked to 400 mesh carbon-coated copper grids (Electron Microscopy Sciences, CF400-CU) and incubated at room temperature for 15 minutes. The grids were washed 3x with 50 µL of ddH_2_O and wadded dry with filter paper to remove excess sample and water. The carbon grids were negatively stained with 2 % uranyl acetate. The grids were air-dried for 15 minutes and visualized immediately.

Western immunoblotting was used to validate the presence of EV biomarkers (**Table 1**). MEV pellets were lysed with 100 μL of lysis buffer containing protease inhibitor cocktail with ethylenediaminetetraacetic acid (EDTA) (Bioshop, PIC001.1) (1:5000, v/v) and Phenylmethanesulfonyl fluoride (Bioshop, PMS123.5) in cell extraction buffer (Thermofisher Scientific, FNN0011),). After 30 minutes of incubation with intermittent vortexing, samples were centrifuged at 13,000 rpm at 4 C for 10 minutes to isolate protein samples. Pierce BCA protein assay (Thermofisher Scientific, ZB385879) was used to determine the protein concentration.

Protein samples were normalized to a uniform concentration with the cell extraction buffer and combined with 2X SDS (1:1 v/v) (2X SDS, 4 % SDS v/v, 20 % glycerol v/v, 0.2 % bromophenol blue w/v, 100 mM of Tris base in ddH_2_O) and 10 % β-mercaptoethanol (v/v) and denatured by heating at 95 C for 10 minutes (52). The abundance of two tetraspanins (CD9 and CD81) and an EV biogenesis marker (syntenin-1) were measured in soluble proteins isolated from human MEVs. Calnexin was used as the cellular marker (the negative control). Total soluble protein from HMC3 cells were resolved along side MEV protein lysates to provide insight into cellular distribution and EV distribution of these markers. Antibodies from the EV characterization panel was used for analysis (Cell signaling, 74220T).

### RNA extraction, cDNA synthesis, and quantitative reverse transcription polymerase chain reaction

RNA was extracted from neonatal liver and hypothalamus using TRIzol reagent (Thermofisher Scientific, 15596026) according to manufacturer’s instructions. Briefly, 50-100 μg of frozen tissue was powdered with liquid nitrogen using a motor and pestle and homogenized using an 18-gauge needle with 1 mL of ice cold TRIzol (w/v). The resulting supernatant containing nucleic acids were loaded onto RNA Clean and Concentrator columns (Zymo Research, R1017) and processed as per manufacture’s instructions. Nanodrop One/One^C^ Microvolume-UV/Vis spectrophotometer (ThermoFisher Scientific: ND-ONE-W) was used to determine RNA concentration (ng/μL) and purity (A260:A280 and A260:A230). Samples with A260:A280 and A260:A230 ratios between 1.5-2.0 were used for complementary DNA (cDNA) synthesis. Samples with A260:A280 and A260:A230 lower than 1.5 were further cleaned with RNA clean & concentrator kit (Zymo Research, R1017). The integrity of RNA samples was tested by resolving the samples on a 1 % TAE agarose gel (1:1, v/v, 2x RNA loading dye) (Life technology, R0641) stained with Red Safe (Froggabio, 21141) as per Storm et al., (2025). The gel was resolved with a Sub-Cell GT Agarose Gel Electrophoresis system (BioRad, 1704401) at 130 V for 45 minutes. Extracted RNA was converted to cDNA using the High-Capacity Reverse Transcription kit (Applied Biosystems, 4368814) using the T100^TM^ Thermal Cycler (BioRad, 1861096).

Primers (**Table 2**) were designed with nucleotide sequence information available at the National Centre for Biotechnology Information (NCBI): http://www.ncbi.nlm.nih.gov or chosen from published literature. Primer sequences were generated using Primer-BLAST (National Library of Medicine) using the coding sequence of target genes and tested with OligoAnalyzer (Integrated DNA Technologies) using the following parameters: % GC levels = 40-60 %, melting temperature = 55-65 C, hairpin temperature < 45 C, self-dimer ΔG < 15 %, and heterodimer ΔG < 20 %. Primers were purchased from Eurofins Genomics (Louisville, KY, USA).

Absolute transcript abundance in the form of quantity means (QM) were measured via quantitative reverse transcription polymerase chain reaction (RT-qPCR) with Fast SYBR Green PCR Chemistry (Applied Biosystems, 4385612). Targets analyzed: *toll-like receptor 4 (TLR4),* I *kappa B kinase complex* (*I*κκβ)*, NF-kappa-B inhibitor alpha* (*I*κ*B*α)*, NF-*κ*B, NLRP3, interleukin-18 (IL-18),* and *caspase-1*. A temperature curve was performed using a pool sample containing all tested samples to determine annealing temperature per primer pair. An 8-point standard curve ranging from 500 ng/μL to 3.91 ng/μL consisting of female and male samples across treatment groups was performed to determine the optimal cDNA loading amount per primer pair. Melt curve analysis was conducted to determine the specificity of target primers, where primer pairs with a single sharp melt peak were used for quantification. Three candidate reference genes were tested, including glyceraldehyde-3-phosphate dehydrogenase (*GAPDH*), tyrosine 3-monooxygenase activation protein zeta (*YWAZ*), and glucuronidase beta (*GUSB*). RT-qPCR was performed with 3 technical triplicates per biological replicate and 4 biological replicates/sex/diet/treatment. RT-qPCR protocol used in the study: 95 C for 20 s followed by 40 cycles amplification (95 C for 3 s and 60 C for 30 s) and 95 C for 15s and 60 C for 69 s for melt curve and ended with 95 C holding step for 15 s.

### Protein extraction and western immunoblotting

Total soluble protein was extracted from liver and hypothalamus from the same female and male offspring that were used for RT-qPCR using a lysis buffer, 1:5 (w/v) (Miliplex, 43-040) as per manufacture’s instructions. The lysis buffer was supplemented with protease inhibitor cocktail with ethylenediaminetetraacetic acid (Bioshop, PIC001.1), 1mM of sodium orthovanadate (Bioshop, SOV850.25), 10 mM of sodium fluoride (Bioshop, SF001), and 10 % (v/v) β-mercaptoethanol (Bioshop, MER002.500) (53). Samples were incubated on ice for 30 minutes with intermittent vortexing and centrifuged at 14,000 rcf at 4 C for 20 minutes. Protein concentration was measured using a bicinchoninic acid assay (BCA) (Thermo Scientific, ZB385879), and quantified with a multimode microplate reader (BioTek Synergy, H1M2). Bromophenol blue loading dye containing SDS (Bioshop: SDS001.1) and β*-*mercaptoethanol (10 %, v/v) (Bioshop: MERC002.500) were added to the samples 1:1 (v/v). The samples were vortexed and heated to 95 °C for 10 minutes to denature the proteins. Converted lysates were stored at −20 °C.

Protein abundance of TLR4, phosphorylated-Iκκ α/β (Serine 176 and 180), phosphorylated-IκBα (Serine 32) NF-κB, phosphorylated-NF-κB (Serine 536), NLRP3 and caspase-1 were determined using western immunoblotting (Table 3). Briefly, 10 - 30 µg of soluble proteins were resolved on 6 %, 8 % and 12 % SDS-Acrylamide gels at 180 V for 70 minutes (BioRad, PowerPac Basic) with 5 μL of protein ladder (FroggoBio, PM008-0500). A rodent liver sample was resolved alongside the target samples in all immunoblots and used as the interblot converter to enable statistical comparisons across immunoblots. The percentage of the resolving gel was based on the predicted molecular weight of the protein targets. A standard curve with 5 – 35 μg of organ specific pool sample was used to determine the optimal protein amount to load for each target. Proteins were electroblotted to a 0.45 µM PVDF membrane (BioRad, 10026934) using the Fast transfer system (BioRad, Trans-Blot Turbo transfer system, 1704150). Post-transfer, the gels were stained with Coomassie blue solution (0.25 % of Coomassie brilliant blue G-250, Bioshop 6104581; 7.5 % of acetic acid, Fisher Chemicals 64197 and 50 % of methanol, Bioshop 67561) and destained with destaining solution (10 % of acetic acid and 25 % of methanol). This was done to ensure successful transfer of proteins in the predicted molecular weight. 1-10 % casein (v/v, 1X Tris-buffered saline with Tween 20) was used as the blocking reagent to minimize unspecific binding and membranes were incubated with primary and secondary antibody (Bioshop, APA007P.2) as per conditions described in **Table 3**.

1X Tris-buffered saline with Tween 20 was used for all washing steps. Membranes were visualized with SuperSignal^TM^ West Pico PLUS Chemiluminescent Substrate (Thermo Scientific, 34580) and the chemiluminescent signals were captured with Bio-Rad ChemiDoc™ MP imaging system (12003154) using Image Lab Software (version 6.1). The immunoblots were stained with Coomassie brilliant blue stain (0.25 % of Coomassie brilliant blue G-250, 7.5 % of acetic acid, and 50 % of methanol) and destained with the destaining solution. Immunoblot band intensities were normalized against the summed intensity of Coomassie stained protein bands in the same lane (without including the target band) to normalize against protein loading irregularities. This method was shown to be more accurate than using a single constantly expressed protein target for normalization (54). Band intensities were quantified using ImageJ (version 1.5.4).

### Statistical analysis

Statistical analysis was done using SPSS (IBM version 29) and all figures were constructed using GraphPad Prism (version 8.4.2) and BioRender software. Prior to conducting parametric analyses, normality was assessed using the Shapiro-Wilk test (55). and Levene’s test was used to test the homogeneity of variance (p>0.05). When data were not normally distributed, extreme outliers (3 times interquartile range of the mean) were examined using the SPSS outlier boxplot function and removed from the dataset, only when necessary. No more than n=1 extreme outlier were removed per treatment group. qPCR and western immunoblotting data were analyzed within sex and tissue.

A factorial general linear model (GLM) univariate (sex, diet, MEV treatment) was used to analyze transcript and protein data (n=4 biological replicates/sex/diet/treatment/tissue). To conduct pairwise comparisons between diet and treatment groups, Tukey’s honestly significant test (Tukey HSD) was used. Data that did not achieve normality was analyzed with Kruskal-Wallis H test followed with Dunn’s post hoc test or Mann Whitney U test. A p-value<0.05 was considered statistically significant for all analyses. Data are represented as mean ± standard error of means (SEM).

## Results

### Characterization of MEV morphology, particle size distribution, and extracellular markers

To characterize MEV morphology in samples, TEM with 2 % UA negative staining was used (**Figure 2A**). A majority of MEVs retained the circular nanovesicle structure with intact cargo. The exterior membrane and internal cargo were successfully stained. The pellet fraction was relatively clear of cellular debris but contained some protein aggregation. Due to the complexity and heterogeneity of unpasteurized human milk, the presence of protein aggregation in the enriched MEV fraction is expected, as reported previously (15, 56, 57). The corresponding supernatants contained little to no MEVs (**Figure 2B**). The diameter of MEVs in samples is between 50 - 130 nm.

**Figure 2:**
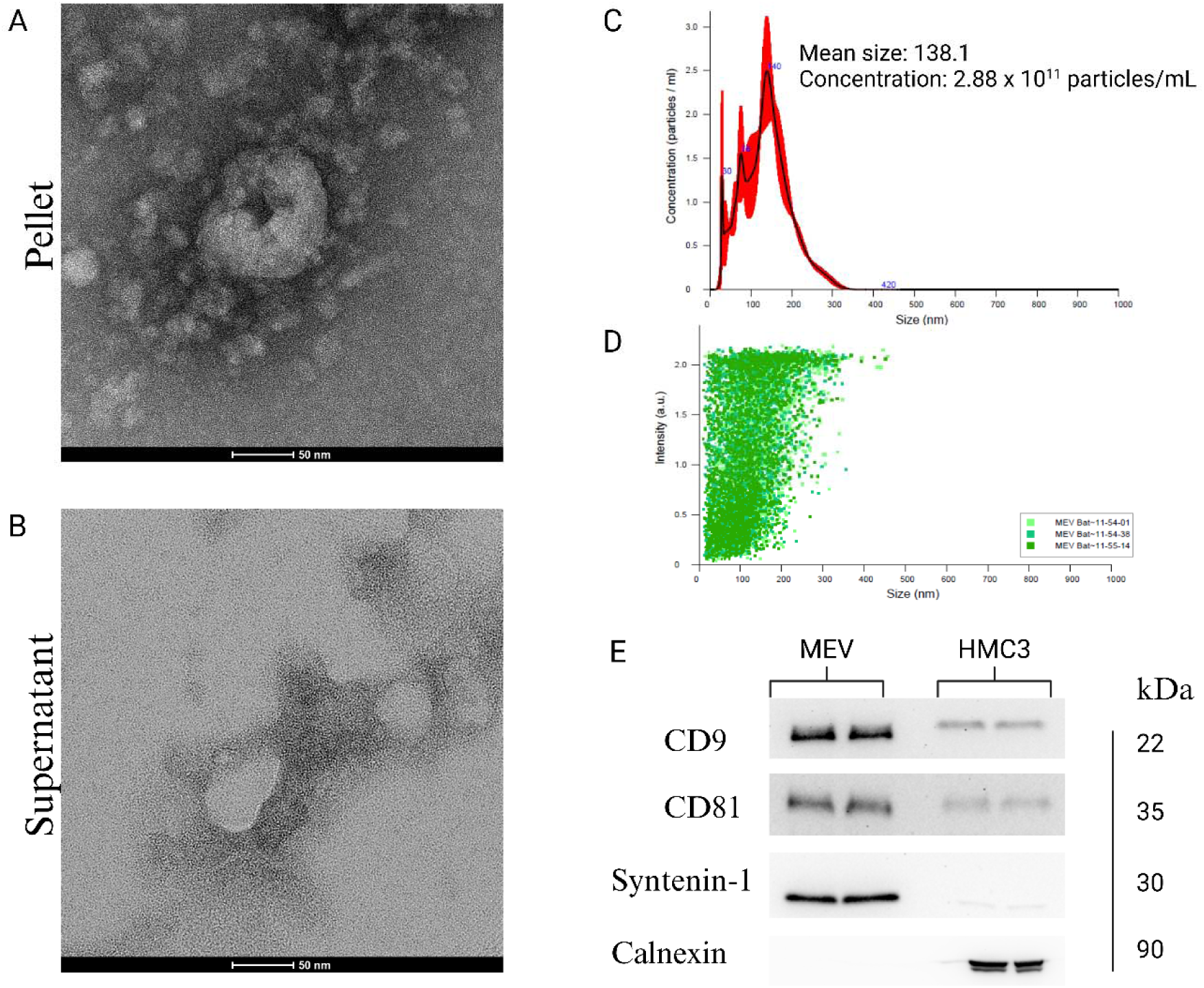
Milk-derived extracellular vesicles (MEVs) characterization. The characterization was done as per Minimal Information for studies of Extracellular vesicles (MISEV) 2023 guidelines (50). (**A**) MEV morphology as determined by transmission electron microscopy (TEM). Scale bar = 50 nm. (**B**) The supernatant from MEV isolations were used as the negative control void of MEVs. (**C**) MEV particle size and concentration as determined by nanoparticle tracking analysis (NTA). A mean particle size of 138.1 nm and a concentration of 2.88 x 10^11^ particles/mL were determined. (**D)** MEV particle distribution depicted by NTA. (**E**) Protein abundance of three extracellular vesicle markers (CD9, CD81, and Syntenin-1) and an endoplasmic reticulum marker used as the negative control (Calnexin), as determined by western immunoblotting. The total soluble protein isolated from human microglia clone 3 (HMC3) cells represent protein abundance in cells versus that of extracellular vesicles. The complete immunoblots of the targets and Coomassie stained membranes (loading control) are included in **Figure S1**.

The size and particle distribution of MEVs were tested with NTA (**Figure 2C**). The mean particle size of the MEVs was determined to be 138.1 nm and the concentration of MEVs was calculated to be 2.88 x 10^11^ particles/mL.

Western immunoblotting was performed to confirm the presence of MEV markers, including CD9, CD81, and syntenein-1 (**Figure 2D**). CD9 and CD81 are tetraspanin proteins that are located on the surface of extracellular vesicles and syntenin-1 is involved in EV biogenesis. The abundance of CD9, CD81, and syntenin-1 confirmed the presence of MEV in our samples.

The three markers were also abundant in the HMC3 cells, albeit in lower levels, compared to MEV samples. The expression of CD9, CD81, and syntenin-1 in microglia is expected as these markers regulates cell adhesion, cell to cell communication, and cell motility (58). The endoplasmic reticulum marker, Calnexin, was prominently abundant in the microglia lysates and was absent in the MEVs, illustrating that the MEV isolations were void of cellular contamination. The complete immunoblots of the targets and Coomassie stained membranes (loading control) are included in **Figure S1.**

### Transcript and protein abundance of NF-kb pathway targets in neonatal liver in response to diet and MEV treatment

*TLR4* transcript abundance (**Figure 3A**) in male neonates remained unchanged in response to HFD exposure (*F* (_1, 12_) = 0.334, p = 0.574) and MEV gavage (*F* (_3, 12_) = 0.983, p = 0.433). *TLR4* transcript in females was influenced by diet (*F* (_1, 11_) = 7.326, p = 0.020), and MEV gavage (*F* (_3, 11_) = 3.913, p = 0.040), where HFD-MEV neonates had lower levels of *TLR4* compared to CHD-MEV neonates (p = 0.027). *TLR4* level was not influenced by sex (*F* (_1, 23_) = 0.788, p = 0.384). At the protein level, (**Figure 3B**), a main effect of sex was seen, with a higher abundance of TLR4 in females than males (*F* (_1, 24_) = 27.357, p<0.001). In males, TLR4 levels remained unchanged in response to diet (*F* _(1, 12)_ = 0.087, p = 0.773) or treatment (*F* _(3, 12)_ = 2.261, p = 0.134), but in females, there was a main effect of MEV treatment (*F* _(3, 12)_ = 21.992, p<0.001) and diet (*F* _(1, 12)_ = 35.541, p<0.001). HFD females who received MEV gavage had lower levels of TLR4 protein compared to HFD females who did not receive MEVs (p<0.001). Also, TLR4 protein levels were higher in HFD females compared to CHD females (p<0.001).

**Figure 3:**
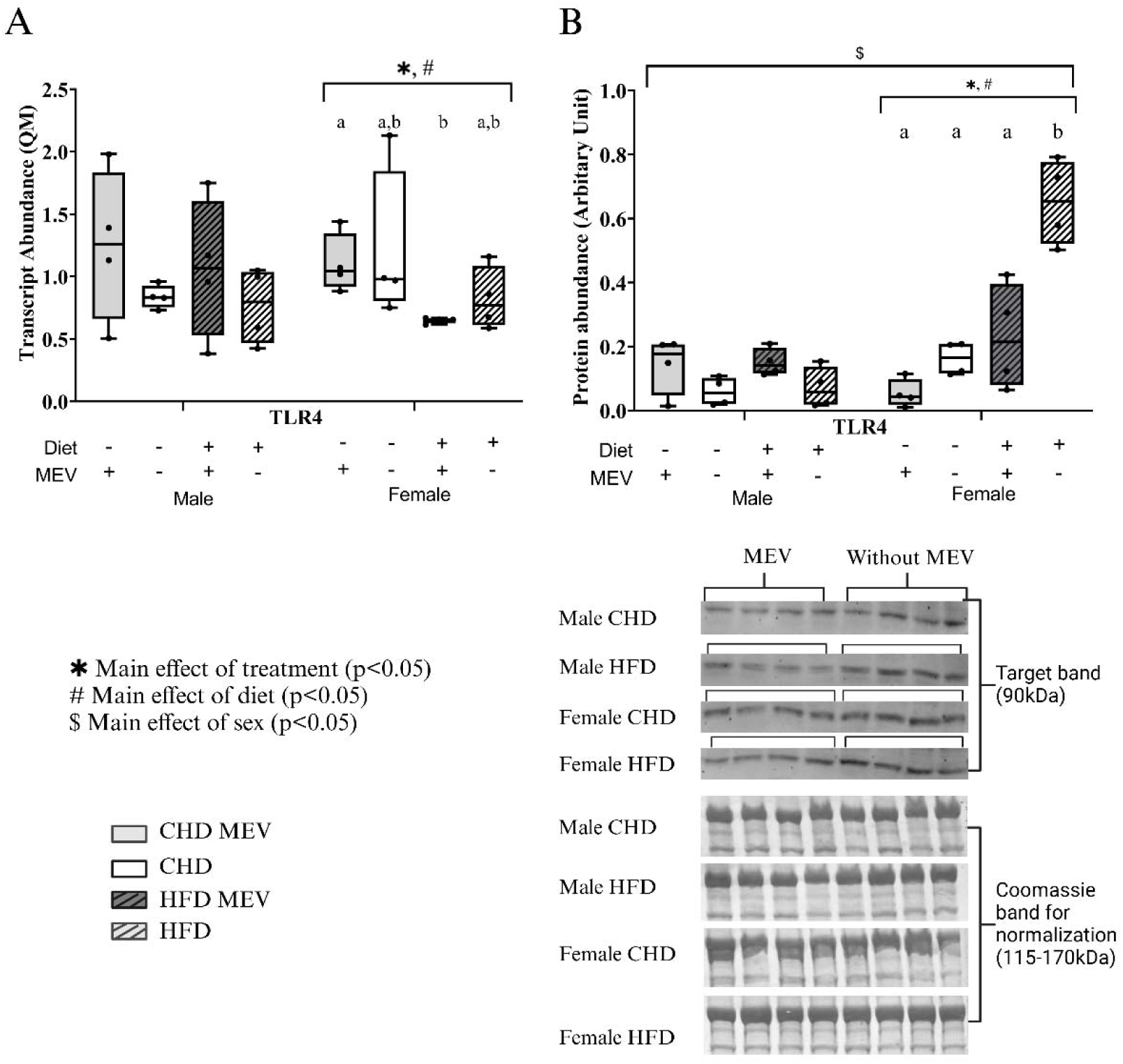
Transcript abundance (Quantity Means) and protein abundance (normalized to Coomassie stain) of NF-κB pathway markers as determined by RT-qPCR and western immunoblotting, respectively in the liver of male and female neonates at postnatal day (PND) 11. (**A**) *TLR4* transcript (**B**) TLR4 protein. CHD MEV = control offspring that received milk-derived extracellular vesicle (MEV) treatment from PND 4-11. CHD = control offspring that did not receive MEV treatment. HFD MEV = offspring with perinatal high fat diet exposure and received MEV treatment from PND 4-11. HFD = offspring with perinatal high fat diet exposure that did not receive MEV treatment. MEV dosage: 1.44×10^10^ particles per grams of body weight. Data are presented with standard error of means. N = 4 biological replicates/ diet/treatment/sex. * Main effect of MEV treatment. # Main effect of diet. $ Main effect of sex. Same letters above the boxplots represents no changes and different letters represent significant difference as determined by Tukey HSD. Effects are statistically significant at p<0.05.

*IKK*β (**Figure 4A**) transcript abundance remained unchanged in males in treatment (*F* _(3, 12)_ = 0.814, p = 0.511) and diet (*F* _(1, 12)_ = 1.340, p = 0.270) and females (*F* _(3, 12)_ = 2.339, p = 0.125) (*F* _(1, 12)_ = 1.253, p = 0.285). No main effect of sex was observed in transcript level as well (*F* _(1, 24)_ = 0.106, p = 0.748). At the protein level, (**Figure 4B**) there was a main effect of sex (*F* _(1, 24)_ = 25.446, p<0.001), where females had higher abundance of p-Iκκ α/β (S176/180). In both sexes, there was a main effect of diet (*F* _(1, 12)_ = 19.215, p<0.001 in male samples) (*F* _(1, 12)_ = 7.678, p=0.017 in female samples) and MEV treatment (*F* _(3, 12)_ = 6.409, p = 0.008 in male samples) (*F* _(3, 12)_ = 3.693, p = 0.043 in female samples). In males, HFD exposure (irrespective of MEV treatment) led to an increase in p-Iκκ α/β (S176/180) compared to CHD-MEV (p = 0.044) and CHD (p = 0.036).

**Figure 4:**
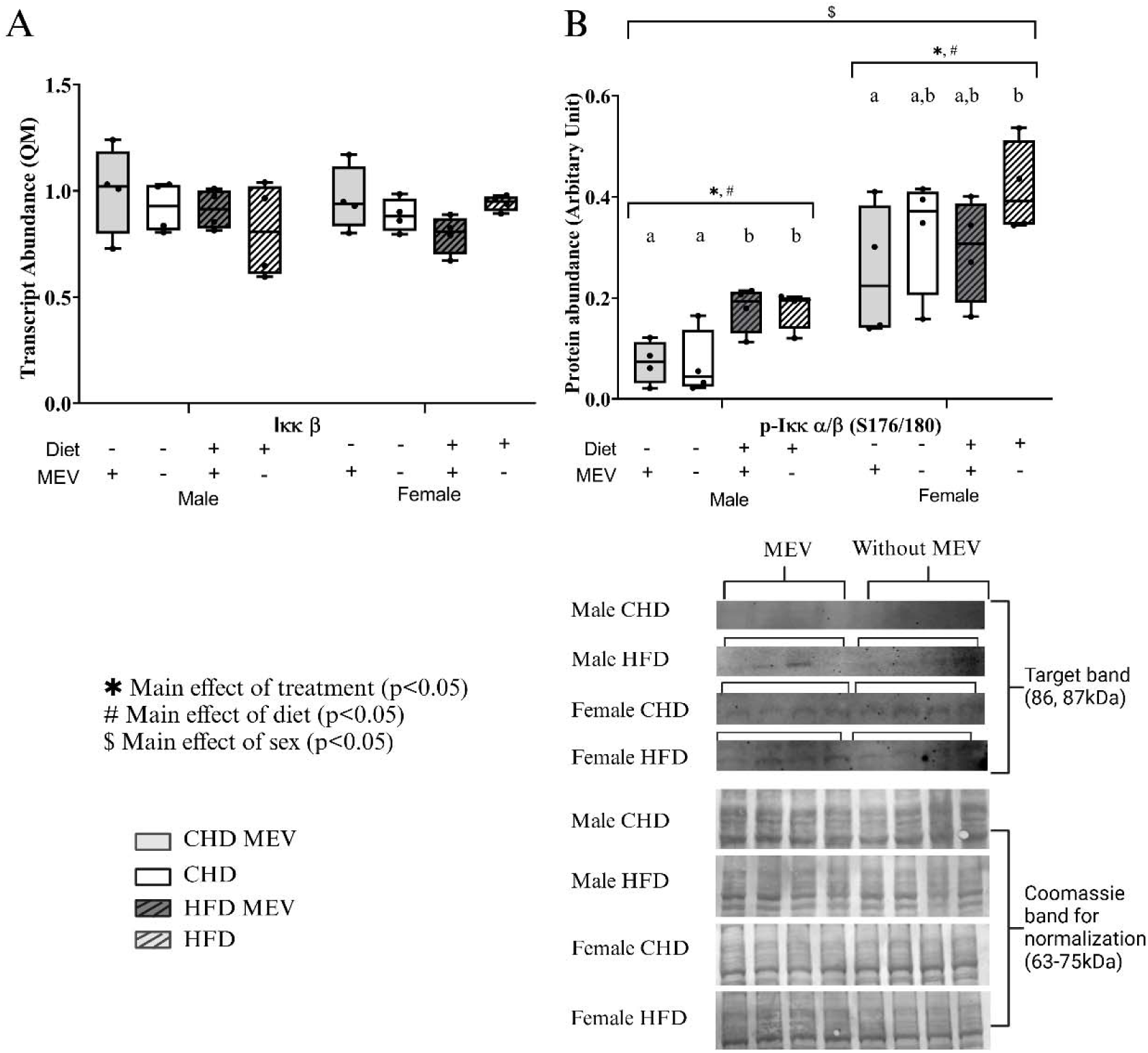
Transcript abundance (Quantity Means) and protein abundance (normalized to Coomassie stain) of NF-κB pathway markers as determined by RT-qPCR and western immunoblotting, respectively in the liver of male and female neonates at postnatal day (PND) 11. (**A**) *I*κκ*-*β transcript (**B**) phosphorylate-Iκκ-α/β protein (S176/180). CHD MEV = control offspring that received milk-derived extracellular vesicle (MEV) treatment from PND 4-11. CHD = control offspring that did not receive MEV treatment. HFD MEV = offspring with perinatal high fat diet exposure and received MEV treatment from PND 4-11. HFD = offspring with perinatal high fat diet exposure that did not receive MEV treatment. MEV dosage: 1.44×10^10^ particles per grams of body weight. Data are presented with standard error of means. N = 4 biological replicates/ diet/treatment/sex. * Main effect of MEV treatment. # Main effect of diet. $ Main effect of sex. Same letters above the boxplots represents no changes and different letters represent significant difference as determined by Tukey HSD. Effects are statistically significant at p<0.05.

*I*κ*B*α transcript abundance illustrated a main effect of sex (*F* _(1, 24)_ = 7.638, p = 0.011). A main effect of diet (*F* _(1, 12)_ = 9.368, p = 0.010) and a main effect of MEV treatment (*F* _(3, 12)_ = 4.023, p = 0.034) were seen in males (**Figure 5A**). In females, a main effect of diet (*F* _(1, 12)_ = 26.548, p<0.001) and MEV treatment (*F* _(3, 12)_ = 9.639, p = 0.002) was seen, where *I*κ*B*α transcript levels were lower in HFD females and HFD-MEV females, compared to CHD and CHD-MEV females (p=0.003). P-IκBα (S32) level (**Figure 5B**) did not change in response to sex (*U* = 92.00, p = 0.184), diet (*F* _(1, 12)_ = 0.117, p = 0.738 in male samples) (*F* _(1, 12)_ = 0.171, p = 0.687 in female samples), or MEV treatment (*F* _(3, 12)_ = 0.197, p = 0.896 in male samples) (*F* _(3, 12)_ = 0.987, p = 0.432 in female samples).

**Figure 5:**
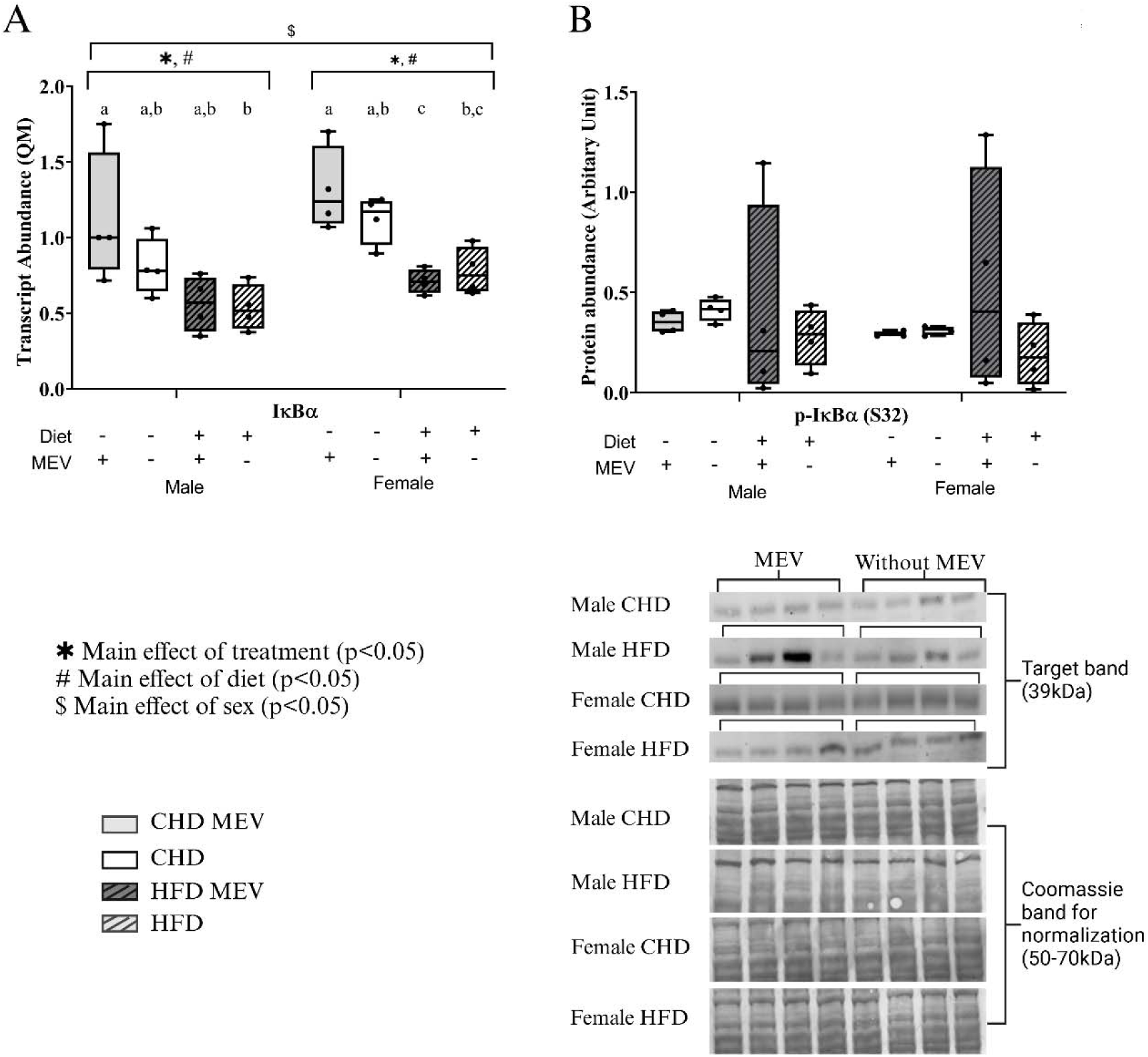
Transcript abundance (Quantity Means) and protein abundance (normalized to Coomassie stain) of NF-κB pathway markers as determined by RT-qPCR and western immunoblotting, respectively in the liver of male and female neonates at postnatal day (PND) 11. (**A**) *I*κ*B*α transcript (**B**) IκBα (S32) protein. CHD MEV = control offspring that received milk-derived extracellular vesicle (MEV) treatment from PND 4-11. CHD = control offspring that did not receive MEV treatment. HFD MEV = offspring with perinatal high fat diet exposure and received MEV treatment from PND 4-11. HFD = offspring with perinatal high fat diet exposure that did not receive MEV treatment. MEV dosage: 1.44×10^10^ particles per grams of body weight. Data are presented with standard error of means. N = 4 biological replicates/ diet/treatment/sex. * Main effect of MEV treatment. # Main effect of diet. $ Main effect of sex. Same letters above the boxplots represents no changes and different letters represent significant difference as determined by Tukey HSD. Effects are statistically significant at p<0.05.

Main effect of sex (*F* _(1, 24)_ = 6.892, p = 0.015), also main effect of diet (*F* _(1, 12)_ = 17.807, p<0.001) and MEV treatment (*F* _(3, 12)_ = 6.960, p = 0.006) were seen in *NF-*κ*B* transcript abundance in males (**Figure 6A**), where *NF-*κ*B* was lower in HFD (p = 0.006) and HFD-MEV (p = 0.025) compared to CHD. *NF-*κ*B* transcript abundance in females remained unchanged in response to diet (*F* _(1, 12)_ = 0.022, p = 0.885) or MEV treatment (*F* _(3, 12)_ = 0.672, p = 0.586). NF-κB total protein levels (**Figure 6B**) remained unchanged between sex (*F* _(1, 24)_ = 1.333, p = 0.260) and diet (*F* _(1, 12)_ = 0.105, p = 0.752 for male samples) (*F* _(1, 12)_ = 1.007, p = 0.335 for female samples). A main effect of MEV treatment was seen in males (*F* _(3, 12)_ = 6.576, p = 0.007) and females (*F* _(3, 12)_ =11.498, p<0.001). With MEV treatment, HFD-MEV males exhibited increased levels of NF-κB compared to HFD males (p = 0.005). The same treatment effect was seen in HFD females (p<0.001). Protein abundance of p-NF-κB (S536)/NF-κB (**Figure 6C**) on the other hand, had no main effect of diet (*F* _(1, 9)_ = 2.546, p = 0.145) and MEV treatment (*F* _(3, 9)_ = 3.339, p = 0.070) in males. In females, main effect of MEV treatment (*F* _(3, 11)_ = 3.483, p = 0.054) and diet (*F* _(1, 11)_ = 2.978, p = 0.112) were not observed. No main effect of sex was seen in p-NF-κB (S536)/NF-κB (*U* = 28, p = 0.534)

**Figure 6:**
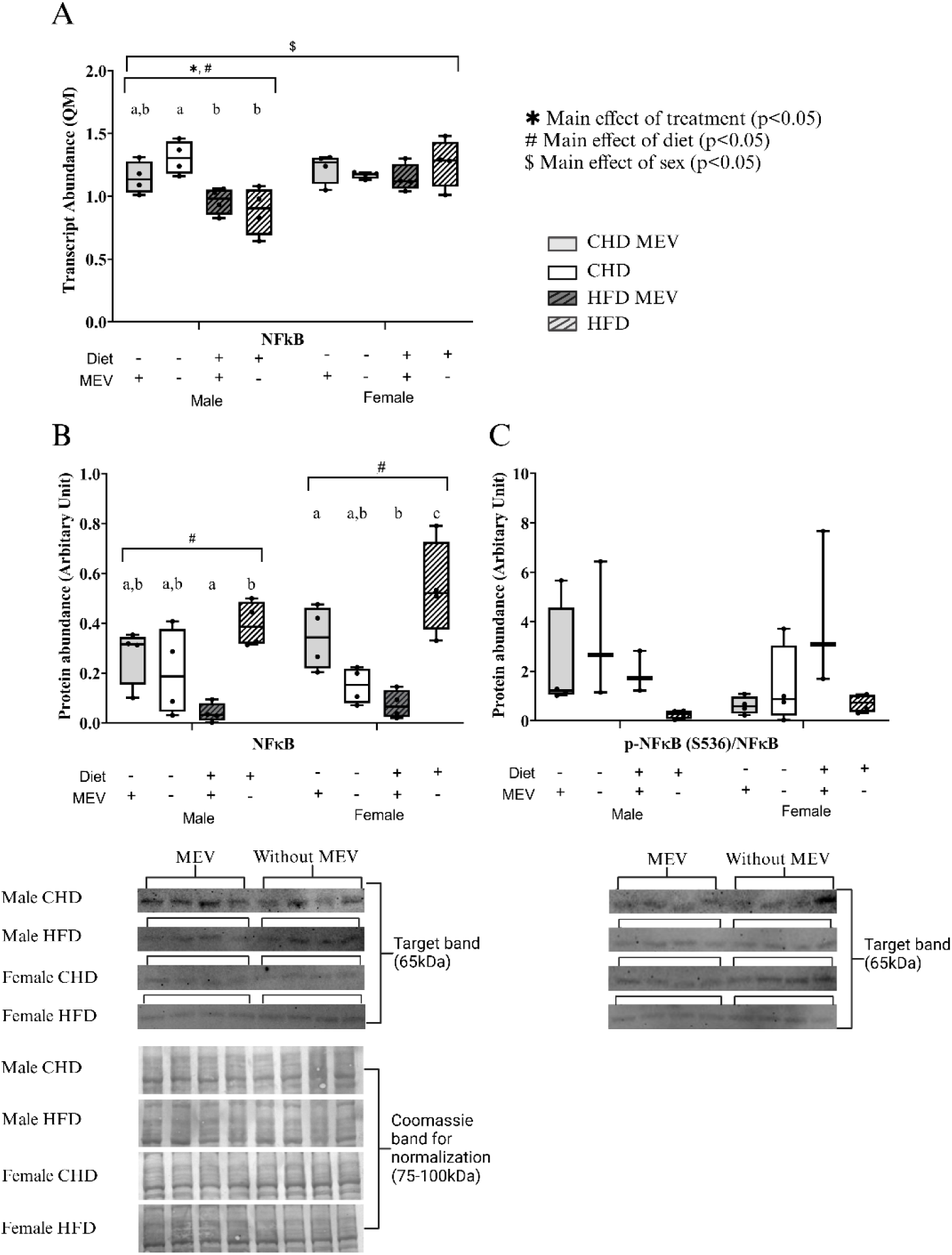
Transcript abundance (Quantity Means) and protein abundance (normalized to Coomassie stain) of NF-κB pathway markers as determined by RT-qPCR and western immunoblotting, respectively in the liver of male and female neonates at postnatal day (PND) 11. (**A**) *NF-*κ*B* transcript (**B**) NF-κB protein (**C**) phosphorylate-NF-κB (S536)/NF-κB protein. CHD MEV = control offspring that received milk-derived extracellular vesicle (MEV) treatment from PND 4-11. CHD = control offspring that did not receive MEV treatment. HFD MEV = offspring with perinatal high fat diet exposure and received MEV treatment from PND 4-11. HFD = offspring with perinatal high fat diet exposure that did not receive MEV treatment. MEV dosage: 1.44×10^10^ particles per grams of body weight. Data are presented with standard error of means. N = 4 biological replicates/ diet/treatment/sex. * Main effect of MEV treatment. # Main effect of diet. $ Main effect of sex. Same letters above the boxplots represents no changes and different letters represent significant difference as determined by Tukey HSD. Effects are statistically significant at p<0.05.

*NLRP*3 transcript abundance in males illustrated a main effect of diet (*F* _(1, 12)_ = 8.492, p=0.013), but not in females (*F* _(1, 12)_ = 0.697, p = 0.420) **(Figure 7A**) No effect of treatment was observed in male (*F* _(3, 12)_ = 2.881, p = 0.080) and in females (*F* _(3, 12)_ = 0.234, p = 0.871) or between sexes (*F* _(1, 24)_ = 0.286, p = 0.598). A main effect of sex (*U* =113, p = 0.590) was not significant in NLRP3 protein abundance (**Figure 7B**). Males exhibited a main effect of diet (*F* _(1, 12)_ = 379.060, p<0.001) and MEV treatment (*F* _(3, 12)_ = 147.067, p<0.001), where HFD males who received MEV treatment had a lower abundance of NLRP3, compared to CHD (p<0.001), CHD-MEV (p<0.001), and HFD (p<0.001). Females illustrated a main effect of diet (*F* _(1, 12)_ = 43.489, p<0.001) and MEV treatment (*F* _(3, 12)_ = 15.173, p<0.001), where NLRP3 protein was higher in HFD females compared to CHD (p<0.004) and CHD-MEV (p<0.001). MEV treatment did not change NLRP3 levels in females.

**Figure 7:**
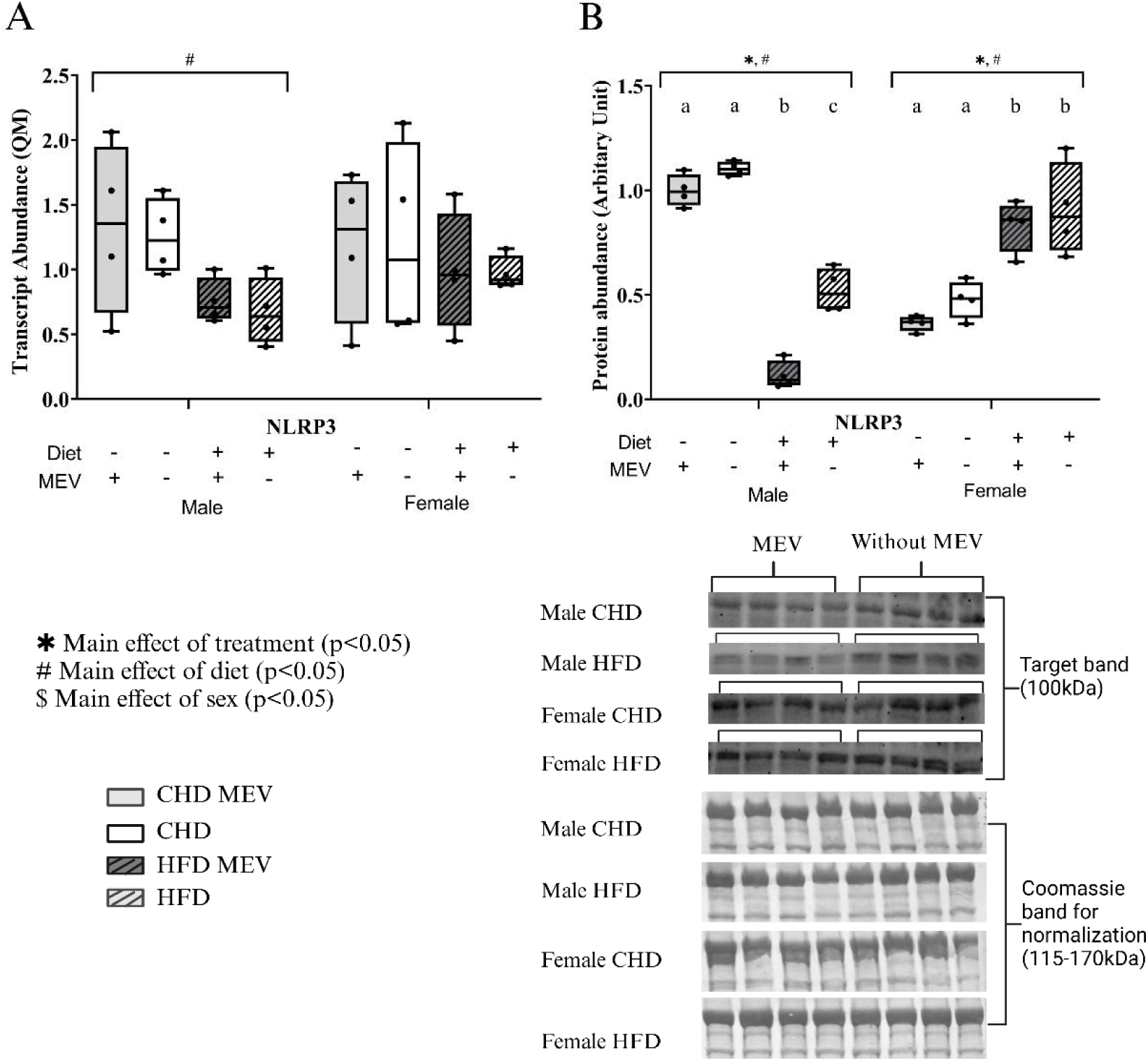
Transcript abundance (Quantity Means) and protein abundance (normalized to Coomassie stain) of NLRP3 inflammasome marker as determined by RT-qPCR and western immunoblotting, respectively in the liver of male and female neonates at postnatal day (PND) 11. (**A**) *NLRP3* transcript (**B**) NLRP3 protein. CHD MEV = control offspring that received milk-derived extracellular vesicle (MEV) treatment from PND 4-11. CHD = control offspring that did not receive MEV treatment. HFD MEV = offspring with perinatal high fat diet exposure and received MEV treatment from PND 4-11. HFD = offspring with perinatal high fat diet exposure that did not receive MEV treatment. MEV dosage: 1.44×10^10^ particles per grams of body weight. Data are presented with standard error of means. N = 4 biological replicates/ diet/treatment/sex. * Main effect of MEV treatment. # Main effect of diet. $ Main effect of sex. Same letters above the boxplots represents no changes and different letters represent significant difference as determined by Tukey HSD. Effects are statistically significant at p<0.05.

*Caspase-1* transcript abundance (**Figure 8A**) remained unchanged in males (diet (*F* _(1, 12)_ = 3.807, p = 0.075) and treatment (*F* _(3, 12)_ = 1.876, p = 0.187)), but females had significant diet (*F* _(1, 12)_ =10.517, p = 0.007) and MEV treatment (*F* _(3, 12)_ = 3.524, p = 0.049) effects. No sex effect was observed (*F* _(1, 24)_ = 0.237, p = 0.631). Pro-caspase-1 protein levels illustrated a main effect of diet in males (*F* _(1, 12)_ = 54.709, p<0.001) and females (*F* _(1, 12)_ = 43.489, p<0.001) (**Figure 8B**). Also, there was a main effect of MEV treatment in males (*F* _(3, 12)_ = 32.059, p<0.001) and females (*F* _(3, 12)_ =15.173, p<0.001). In males, the level of pro-caspase-1 was highest in HFD compared to CHD-MEV (p<0.001) and CHD (p<0.001). Pro-caspase-1 returned to control levels in HFD-MEV males (p<0.001). In females, the level of pro-caspase-1 was reduced in HFD-MEV and HFD compared to CHD-MEV (p = 0.002) and CHD (p = 0.004). Level of cleaved-caspase-1 (**Figure 8C**) remained unchanged in males treatment effect (*F* _(3, 12)_ = 1.212, p = 0.348) and diet effect (*F* _(1, 12)_ = 0.002, p = 0.961) and females treatment effect (*F* _(3, 12)_ = 0.366, p = 0.779) and diet effect (*F* _(1, 12)_ = 0.246, p = 0.629). As well as sex effect between males and females (*F* _(1, 24)_ = 0.026, p = 0.874).

**Figure 8:**
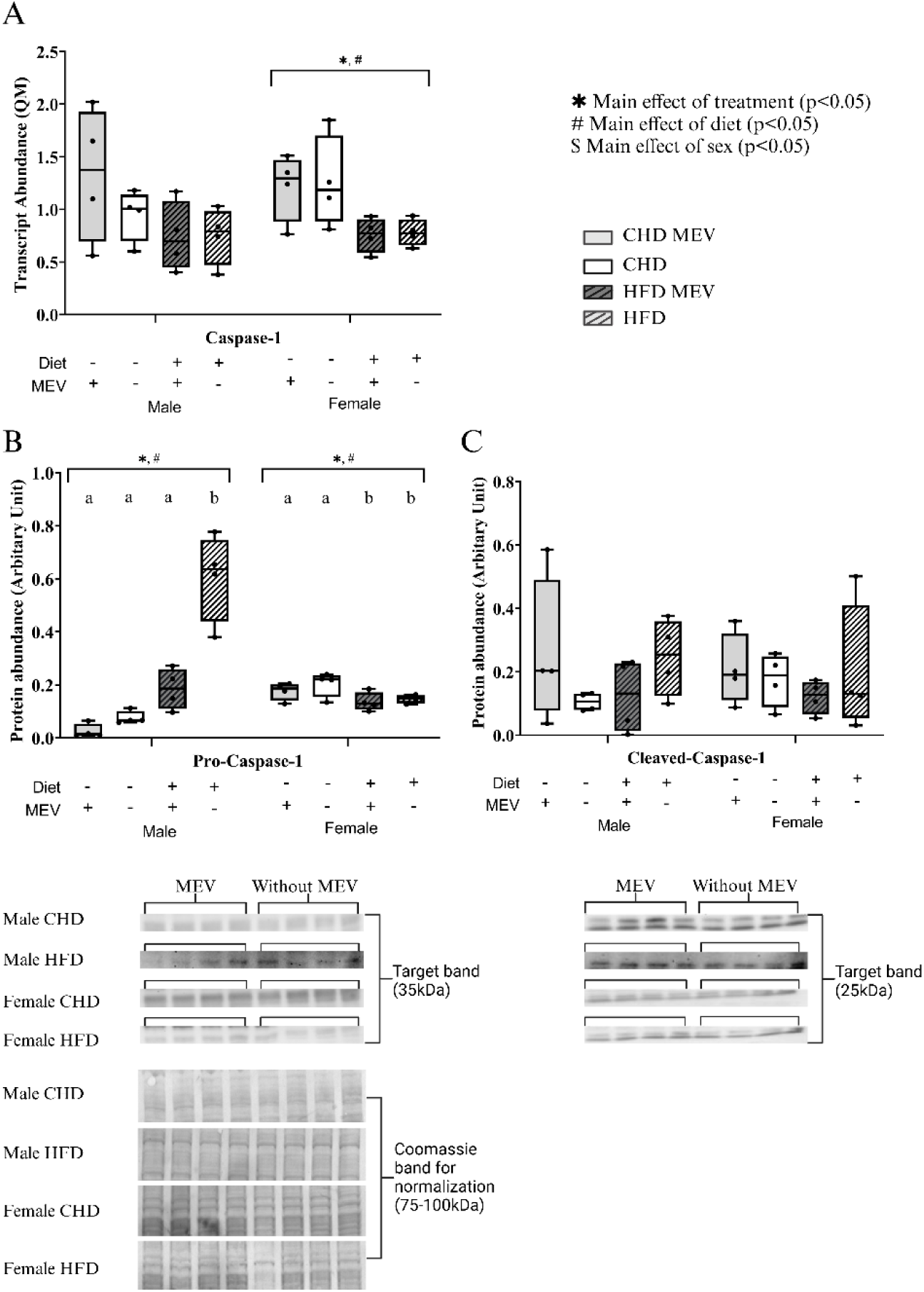
Transcript abundance (Quantity Means) and protein abundance (normalized to Coomassie stain) of NF-κB pathway markers as determined by RT-qPCR and western immunoblotting, respectively in the liver of male and female neonates at postnatal day (PND) 11. (**A**) *Caspase-1* transcript (**B**) pro-caspase-1 protein (**C**) cleaved-caspase-1 protein. CHD MEV = control offspring that received milk-derived extracellular vesicle (MEV) treatment from PND 4-11. CHD = control offspring that did not receive MEV treatment. HFD MEV = offspring with perinatal high fat diet exposure and received MEV treatment from PND 4-11. HFD = offspring with perinatal high fat diet exposure that did not receive MEV treatment. MEV dosage: 1.44×10^10^ particles per grams of body weight. Data are presented with standard error of means. N = 4 biological replicates/ diet/treatment/sex. * Main effect of MEV treatment. # Main effect of diet. $ Main effect of sex. Same letters above the boxplots represents no changes and different letters represent significant difference as determined by Tukey HSD. Effects are statistically significant at p<0.05.

The transcript abundance of IL-18, (**Figure 9**) remained unchanged in response to diet (*F* _(1, 12)_ = 2.148, p = 0.168) and treatment (*F* _(3, 12)_ = 2.453, p = 0.113) in males and females (*F* _(1, 12)_ = 1.770, p = 0.1210) (*F* _(3, 12)_ = 0.756, p = 0.542). Also, no sex effect was observed (*F* _(1, 24)_ = 0.741, p = 0.398).

**Figure 9:**
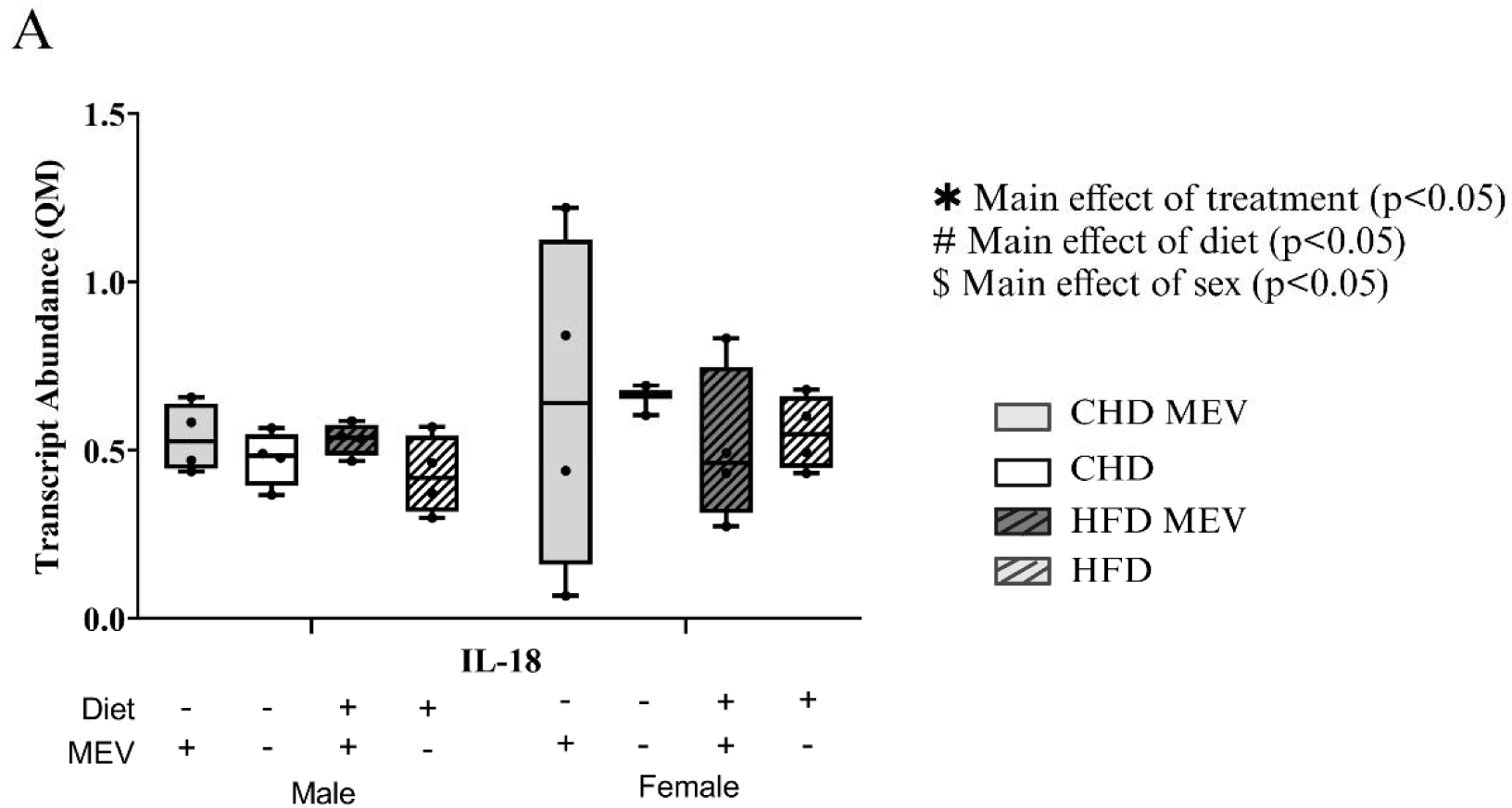
Transcript abundance (Quantity Means) and protein abundance (normalized to Coomassie stain) of NLRP3 inflammasome marker as determined by RT-qPCR and western immunoblotting, respectively in the liver of male and female neonates at postnatal day (PND) 11. (**A**) *IL-18* transcript. CHD MEV = control offspring that received milk-derived extracellular vesicle (MEV) treatment from PND 4-11. CHD = control offspring that did not receive MEV treatment. HFD MEV = offspring with perinatal high fat diet exposure and received MEV treatment from PND 4-11. HFD = offspring with perinatal high fat diet exposure that did not receive MEV treatment. MEV dosage: 1.44×10^10^ particles per grams of body weight. Data are presented with standard error of means. N = 4 biological replicates/ diet/treatment/sex. * Main effect of MEV treatment. # Main effect of diet. $ Main effect of sex. Same letters above the boxplots represents no changes and different letters represent significant difference as determined by Tukey HSD. Effects are statistically significant at p<0.05.

### 3.4.5 Transcript and protein abundance of NF-kb pathway targets in neonatal hypothalamus in response to diet and MEV treatment

Transcript and protein abundance of TLR4, IKKβ, IκBα, NF-κB, NLRP3, caspase-1, and IL-18) in male and female hypothalamus were measured. *TLR4* transcript had a main effect of sex (**Figure 10A**) (*F* _(1, 22)_ = 5.244, p = 0.032), but remained unchanged in response to diet (*F* _(1, 12)_ = 2.782, p=0.121 in males) (*F* _(1, 10)_ = 0.000, p = 0.996 in females) and treatment (*F* _(3, 12)_ = 1.783, p = 0.204 in males) (*F* _(3, 10)_ = 1.561, p = 0.259 in females). In TLR4 protein level, a main effect of diet (**Figure 10B**) in males (*F* _(1, 10)_ = 60.896, p<0.001) and females (*F* _(1, 10)_ = 36.027, p<0.001) and a main effect of treatment in males (*F* _(3, 10)_ = 21.040, p<0.001) and females (*F* _(3, 10)_ = 12.755, p<0.001) was present. In males TLR4 decreased in HFD-MEV (p<0.001) and HFD (p = 0.002) compared to CHD. Same trend was seen in females, where TLR4 was lower in HFD-MEV (p = 0.002) and HFD (p = 0.014). Main effect of sex was not observed in TLR4 protein abundance (*F* _(1, 20)_ = 0.603, p = 0.447).

**Figure 10:**
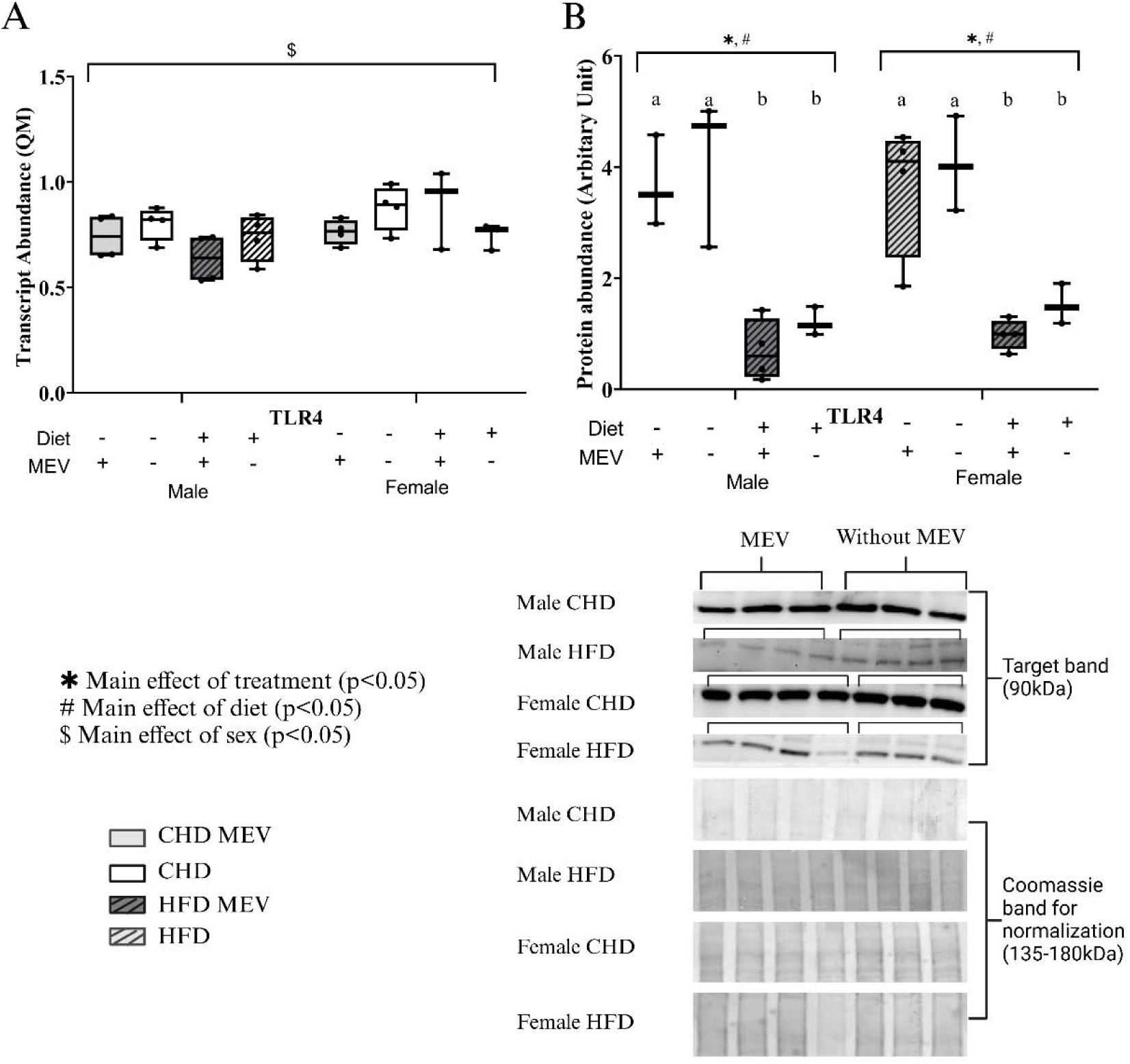
Transcript abundance (Quantity Means) and protein abundance (normalized to Coomassie stain) of NF-κB pathway markers as determined by RT-qPCR and western immunoblotting, respectively in the hypothalamus of male and female neonates at postnatal day (PND) 11. (**A**) *TLR4* transcript (**B**) TLR4 protein. CHD MEV = control offspring that received milk-derived extracellular vesicle (MEV) treatment from PND 4-11. CHD = control offspring that did not receive MEV treatment. HFD MEV = offspring with perinatal high fat diet exposure and received MEV treatment from PND 4-11. HFD = offspring with perinatal high fat diet exposure that did not receive MEV treatment. MEV dosage: 1.44×10^10^ particles per grams of body weight. Data are presented with standard error of means. N = 4 biological replicates/ diet/treatment/sex. * Main effect of MEV treatment. # Main effect of diet. $ Main effect of sex. Same letters above the boxplots represents no changes and different letters represent significant difference as determined by Tukey HSD. Effects are statistically significant at p<0.05.

There was a main effect of sex (*F* _(1, 24)_ =16.424, p<0.001) in *I*κκ*-*β transcript abundance. No changes were seen in males between treatment groups (*F* _(3, 12)_ = 0.215, p = 0.884) and diet (*F* _(1, 12)_ = 0.070, p = 0.796). However, in females, there was a main effect of treatment (*F* _(3, 12)_ = 9.675, p = 0.002) and diet (*F* _(1, 12)_ = 24.437, p<0.001) (**Figure 11A**), where *I*κκ*-*β level increased in HFD (p = 0.024) and HFD-MEV (p = 0.008) compared to CHD and CHD-MEV. There was also a main effect of sex in p-Iκκ-α/β (S176/180) (*U* = 36, p = 0.004), where females had lower protein abundance than males (**Figure 11B**). No changes were seen in males between treatment groups (*F* _(3, 10)_ = 1.959, p = 0.184) and diet groups (*F* _(1, 10)_ = 3.564, p = 0.088). In females, there was a main effect of treatment (*F* _(3, 10)_ = 8.905, p = 0.004) and diet (*F* _(1, 10)_ = 24.200, p<0.001), where levels of p-IKK-α/β (S176/180) were higher in HFD-MEV (p = 0.006) and HFD (p = 0.019) than CHD-MEV samples.

**Figure 11:**
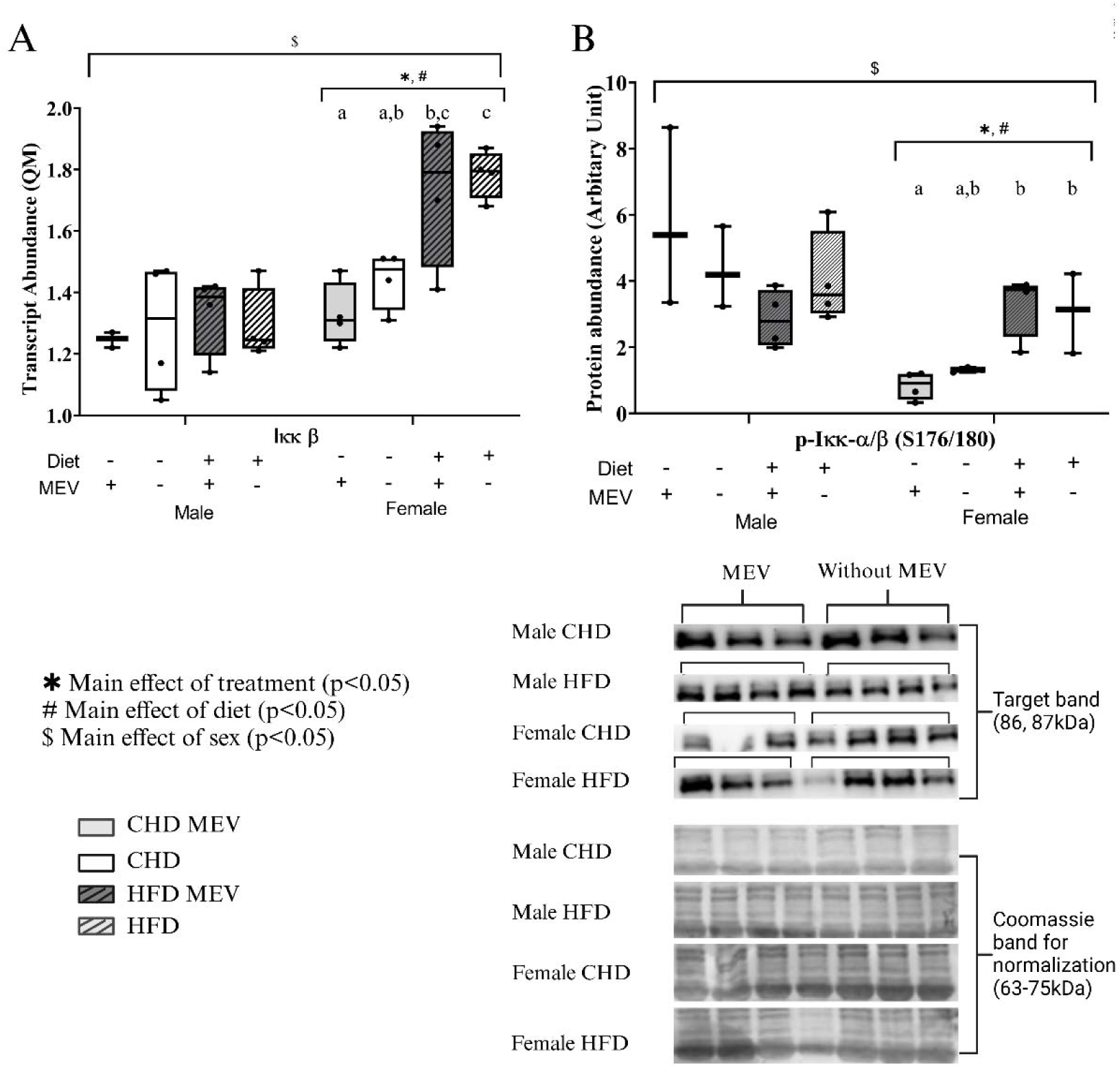
Transcript abundance (Quantity Means) and protein abundance (normalized to Coomassie stain) of NF-κB pathway markers as determined by RT-qPCR and western immunoblotting, respectively in the hypothalamus of male and female neonates at postnatal day (PND) 11. (**A**) *I*κκ*-*β transcript (**B**) phosphorylate-Iκκ-α/β protein (S176/180). CHD MEV = control offspring that received milk-derived extracellular vesicle (MEV) treatment from PND 4-11. CHD = control offspring that did not receive MEV treatment. HFD MEV = offspring with perinatal high fat diet exposure and received MEV treatment from PND 4-11. HFD = offspring with perinatal high fat diet exposure that did not receive MEV treatment. MEV dosage: 1.44×10^10^ particles per grams of body weight. Data are presented with standard error of means. N = 4 biological replicates/ diet/treatment/sex. * Main effect of MEV treatment. # Main effect of diet. $ Main effect of sex. Same letters above the boxplots represents no changes and different letters represent significant difference as determined by Tukey HSD. Effects are statistically significant at p<0.05.

A main effect of diet (*F* _(1, 12)_ = 17.800, p = 0.001) and treatment (*F* _(3, 12)_ = 8.989, p = 0.002) was seen in *I*κ*B*α transcript level in males (**Figure 12A**), where *I*κ*B*α was higher in HFD-MEV compared to CHD (p = 0.002) and CHD-MEV (p = 0.006) but remained unchanged with MEV treatment (p = 0.921). In females, there was a main effect of treatment (*F* _(3, 12)_ = 3.913, p = 0.037), and there was no main effect of diet (*F* _(1, 12)_ = 4.375, p = 0.058). There was no main effect of sex at the transcript level (*F* _(1, 24)_ = 0.745, p = 0.397). Nevertheless, there was a main effect of sex (*U* =143, p = 0.039) in p-IκBα (S32) (**Figure 12B**), where HFD females had more p-IκBα. MEV treatment led to changes in male and female p-IκBα (S32). A main effect of diet was seen in males (*U* =7, p = 0.029) and females (*U* = 49, p = 0.001). In females, there was also a main effect of MEV treatment (*H* _(3)_ = 10.667, p = 0.014), where p-IκBα (S32) was higher in HFD-MEV (p = 0.043) and HFD (p = 0.015) compared to CHD-MEV and CHD. Main effect of treatment was not seen in males (*H* _(3)_ = 6.062, p = 0.109).

**Figure 12:**
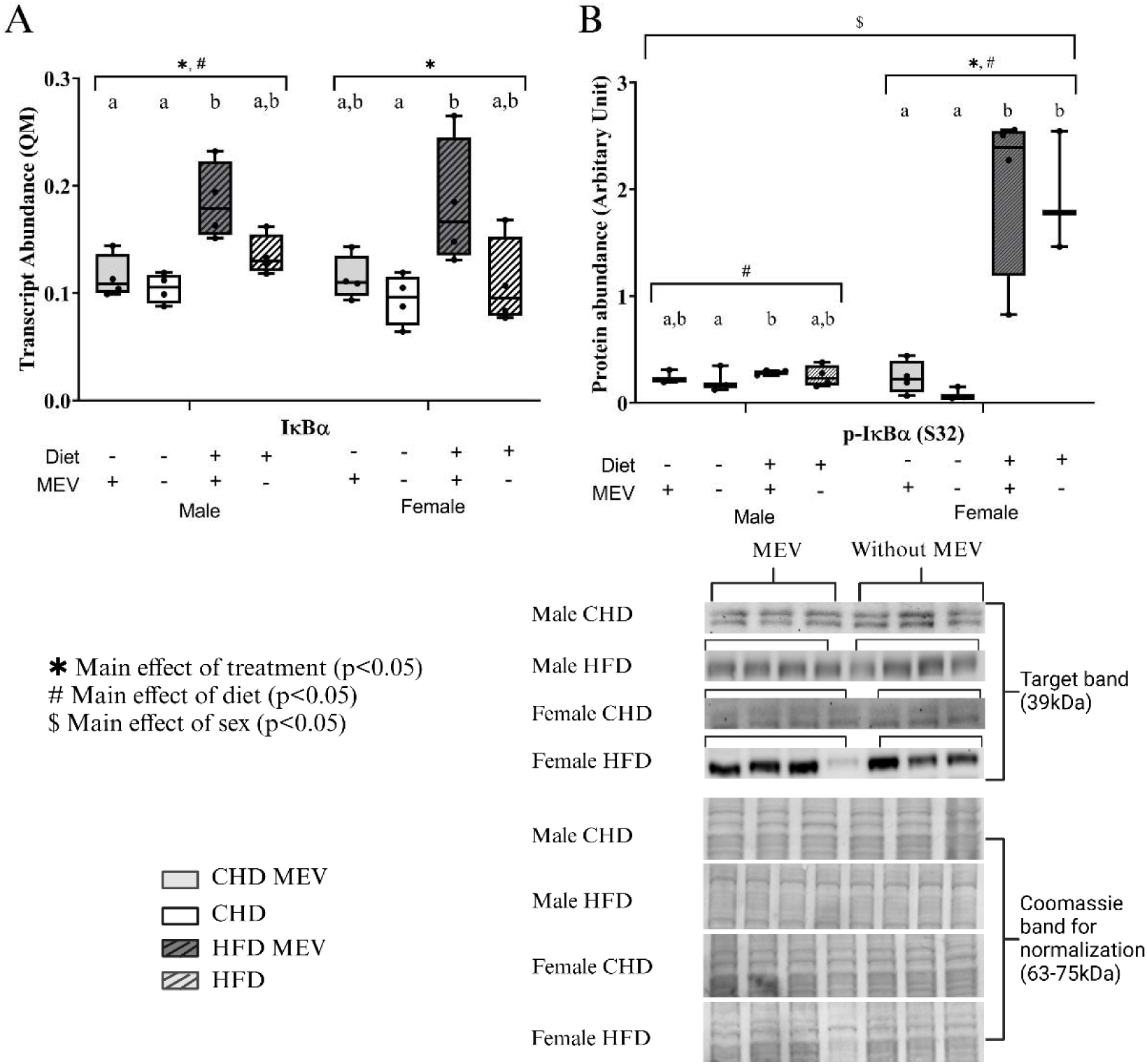
Transcript abundance (Quantity Means) and protein abundance (normalized to Coomassie stain) of NF-κB pathway markers as determined by RT-qPCR and western immunoblotting, respectively in the hypothalamus of male and female neonates at postnatal day (PND) 11. (**A**) *I*κ*B*α transcript (**B**) IκBα (S32) protein. CHD MEV = control offspring that received milk-derived extracellular vesicle (MEV) treatment from PND 4-11. CHD = control offspring that did not receive MEV treatment. HFD MEV = offspring with perinatal high fat diet exposure and received MEV treatment from PND 4-11. HFD = offspring with perinatal high fat diet exposure that did not receive MEV treatment. MEV dosage: 1.44×10^10^ particles per grams of body weight. Data are presented with standard error of means. N = 4 biological replicates/ diet/treatment/sex. * Main effect of MEV treatment. # Main effect of diet. $ Main effect of sex. Same letters above the boxplots represents no changes and different letters represent significant difference as determined by Tukey HSD. Effects are statistically significant at p<0.05.

*NF-*κ*B* transcript abundance (**Figure 13A)** had a main effect of diet (*F* _(1, 12)_ = 12.351, p = 0.004) and treatment (*F* _(3, 12)_ = 5.025, p = 0.017) in males, where HFD-MEV males had higher levels of *NF-*κ*B* than CHD (p = 0.020) and CHD-MEV males (p = 0.044). Same trend was seen in females, where both diet (*F* _(1, 12)_ = 25.354, p<0.001) and treatment effect (*F* _(3, 12)_ = 9.828, p = 0.001) were seen and the level of *NF-*κ*B* is higher in HFD-MEV than CHD (p = 0.003) and CHD-MEV (p = 0.003). No main effect of sex was seen between males and females in *NF-*κ*B* transcript abundance (*F* _(1, 24)_ = 0.617, p = 0.440). Total NF-κB protein (**Figure 13B**) remained unchanged between diet and treatment groups in males (*F* _(1, 10)_ = 0.016, p = 0.902) (*F* _(3, 10)_ = 2.204, p = 0.151) and females (*F* _(1, 10)_ = 4.930, p = 0.051) (*F* _(3, 10)_ = 3.182, p = 0.572), although there was a main effect of sex (*U* = 35, p = 0.003). P-NF-κB (S536)/NF-κB (**Figure 13C**) showed the main effect of diet (*F* _(1, 10)_ = 6.905, p = 0.025) and MEV treatment (*F* _(3, 10)_ = 21.350, p<0.001) in males, where HFD-MEV had a higher level of p-NF-κB (S536)/NF-κB than HFD (P<0.001). However, in females no effect of diet (*F* _(1, 9)_ = 4.015, p = 0.076) and treatment was seen (*F* _(3, 9)_ = 2.360, p = 0.139). Sex effect was not seen in p-NF-κB (S536)/NF-κB (*U* = 81, p = 0.650).

**Figure 13:**
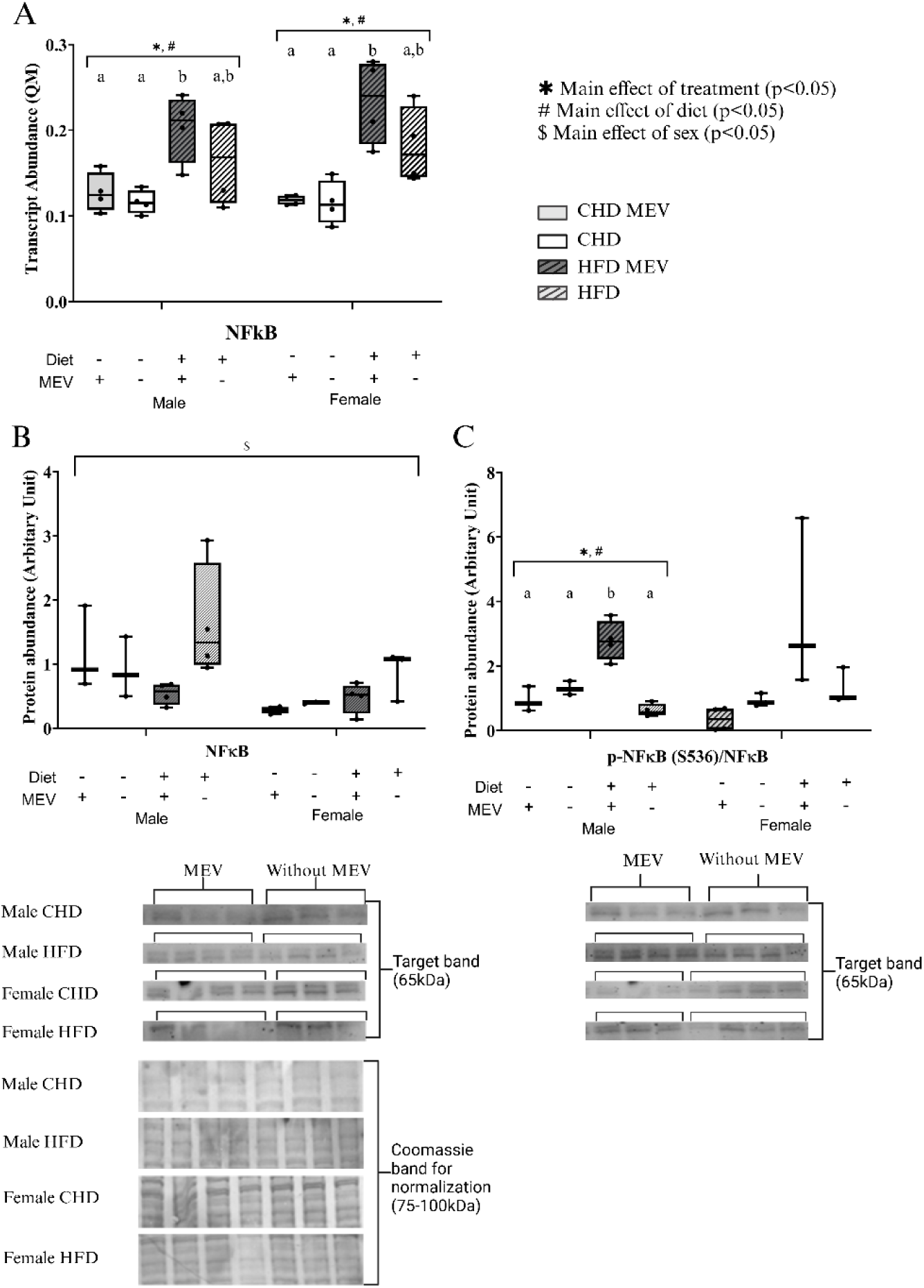
Transcript abundance (Quantity Means) and protein abundance (normalized to Coomassie stain) of NF-κB pathway markers as determined by RT-qPCR and western immunoblotting, respectively in the hypothalamus of male and female neonates at postnatal day (PND) 11. (**A**) *NF-*κ*B* transcript (**B**) NF-κB protein (**C**) phosphorylate-NF-κB (S536)/NF-κB protein. CHD MEV = control offspring that received milk-derived extracellular vesicle (MEV) treatment from PND 4-11. CHD = control offspring that did not receive MEV treatment. HFD MEV = offspring with perinatal high fat diet exposure and received MEV treatment from PND 4-11. HFD = offspring with perinatal high fat diet exposure that did not receive MEV treatment. MEV dosage: 1.44×10^10^ particles per grams of body weight. Data are presented with standard error of means. N = 4 biological replicates/ diet/treatment/sex. * Main effect of MEV treatment. # Main effect of diet. $ Main effect of sex. Same letters above the boxplots represents no changes and different letters represent significant difference as determined by Tukey HSD. Effects are statistically significant at p<0.05.

*NLRP3* transcript abundance illustrated a main effect of sex, where females had higher abundance than males (*F* _(1, 24)_ = 8.861, p = 0.007) (**Figure 14A**). In females, there was a main effect of diet (*F* _(1, 12)_ = 5.037, p = 0.044), and no changes were seen in female treatment (*F* _(3, 10)_ = 2.436, p = 0.115) and males for diet (*F* _(1, 12)_ = 0.002, p = 0.970) or treatment (*F* _(3, 12)_ = 0.881, p = 0.479). In males, NLRP3 protein (**Figure 14B**) levels were influenced by diet (*F* _(1, 10)_ = 7.918, p = 0.018) and treatment (*F* _(3, 10)_ = 10.883, p = 0.002), where NLRP3 was highest in HFD males compared to CHD-MEV (p = 002) and CHD (p = 0.040). Interestingly, NLRP3 protein levels returned to control levels in HFD males who received MEVs (p = 0.004). In females, there was a main effect of treatment (*F* _(3, 10)_ = 5.153, p = 0.021), where NLRP3 protein decreased in abundance in HFD-MEV compared to HFD (p = 0.023), but no main effect of diet (*F* _(1, 10)_ = 1.229, p = 0.293). There was also no main effect of sex in NLRP3 protein abundance (*F* _(1, 20)_ = 2.932, p = 0.102).

**Figure 14:**
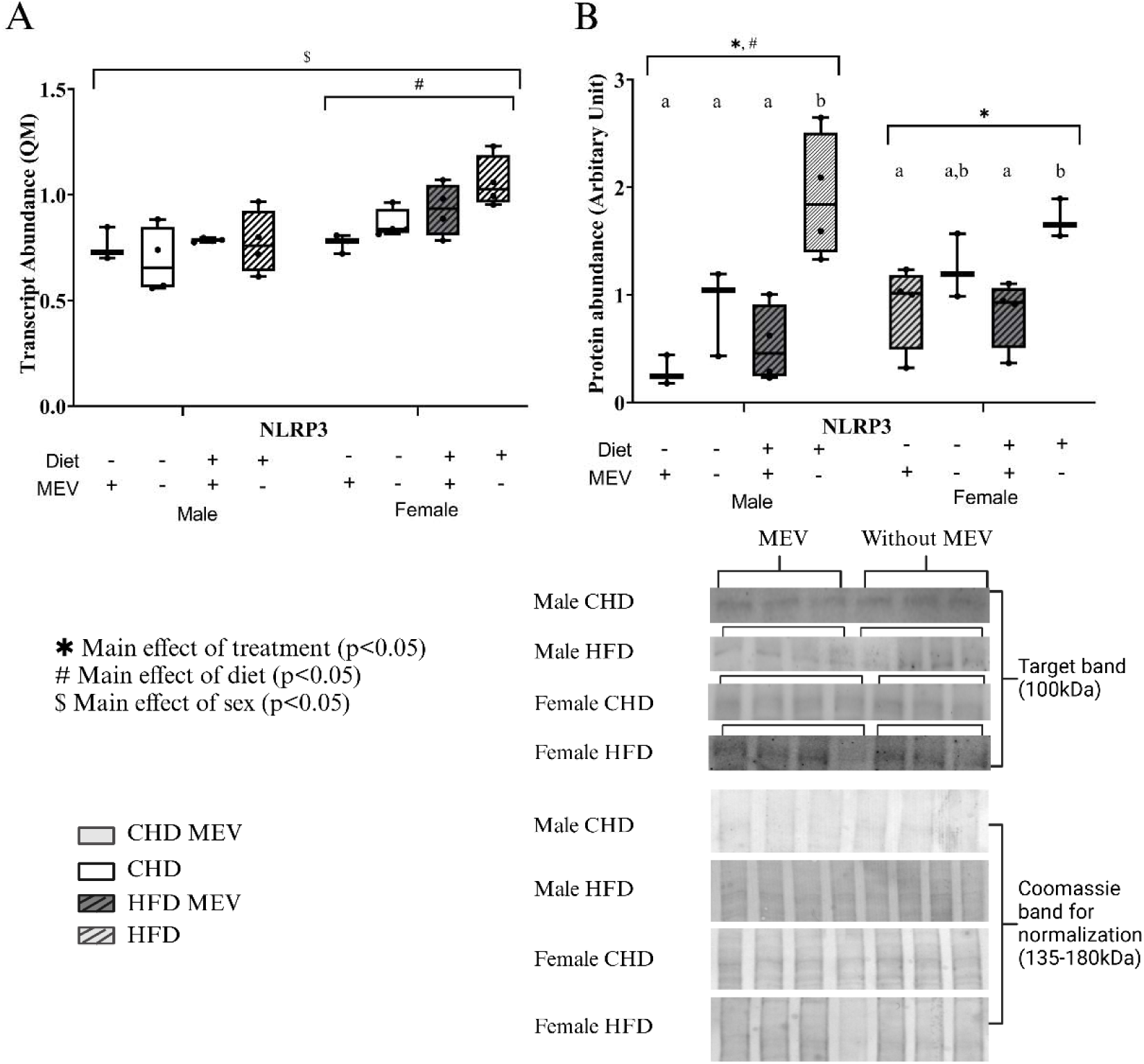
Transcript abundance (Quantity Means) and protein abundance (normalized to Coomassie stain) of NLRP3 inflammasome marker as determined by RT-qPCR and western immunoblotting, respectively in the hypothalamus of male and female neonates at postnatal day (PND) 11. (**A**) *NLRP3* transcript (**B**) NLRP3 protein. CHD MEV = control offspring that received milk-derived extracellular vesicle (MEV) treatment from PND 4-11. CHD = control offspring that did not receive MEV treatment. HFD MEV = offspring with perinatal high fat diet exposure and received MEV treatment from PND 4-11. HFD = offspring with perinatal high fat diet exposure that did not receive MEV treatment. MEV dosage: 1.44×10^10^ particles per grams of body weight. Data are presented with standard error of means. N = 4 biological replicates/ diet/treatment/sex. * Main effect of MEV treatment. # Main effect of diet. $ Main effect of sex. Same letters above the boxplots represents no changes and different letters represent significant difference as determined by Tukey HSD. Effects are statistically significant at p<0.05.

*Caspase-1* (**Figure 15A**) transcript abundance remained unchanged in diet and treatment males (*F* _(1, 12)_ = 0.700, p = 0.419) (*F* _(3, 12)_ = 1.216, p = 0.346) and females (*F* _(1, 12)_ = 0.680, p = 0.426) (*F* _(3, 12)_ = 0.350, p = 0.790), also no main effect of sex (*F* _(1, 24)_ = 1.586, p = 0.220). Pro-caspase-1 protein levels (**Figure 15B**) illustrated a main effect of sex (*U* =191, p<0.001), where females had higher protein abundance than males. There was a treatment effect (*H* _(3)_ = 9.167, p = 0.027) and a diet effect (*U* = 7, p = 0.029) in males, where in HFD-MEV males, pro-caspase-1 decreased in abundance compared to HFD (p = 0.035) and CHD (p = 0.004). In females, pro-caspase-1 remained unchanged with diet (*F* _(1, 10)_ = 2.874, p = 0.121) and treatment (*F* _(3, 10)_ = 2.144, p = 0.158). There was a main effect of sex in cleaved-caspase-1 protein abundance (**Figure 15C**) (*F* _(1, 20)_ = 11.899, p = 0.003). In males, a main effect of diet (*F* _(1, 10)_ = 57.851, p<0.001) and treatment (*F* _(3, 10)_ = 19.919, p<0.001) was present, where HFD-MEV and HFD males had higher cleaved-caspase I compared to CHD-MEV (p = 0.004) and CHD males (p<0.001). In females a main effect of treatment (*F* _(3, 10)_ = 7.548, p = 0.006) was observed but no main effect of diet (*F* _(1, 10)_ = 0.040, p = 0.845). Cleaved-caspase-1 was higher in HFD-MEV than HFD (p = 0.007), and CHD-MEV (p = 0.027).

**Figure 15:**
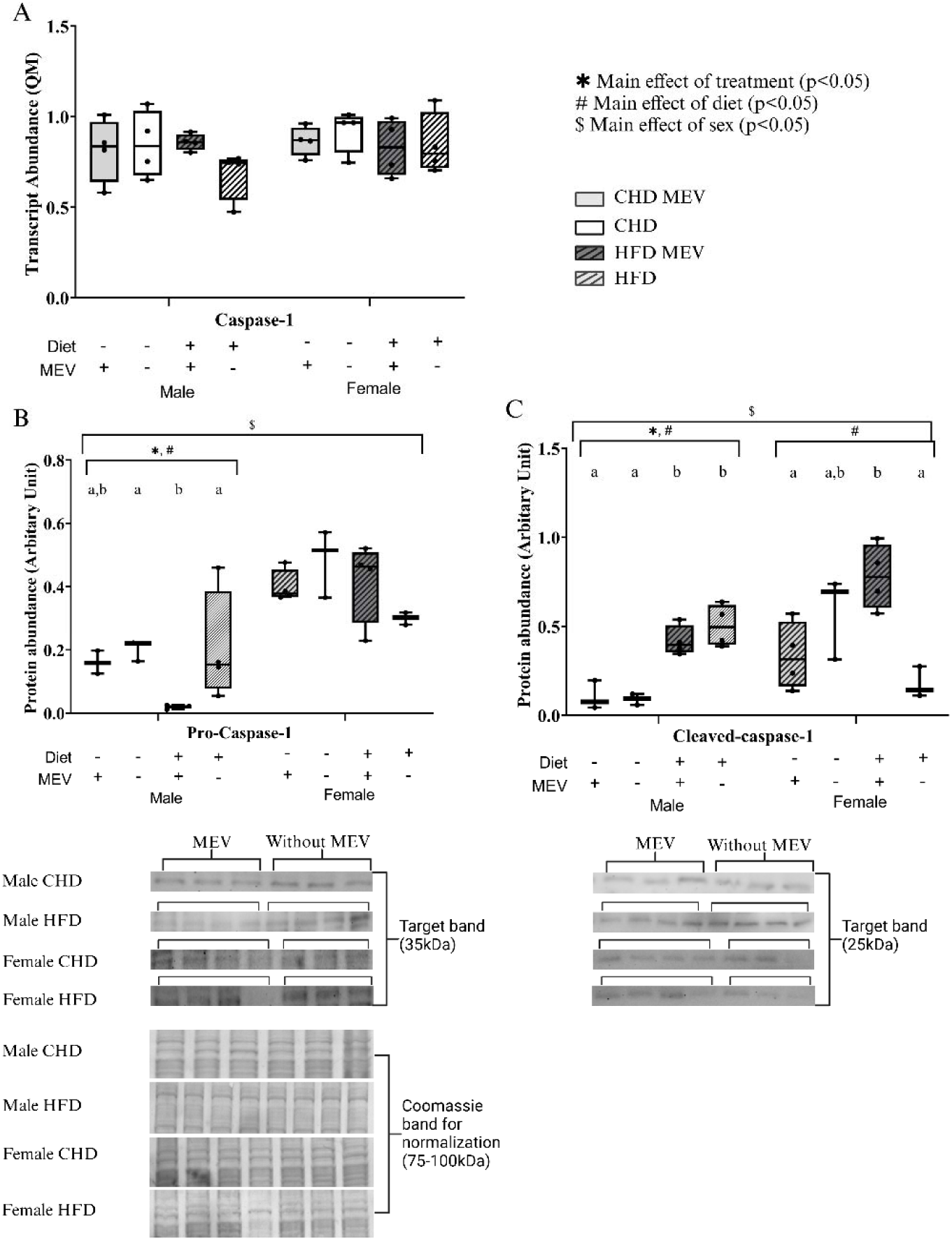
Transcript abundance (Quantity Means) and protein abundance (normalized to Coomassie stain) of NF-κB pathway markers as determined by RT-qPCR and western immunoblotting, respectively in the hypothalamus of male and female neonates at postnatal day (PND) 11. (**A**) *Caspase-1* transcript (**B**) pro-caspase-1 protein (**C**) cleaved-caspase-1 protein. CHD MEV = control offspring that received milk-derived extracellular vesicle (MEV) treatment from PND 4-11. CHD = control offspring that did not receive MEV treatment. HFD MEV = offspring with perinatal high fat diet exposure and received MEV treatment from PND 4-11. HFD = offspring with perinatal high fat diet exposure that did not receive MEV treatment. MEV dosage: 1.44×10^10^ particles per grams of body weight. Data are presented with standard error of means. N = 4 biological replicates/ diet/treatment/sex. * Main effect of MEV treatment. # Main effect of diet. $ Main effect of sex. Same letters above the boxplots represents no changes and different letters represent significant difference as determined by Tukey HSD. Effects are statistically significant at p<0.05.

A main effect of sex was seen for *IL-18* (**Figure 16**) transcript (*F* _(1, 24)_ = 9.902, p = 0.004), where HFD females had higher levels of *IL-18*. In males, there was a diet (*F* _(1, 12)_ =4.845, p = 0.048) and treatment effect (*F* _(3, 12)_ = 7.737, p = 0.004). *IL-18* transcript levels returned to CHD levels with MEV treatment in HFD males (p = 0.015). In females, there was a main effect of diet (*F* _(1, 12)_ = 24.919, p<0.001) and treatment (*F* _(3, 12)_ = 8.544, p=0.003), where *IL-18* transcript abundance was higher in HFD-MEV and HFD females compared to CHD-MEV (p = 0.038) and CHD females (p = 0.009).

**Figure 16:**
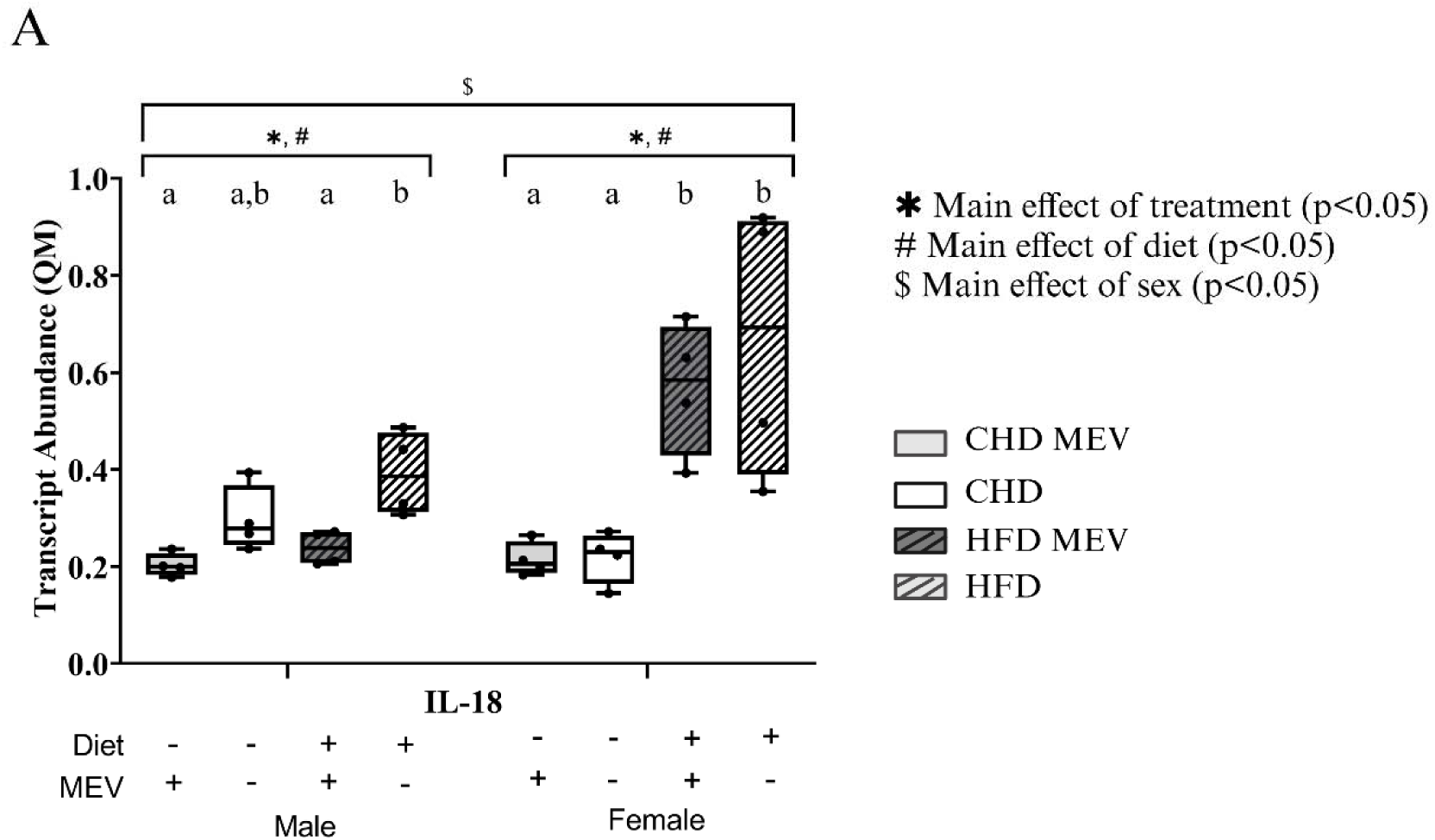
Transcript abundance (Quantity Means) and protein abundance (normalized to Coomassie stain) of NLRP3 inflammasome marker as determined by RT-qPCR and western immunoblotting, respectively in the hypothalamus of male and female neonates at postnatal day (PND) 11. (**A**) *IL-18* transcript. CHD MEV = control offspring that received milk-derived extracellular vesicle (MEV) treatment from PND 4-11. CHD = control offspring that did not receive MEV treatment. HFD MEV = offspring with perinatal high fat diet exposure and received MEV treatment from PND 4-11. HFD = offspring with perinatal high fat diet exposure that did not receive MEV treatment. MEV dosage: 1.44×10^10^ particles per grams of body weight. Data are presented with standard error of means. N = 4 biological replicates/ diet/treatment/sex. * Main effect of MEV treatment. # Main effect of diet. $ Main effect of sex. Same letters above the boxplots represents no changes and different letters represent significant difference as determined by Tukey HSD. Effects are statistically significant at p<0.05.

Transcript and protein data for male and female neonates who received PBS gavage is included in supplementary **Table 4** and **5.**

## Discussion

In this study we investigated the effects of MEV treatment in modulating inflammatory outcomes associated with the NF-κB pathway and NLRP3 inflammasome in male and female neonates exposed to HFD-MO. Male and female neonates exposed to HFD-MO and received MEV treatment exhibited an overall decrease in NF-κB pathway activation and NLRP3 inflammasome formation compared to HFD-MO counterparts who did not receive MEV treatment. Further, protein abundance of candidate pro-inflammatory markers in both pathways returned to baseline with MEV treatment in the liver and hypothalamus. Taken together our data suggests that MEVs provide anti-inflammatory and pro-survival benefits to offspring during postnatal development.

Previous studies have shown protective benefits of exclusive breast/chest feeding on hypothalamic functions (59) and improved metabolic and inflammatory outcomes in the liver (60). However, the anti-inflammatory benefits that can be attributed to MEVs bioactivity in the liver and hypothalamus in HFD-MO neonates remains to be explored. NF-κB signaling pathway control pro-inflammatory response and promotes NLRP3 inflammasome formation. NLRP3 inflammasome leads to pyroptotic cell death and tissue damage (61). NLRP3 inflammasome also play important roles in processing pro-inflammatory precursors such as pre-IL-18 and pre-IL-1β (62) into active cleaved forms (63). Further, HFD-MO exposure increased pro-inflammation in the liver and promoted apoptosis (64, 65) via NF-κB-mediated NLRP3 inflammasome formation.

Liver is a major metabolic organ that governs energy metabolism and the fuel choice o for ATP synthesis. Liver is responsible for macronutrient metabolism, including carbohydrates, fats, and proteins (66) and regulates energy homeostasis of various tissues such as skeletal muscle, adipose tissues, and the brain. Increased adiposity due to HFD-MO exposure is a risk factor for developing non-alcoholic fatty liver disease (NAFLD) in adolescence and adulthood (67). Studies show that breast/chest feeding for at least the first six months of life decreased the occurrence of NAFLD in offspring (68). Liver is also central to extracellular vesicle repackaging and distribution and the pathophysiological role of extracellular vesicles in liver disease and function is well documented (69). Our data indicated that MEV treatment of offspring with HFD-MO exposure led to an attenuation of NF-κB-mediated NLRP3 inflammasome formation in the liver, where protein abundance of TLR4, NF-κB, NLRP3, and pro-caspase-1 returned to baseline levels in males and females, compared to HFD offspring who did not receive MEV treatment. Minimal MEV treatment effects were seen in the CHD neonates across sex.

TLR4 is a toll-like receptor that initiates the activation of the NF-κB pathway by associating with the adaptor protein myeloid differentiating factor 88 (MyD88) (70). TLR4 protein abundance decreased with MEV treatment in the HFD female liver and returned to baseline levels seen in CHD offspring (**Figure 3A&B**). Given TLR4 abundance is strongly associated with the activation of NF-κB (71, 72) this reduction may indicate that MEVs were able to inhibit this signal transduction cascade from the top down. It is likely that the inverse correlation between MEV treatment and decreased protein abundance of TLR4 can be attributed to post-transcriptional activity of miR-148a. miR-148a is a top 10 highly abundant miRNA in human MEVs and bind to the 3’UTR of TLR4 mRNA and inhibit translation (73).

IκBα is the negative regulator of NF-κB and prevents its nuclear translocation. p-Iκκ α/β (S176/180) promote the phosphorylation and subsequent proteasomal degradation of p-IκBα (S32), which facilitates NF-κB nuclear translocation and the increased expression of pro-inflammatory genes, including precursor of interleukins (IL-18 and IL-1β) and NLRP3 (74).

Higher levels of p-Iκκ α/β (S176/180) (**Figure 4**) and lower levels of p-IκBα (S32) (**Figure 5**) was seen in HFD males and females, but this remained unchanged with MEV treatment. A study investigating the effects of MEVs on reversing necrotizing enterocolitis (NEC) in neonatal mice also failed to observe MEV effects on Iκκ and IκBα but showed that MEVs were able to reduce the abundance of IL-1β and NF-κB (29). Nevertheless, immortalized murine microglia polarized with lipopolysaccharide (LPS) and treated with human MEVs illustrated a robust regulation of p-Iκκ α/β (S176/180) and p-IκBα (S32) with MEV treatment, where an overall decrease in both phosphorylation forms were seen (33). These studies combined with ours illustrate the heterogeneity of NF-κB pathway modulation by MEVs during acute versus chronic stress in cells and systems.

The core regulators of the NF-κB and NLRP3 pathways responded to MEV treatment. NF-κB transcription factor promote the production of pro-inflammatory cytokines, such as IL-1β and IL-18 (75) and facilitate the formation of NLRP3 inflammasome (35). Total NF-κB protein returned to baseline in HFD male and female neonates who received MEV treatment, compared to HFD neonates who did not (**Figure 6**). However, limited changes were seen in p-NFκB (S536)/NFκB. NLRP3 (**Figure 7**) and caspase-1 (**Figure 8A&B**) are central components of the NLRP3 inflammasome and are direct transcriptional targets of p-NF-κB (76). Interestingly, both proteins decreased in abundance in HFD males with MEV treatment compared to HFD males who did not receive MEVs (77). Taken together, our data suggested that MEVs may attenuate NLRP3 inflammasome formation in the liver by reducing the availability of two central components of the inflammasome (78). Similar results were reported in a NEC model with neonatal mice, where MEV treatment reduced NLRP3 protein levels in lung tissue (29). In the same study, MEVs and their cargo were shown to alleviate pro-inflammation by reducing caspase-1 production.

The transcript abundance of the pro-inflammatory interleukin, IL-18, a downstream target of NFκB signaling and an integral component of the NLRP3 inflammasome, remained unchanged in response to MEV treatment across sex (**Figure 9**) Filler et al., (2023) reported similar results with IL-18, where IL-18 were most responsive at the protein level compared to transcript (29).

Overall, a strong effect of sex, diet, and MEV treatment was evident in the liver. This correlated to previous studies reporting robust sex-specific responses to HFD-MO in offspring. For example, male neonatal rats with HFD-MO exposure had higher fat accumulation, adiposity, and hyperleptinemia (79), and male offspring were more susceptible to developing obesity (80). Further, pro-inflammatory responses were higher in males, including higher pro-inflammatory cytokines production and secretion (81). However, anti-inflammatory responses in response to drug treatment were also higher in males (82) with increased IL-10 production and TNF-α inhibition (83). We see similar effects herein (**Figure 17**), where male neonates more impacted by HFD-MO exposure and exhibited increased activation of NF-κB mediated NLRP3 inflammasome, yet positively responded to MEV treatment, compared to female neonates at PND 11.

**Figure 17:**
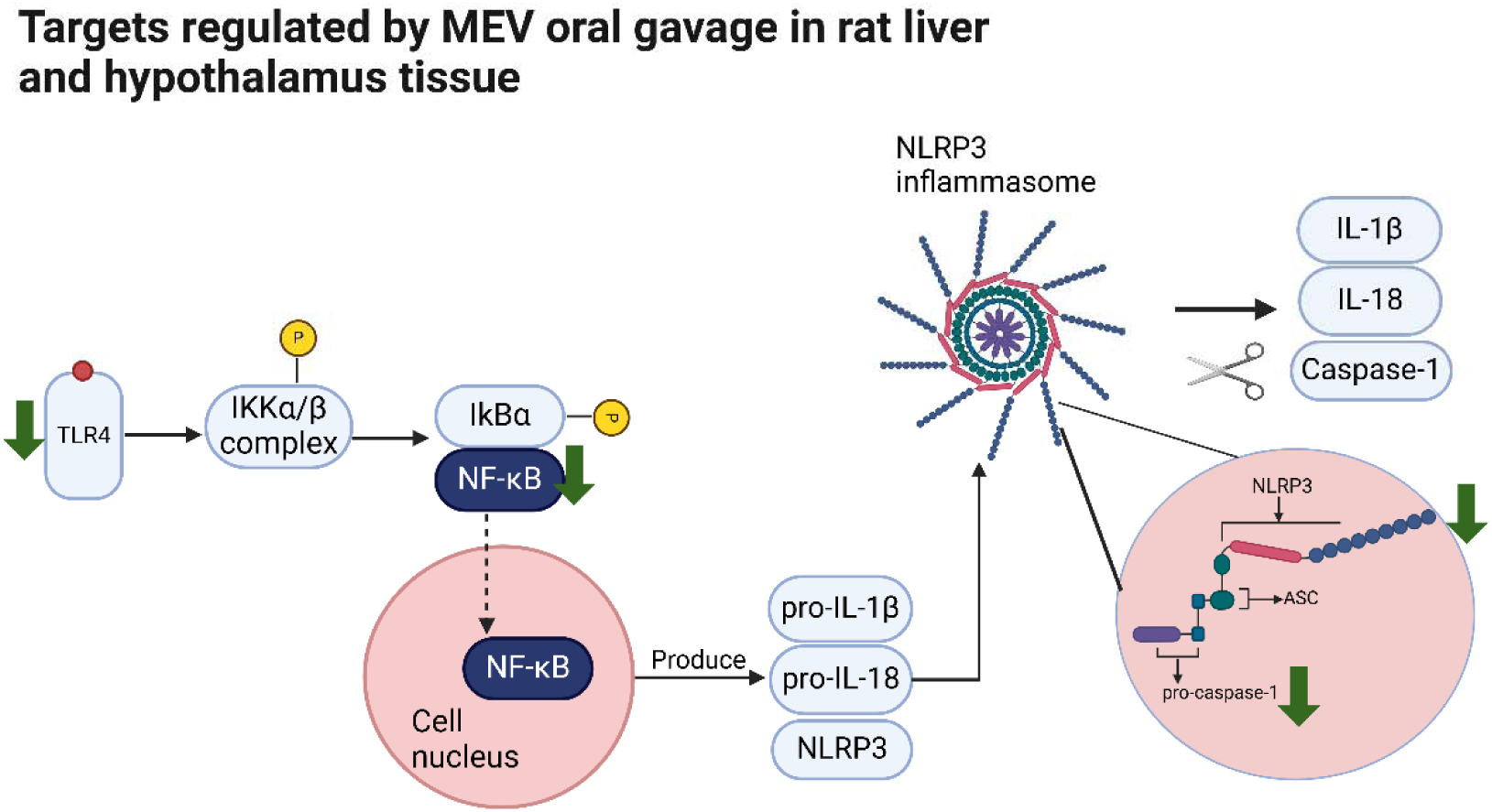
Simplified TLR4 initiated NF-κB signaling activation pathway, showing regulation status of each measured target by MEV oral gavage to Long Evans neonatal rats liver and hypothalamus tissue.

The same targets were measured in the hypothalamus. Hypothalamus is the feeding centre of the brain, and regulates metabolic, endocrine responses, and appetite control (38, 84). Studies had shown that offspring born to dams with gestational diabetes, which was a complication of HFD-MO, exhibited reduced neurogenesis and delayed development of the hypothalamus (85, 86). Thus, with HFD-MO exposure, increased neuroinflammation is expected in the neonatal hypothalamus (87, 88). However, protein abundance of TLR4 in HFD males and females was lower than the CHD counterparts (**Figure 10**) and remained unchanged in response to MEV treatment. Previous studies have also reported similar unexpected finding, where chronic LPS challenge lead to decreased abundance of TLR4 (89). They postulated this to be a result of endotoxin tolerance in the brain (90, 91). Endotoxins that were released from the gut microbiota in response to chronic pro-inflammation are transported to various brain regions and bind repeatedly with TLR4 and induce endotoxin tolerance (87, 88). This could be the reason why we see reduced protein abundance of TLR4 in the hypothalamus of HFD neonates.

Higher transcript and protein levels of Iκκ-β (**Figure 11**) and IκBα (**Figure 12**) were seen in offspring with HFD exposure. This may indicate that although TLR4 abundance is lower, perinatal HFD exposure led to the activation of NF-κB pathway through intermediary steps (92). Iκκ-β and IκBα were also not responsive to MEV treatment in the hypothalamus across sex. This correlated with results in the liver, indicating that similar HFD driven mechanisms were present in metabolically active tissues in neonates.

The transcript and protein abundance of NF-κB (**Figure 13**) remained high in HFD neonates irrespective of MEV treatment. This correlated with p-IκBα (S32) levels in the hypothalamus. The slight increase in p-NF-κB (S536)/NF-κB in the hypothalamus of males with HFD-MEV exposure may indicate that NF-κB activation occurs through alternate pathways (93) with proteins related to IκB family, including IκBγ or Bcl-3 (94).

NLRP3 protein abundance increased in HFD neonates but with MEV treatment the levels returned to baseline and was comparable to CHD offspring (**Figure 14**). This indicates that like the liver, in the hypothalamus MEV treatment attenuates NLRP3 inflammasome formation. This is further confirmed by the protein abundance of pro and cleaved caspase 1. NLRP3 inflammasome when formed lead to the conversion of inactive pro-caspase-1 into active cleaved-caspase-1 and promotes apoptosis (78). In HFD neonates (**Figure 15**), with MEV treatment, an increase abundance of the inactive pro-caspase-I and a lower abundance of the active cleaved-caspase-1 is seen. IL-18 transcript abundance followed a very similar trend in males, where HFD males with MEV treatment had lower levels of IL-18 in the hypothalamus compared to HFD males without MEV treatment (**Figure 16**). Taken together, our data suggests that perinatal HFD exposure promotes NF-κB activation and NLRP3 inflammasome formation in male and female neonates, but MEVs treatment during early postnatal life can revert the pro-inflammatory effects in the hypothalamus.

MEVs exerted anti-inflammatory effects in both liver and hypothalamus in HFD neonates during the stress hyporesponsive period that overlaps with peak lactation in rats. MEV effects were not as prominent in the hypothalamus compared to the liver, likely due to varying rates of distribution and localization of MEVs across tissues and the use of whole hypothalamus for analysis instead of exploring MEV regulation in select nuclei, such as the paraventricular nucleus (25). In addition, offsprings with perinatal HFD exposure exhibited heightened responses to MEV treatment than their CHD counterparts (**Figure 17**). This suggested that MEVs may exert cytoprotective benefits to stressed systems more so than baseline controls in order to promote homeostasis (41, 95). Indeed, the downregulation of NF-κB pathway activation and NLRP3 inflammasome formation in liver may decreased the risk of liver oxidation (40) and hepatic pathologies such as NAFLD (96) in neonates. Cytoprotective benefits in the hypothalamus help to improve endocrine regulation of insulin, leptin, and ghrelin as well as help mitigate the hypothalamic pituitary adrenal axis-induced stress response in neonates during critical periods of early life (97, 98).

## Acknowledgement

We would like to thank Mr. Greg Wasney and the staff at the Structural and Biophysical Core Facility at the Hospital for Sick Children (Toronto, ON, Canada) for performing nanoparticle tracking analysis. We would like to thank the NorthernStar Mothers Milk Bank (Calgary, AB, Canada) for providing the human donor milk that was used for MEV isolations. We would like to thank the staff at the Manitoba Institute for Materials for providing technical support with the FEI Talos F200x S/TEM transmission electron microscope. Mrs. Robyn Cole and Mr. Dan Wasyliw at the University of Winnipeg vivarium for proving support with animal care.

## Abbreviations used in this paper

ASC, Apoptosis-associated speck-like protein containing a CARD; BCA, bicinchoninic acid assay; cDNA, complementary DNA; CHD, Control diet; DoHAD, Developmental origins of Health and Disease; EDTA, ethylenediaminetetraacetic acid; GAPDH, glyceraldehyde-3-phosphate dehydrogenase; GLM, General linear model; GUSB, Glucuronidase beta; HFD, High fat diet; IL-1β, interleukin-1β; IL-18, interleukin-18; IκBα, NF-kappa-B inhibitor alpha; Iκκ, I kappa B kinase complex; MEVs, Milk-derived extracellular vesicles; mRNA, messenger RNA; miRNA, microRNA; MyD88, myeloid differentiating factor 88; NCBI, National Centre for Biotechnology Information; NAFLD, non-alcoholic fatty liver disease; NEC, Necrotizing enterocolitis; NF-κB, nuclear factor κB; NLRP3, NLR family pyrin domain containing 3; NTA, Nanoparticle tracking analysis; MO, maternal obesity; PBS, phosphate-buffered saline; QM, Quantity means; RT-qPCR, Quantitative reverse transcription polymerase chain reaction; SEM, standard error of mean; TBST, 1X Tris-buffered saline with Tween 20; TEM, Transmission electron microscope; TLR-4, toll-like receptor 4; Tukey’s HSD, Tukey’s honestly significant test; YWAZ, tyrosine 3-monooxygenase activation protein zeta.

## Author contribution

JL: Data curation, validation, analysis, writing – original draft, writing – review and editing.

JAS: Data curation.

MFO: Data curation.

SW: Conceptualization, formal analysis, funding acquisition, supervision, writing – original draft, writing – review and editing.

