## Supplemental figures and supplementary tables for "Milk-derived extracellular vesicles mitigate NF-κB pathway and NLRP3 inflammasome formation in Long Evans neonates"

**Supplementary figures**

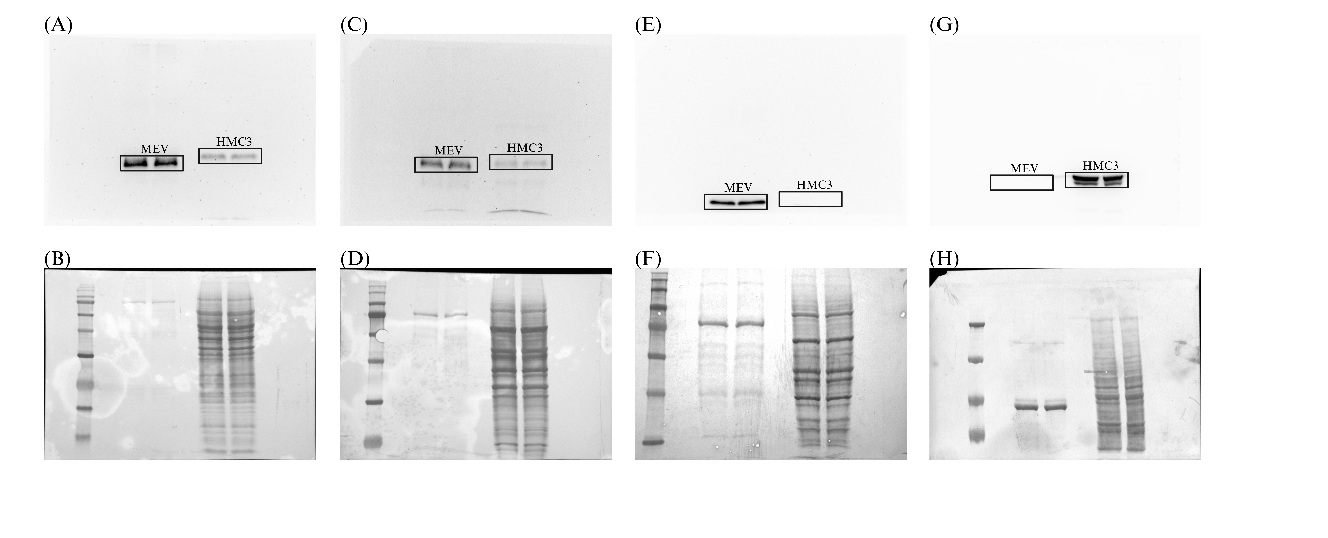

**Figure S1**: Complete western immunoblotting images of MEV characterization targets. (**A**) CD9 enhanced chemiluminescence. (**B**) CD9 coomassie stained membrane. (**C**) CD81 enhanced chemiluminescence. (**D**) CD81 coomassie stained membrane (**E**) Syntenin-1 enhanced chemiluminescence. (**F)** Syntenin-1 coomassie stained membrane. (**G**) Calnexin enhanced chemiluminescence. (**H**) Calnexin coomassie stained membrane. The first lane in all immunoblots represents the protein ladder (5-245 kDa range). IPG = two rodent protein lysates used as the interblot converters. The same samples were included in all electroblots across sex, diets, and treatments. Each protein band represent a single biological replicate. The black boxes identify the target bands.

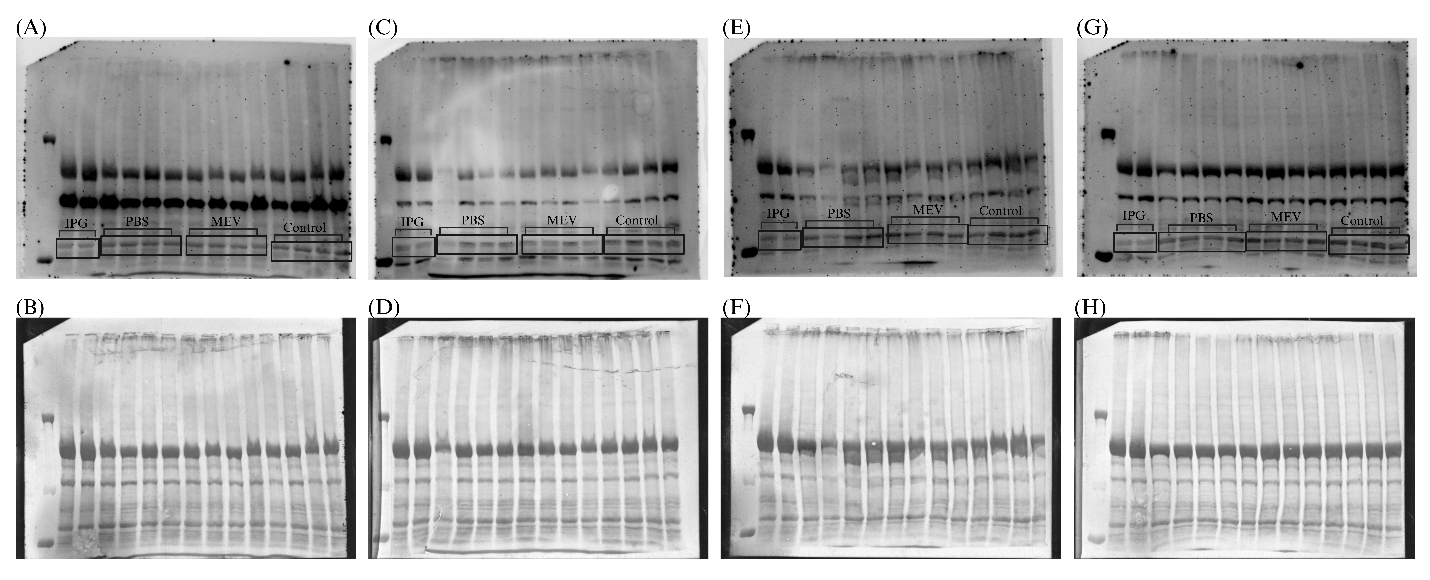

**Figure S2**: Complete western immunoblot images of Toll-like Receptor 4 (TLR4:90 kDa) in the liver of female and male neonates at postnatal day (PND) 11. (**A**) Male neonates with control diet (CHD). (**B**) Male CHD neonates Coomassie stained membrane. (**C**) Male neonates with high fat diet (HFD). (**D**) Male HFD neonates Coomassie stained membrane (**E**) Female neonates with CHD. (**F)** Female CHD neonates Coomassie stained membrane. (**G**) Female neonates with HFD diet. (**H**) Female HFD neonates Coomassie stained membrane. The first lane in all immunoblots represents the protein ladder (10-190 kDa). IPG = two rodent protein lysates used as the interblot converters. The same samples were included in all electroblots across sex, diets, and treatments. PBS = neonates received phosphate-buffered saline as vehicle control, MEV = neonates received milk-derived extracellular vesicles and Control = control neonates that did not receive anything

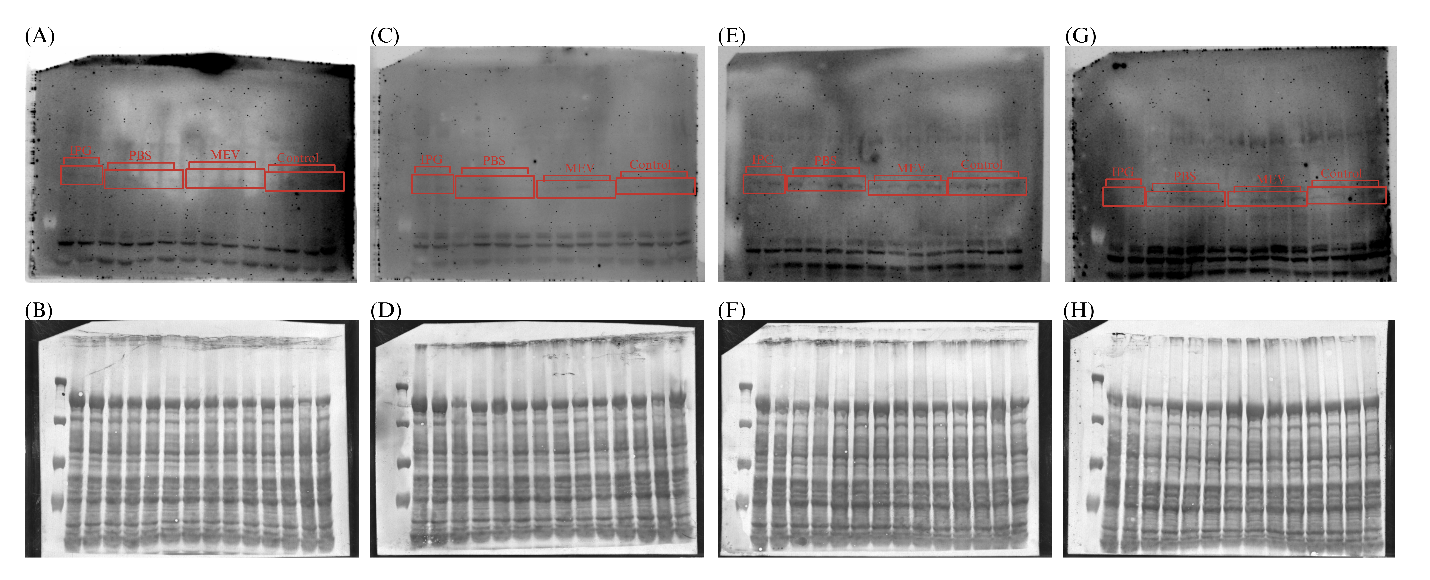

**Figure S3**: Complete western immunoblot images of phosphorylated-IKKα/β at Serine 176/180 (87,88 kDa) at PND11 in neonatal rat liver. (**A**) Male neonates with control diet (CHD). (**B**) Male CHD neonates Coomassie stained membrane. (**C**) Male neonates with high fat diet (HFD). (**D**) Male HFD neonates Coomassie stained membrane (**E**) Female neonates with CHD. (**F)** Female CHD neonates Coomassie stained membrane. (**G**) Female neonates with HFD diet. (**H**) Female HFD neonates Coomassie stained membrane. The first lane in all immunoblots represents the protein ladder (10-190 kDa). IPG = two rodent protein lysates used as the interblot converters. The same samples were included in all electroblots across sex, diets, and treatments. PBS = neonates received phosphate-buffered saline as vehicle control, MEV = neonates received milk-derived extracellular vesicles and Control = control neonates that did not receive anything

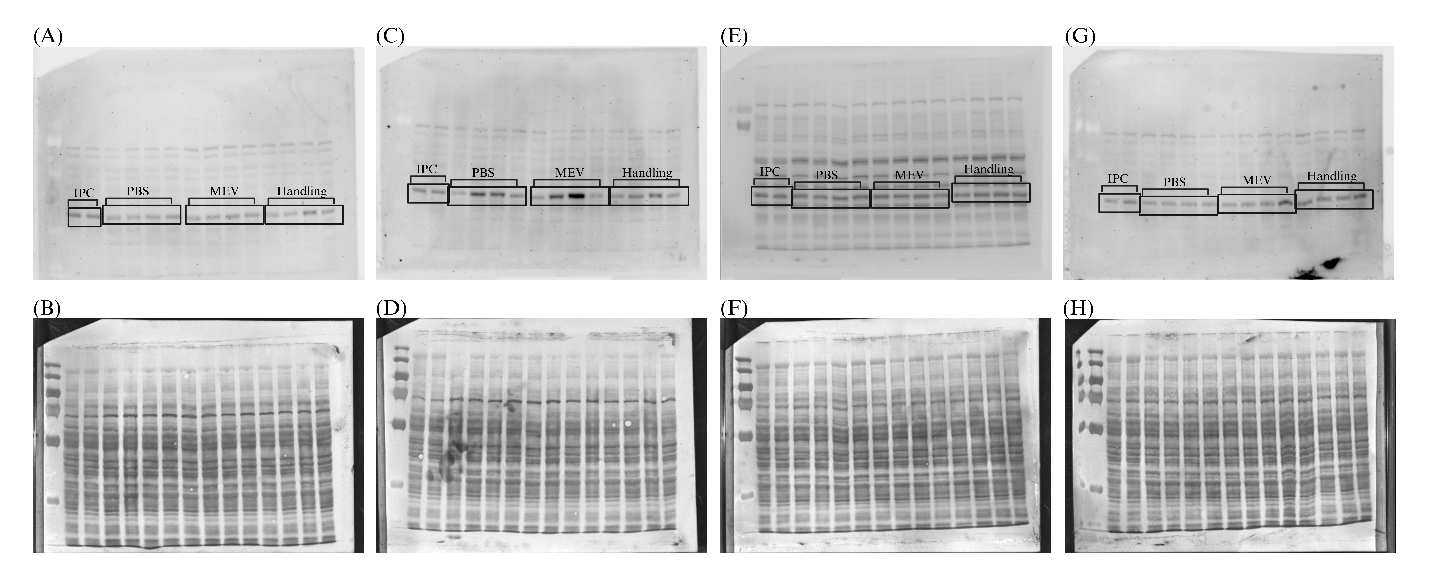

**Figure S4**: Complete western immunoblot images of phosphorylated-IκBα (Serine 32) (39 kDa) at PND11 in neonatal rat liver. (**A**) Male neonates with control diet (CHD). (**B**) Male CHD neonates Coomassie stained membrane. (**C**) Male neonates with high fat diet (HFD). (**D**) Male HFD neonates Coomassie stained membrane (**E**) Female neonates with CHD. (**F)** Female CHD neonates Coomassie stained membrane. (**G**) Female neonates with HFD diet. (**H**) Female HFD neonates Coomassie stained membrane. The first lane in all immunoblots represents the protein ladder (10-190 kDa). IPG = two rodent protein lysates used as the interblot converters. The same samples were included in all electroblots across sex, diets, and treatments. PBS = neonates received phosphate-buffered saline as vehicle control, MEV = neonates received milk-derived extracellular vesicles and Control = control neonates that did not receive anything

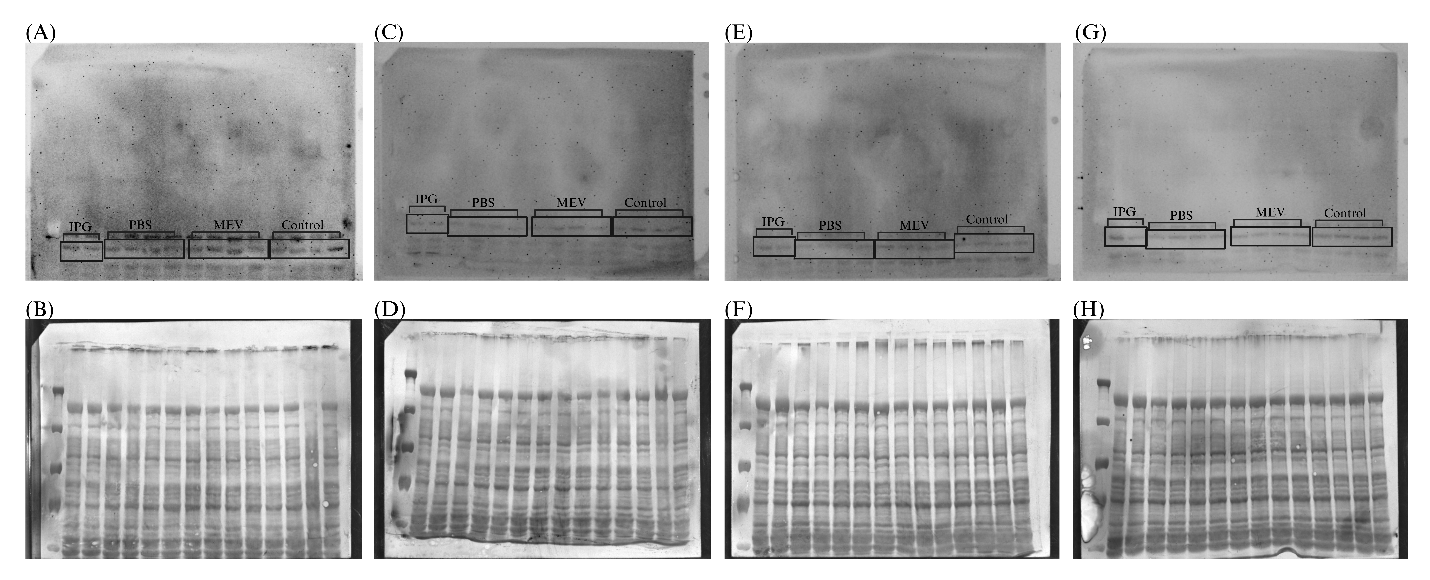

**Figure S5**: Complete western immunoblot images of NF-κB (65 kDa). (**A**) Male neonates with control diet (CHD). (**B**) Male CHD neonates Coomassie stained membrane. (**C**) Male neonates with high fat diet (HFD). (**D**) Male HFD neonates Coomassie stained membrane (**E**) Female neonates with CHD. (**F)** Female CHD neonates Coomassie stained membrane. (**G**) Female neonates with HFD diet. (**H**) Female HFD neonates Coomassie stained membrane. The first lane in all immunoblots represents the protein ladder (10-190 kDa). IPG = two rodent protein lysates used as the interblot converters. The same samples were included in all electroblots across sex, diets, and treatments. PBS = neonates received phosphate-buffered saline as vehicle control, MEV = neonates received milk-derived extracellular vesicles and Control = control neonates that did not receive anything

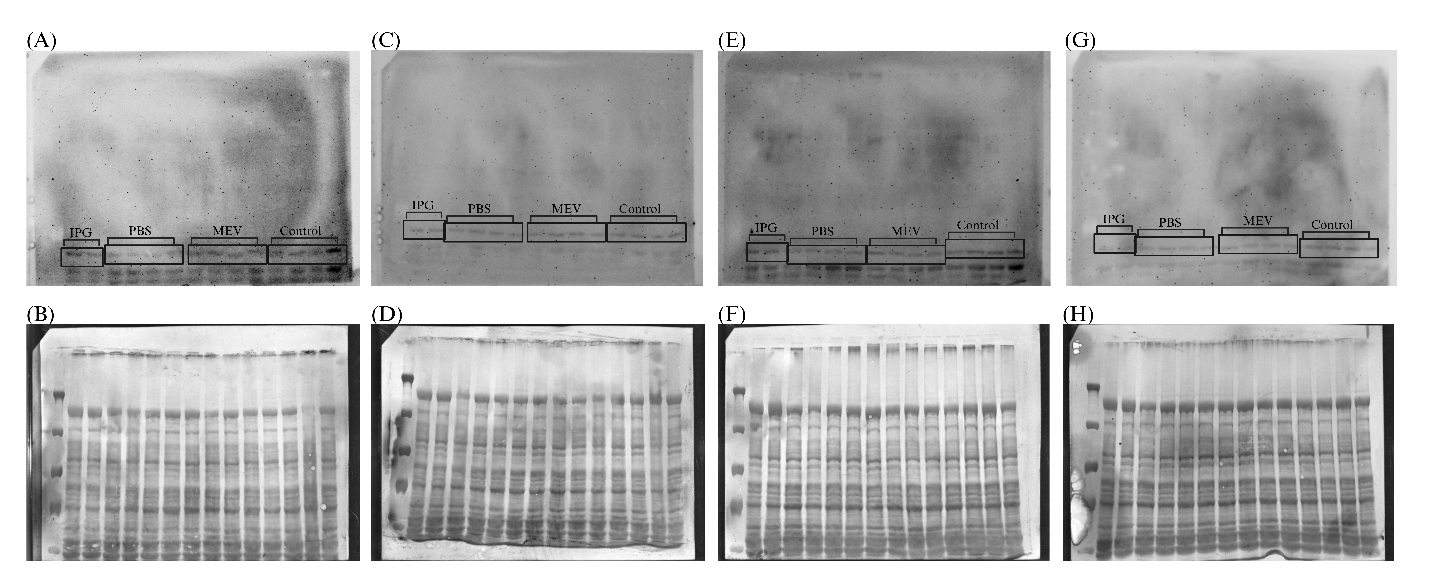

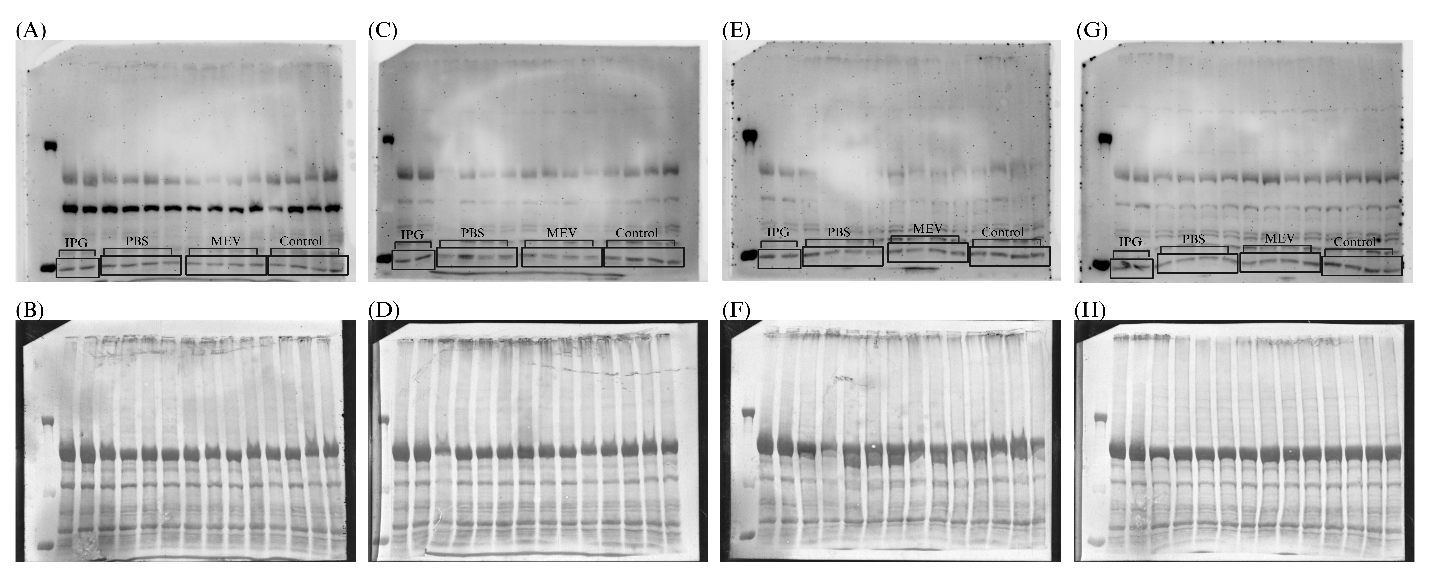
**Figure S6**: Complete western immunoblot images of phosphorylated-NF-κB at Serine 536 (65 kDa) at PND11 in neonatal rat liver. (**A**) Male neonates with control diet (CHD). (**B**) Male CHD neonates Coomassie stained membrane. (**C**) Male neonates with high fat diet (HFD). (**D**) Male HFD neonates Coomassie stained membrane (**E**) Female neonates with CHD. (**F)** Female CHD neonates Coomassie stained membrane. (**G**) Female neonates with HFD diet. (**H**) Female HFD neonates Coomassie stained membrane. The first lane in all immunoblots represents the protein ladder (10-190 kDa). IPG = two rodent protein lysates used as the interblot converters. The same samples were included in all electroblots across sex, diets, and treatments. PBS = neonates received phosphate-buffered saline as vehicle control, MEV = neonates received milk-derived extracellular vesicles and Control = control neonates that did not receive anything

**Figure S7**: Complete western immunoblot images of NLRP3 (100 kDa) at PND11 in neonatal rat liver. (**A**) Male neonates with control diet (CHD). (**B**) Male CHD neonates Coomassie stained membrane. (**C**) Male neonates with high fat diet (HFD). (**D**) Male HFD neonates Coomassie stained membrane (**E**) Female neonates with CHD. (**F)** Female CHD neonates Coomassie stained membrane. (**G**) Female neonates with HFD diet. (**H**) Female HFD neonates Coomassie stained membrane. The first lane in all immunoblots represents the protein ladder (10-190 kDa). IPG = two rodent protein lysates used as the interblot converters. The same samples were included in all electroblots across sex, diets, and treatments. PBS = neonates received phosphate-buffered saline as vehicle control, MEV = neonates received milk-derived extracellular vesicles and Control = control neonates that did not receive anything

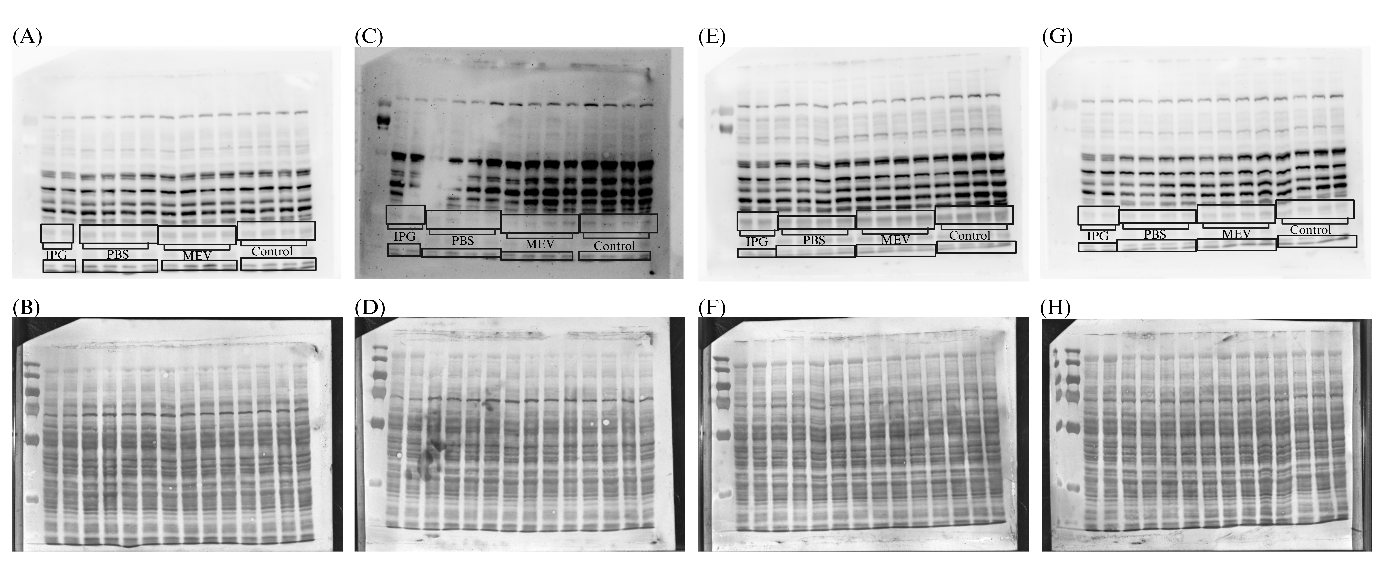

**Figure S8**: Complete western immunoblot images of caspase-1 (25, 35 kDa) at PND11 in neonatal rat liver. (**A**) Male neonates with control diet (CHD). (**B**) Male CHD neonates Coomassie stained membrane. (**C**) Male neonates with high fat diet (HFD). (**D**) Male HFD neonates Coomassie stained membrane (**E**) Female neonates with CHD. (**F)** Female CHD neonates Coomassie stained membrane. (**G**) Female neonates with HFD diet. (**H**) Female HFD neonates Coomassie stained membrane. The first lane in all immunoblots represents the protein ladder (10-190 kDa). IPG = two rodent protein lysates used as the interblot converters. The same samples were included in all electroblots across sex, diets, and treatments. PBS = neonates received phosphate-buffered saline as vehicle control, MEV = neonates received milk-derived extracellular vesicles and Control = control neonates that did not receive anything

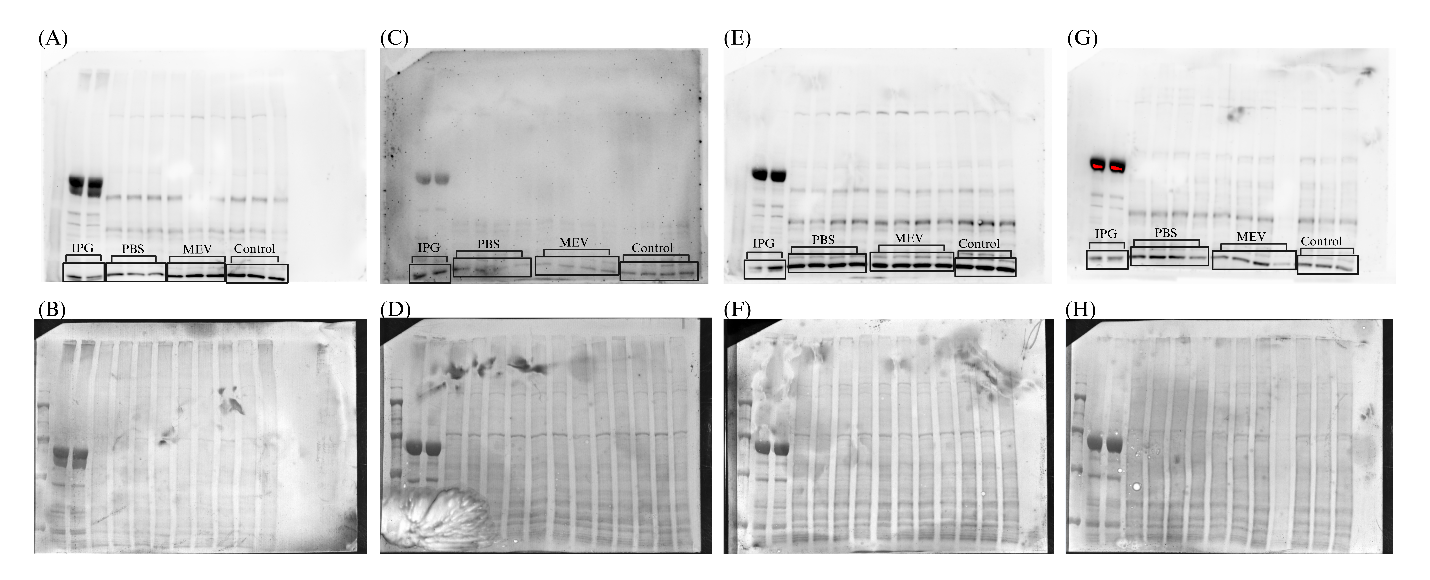

**Figure S9**: Complete western immunoblot images of Toll-like Receptor 4 (TLR4:90 kDa) at PND11 in neonatal rat hypothalamus tissues. (**A**) Male neonates with control diet (CHD). (**B**) Male CHD neonates Coomassie stained membrane. (**C**) Male neonates with high fat diet (HFD). (**D**) Male HFD neonates Coomassie stained membrane (**E**) Female neonates with CHD. (**F)** Female CHD neonates Coomassie stained membrane. (**G**) Female neonates with HFD diet. (**H**) Female HFD neonates Coomassie stained membrane. The first lane in all immunoblots represents the protein ladder (10-190 kDa). IPG = two rodent protein lysates used as the interblot converters. The same samples were included in all electroblots across sex, diets, and treatments. PBS = neonates received phosphate-buffered saline as vehicle control, MEV = neonates received milk-derived extracellular vesicles and Control = control neonates that did not receive anything

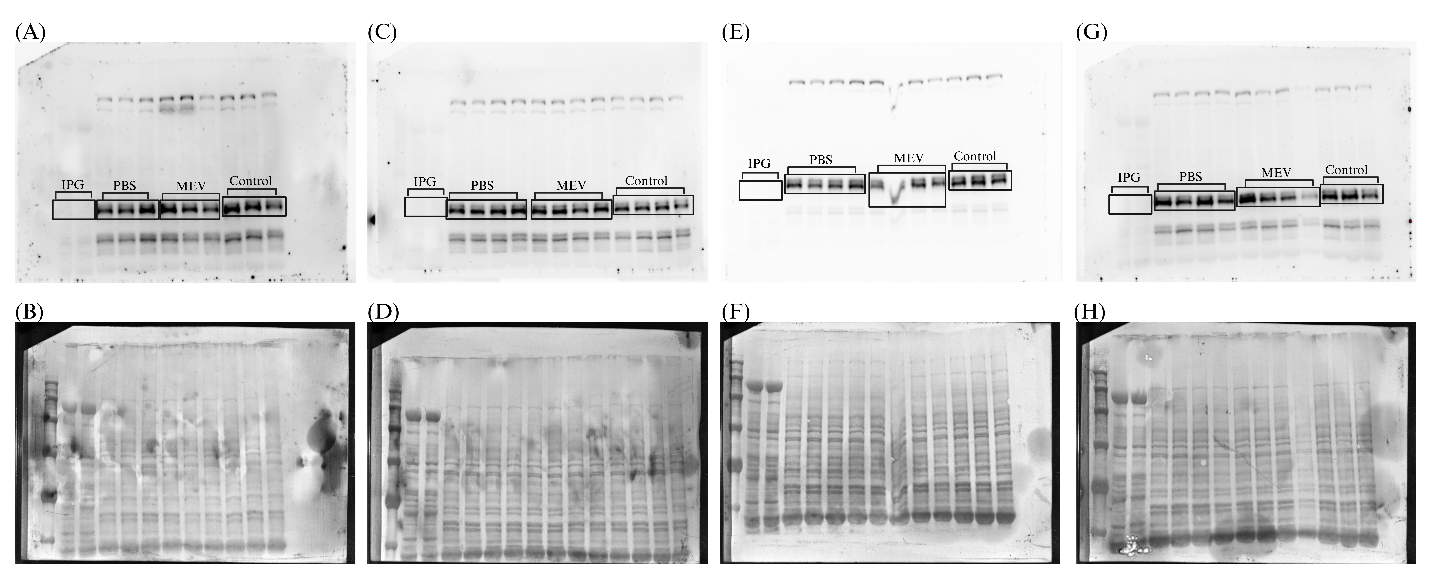

**Figure S10**: Complete western immunoblot images of phosphorylated-IKKα/β at Serine 176/180 (87,88 kDa) at PND11 in neonatal rat hypothalamus tissues. (**A**) Male neonates with control diet (CHD). (**B**) Male CHD neonates Coomassie stained membrane. (**C**) Male neonates with high fat diet (HFD). (**D**) Male HFD neonates Coomassie stained membrane (**E**) Female neonates with CHD. (**F)** Female CHD neonates Coomassie stained membrane. (**G**) Female neonates with HFD diet. (**H**) Female HFD neonates Coomassie stained membrane. The first lane in all immunoblots represents the protein ladder (10-190 kDa). IPG = two rodent protein lysates used as the interblot converters. The same samples were included in all electroblots across sex, diets, and treatments. PBS = neonates received phosphate-buffered saline as vehicle control, MEV = neonates received milk-derived extracellular vesicles and Control = control neonates that did not receive anything.

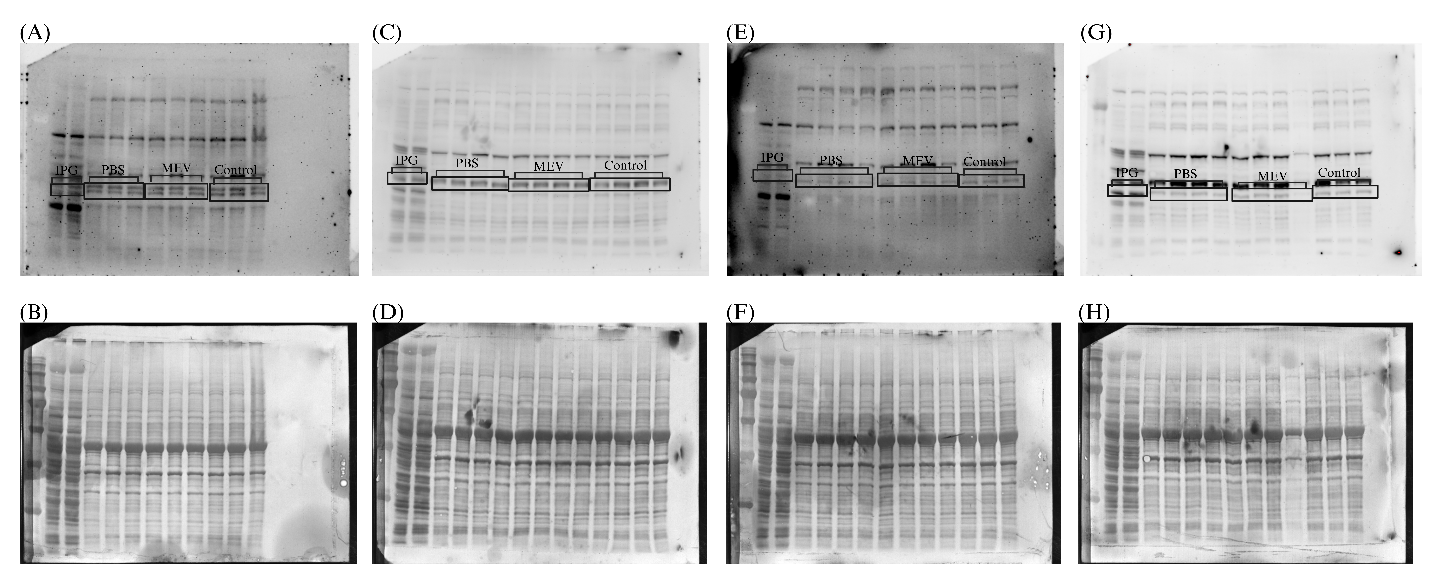

**Figure S11**: Complete western immunoblot images of phosphorylated-IκBα (Serine 32) (39 kDa) at PND11 in neonatal rat hypothalamus tissues. (**A**) Male neonates with control diet (CHD). (**B**) Male CHD neonates Coomassie stained membrane. (**C**) Male neonates with high fat diet (HFD). (**D**) Male HFD neonates Coomassie stained membrane (**E**) Female neonates with CHD. (**F)** Female CHD neonates Coomassie stained membrane. (**G**) Female neonates with HFD diet. (**H**) Female HFD neonates Coomassie stained membrane. The first lane in all immunoblots represents the protein ladder (10-190 kDa). IPG = two rodent protein lysates used as the interblot converters. The same samples were included in all electroblots across sex, diets, and treatments. PBS = neonates received phosphate-buffered saline as vehicle control, MEV = neonates received milk-derived extracellular vesicles and Control = control neonates that did not receive anything.

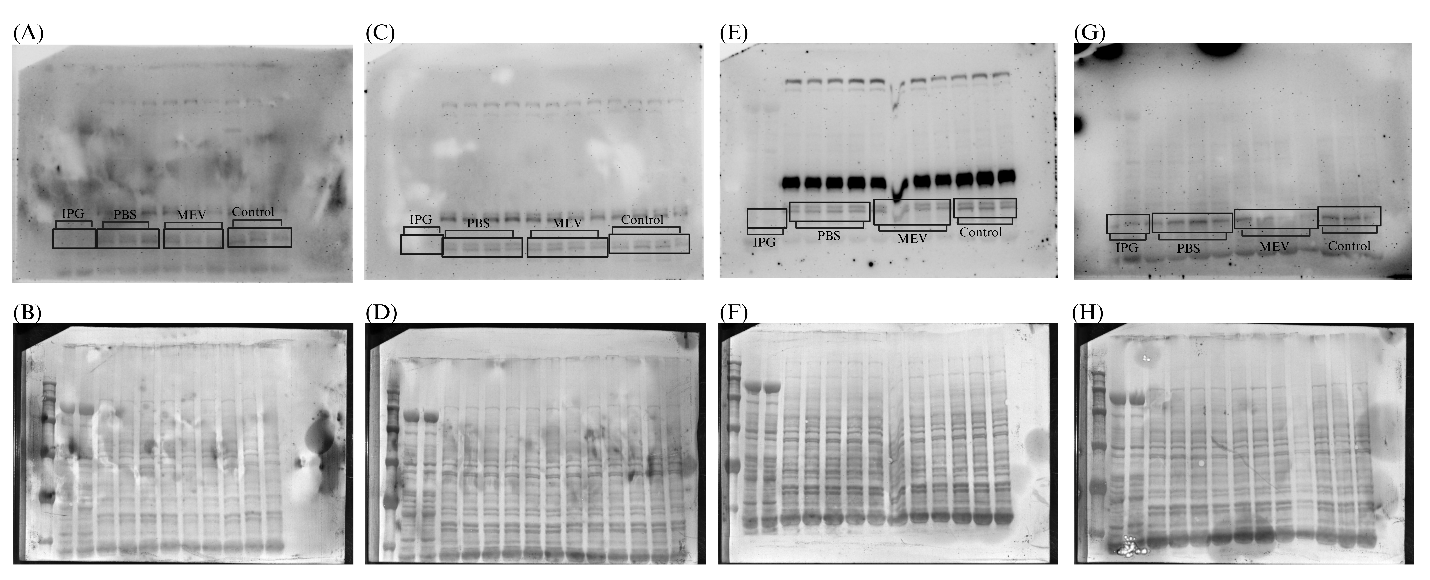

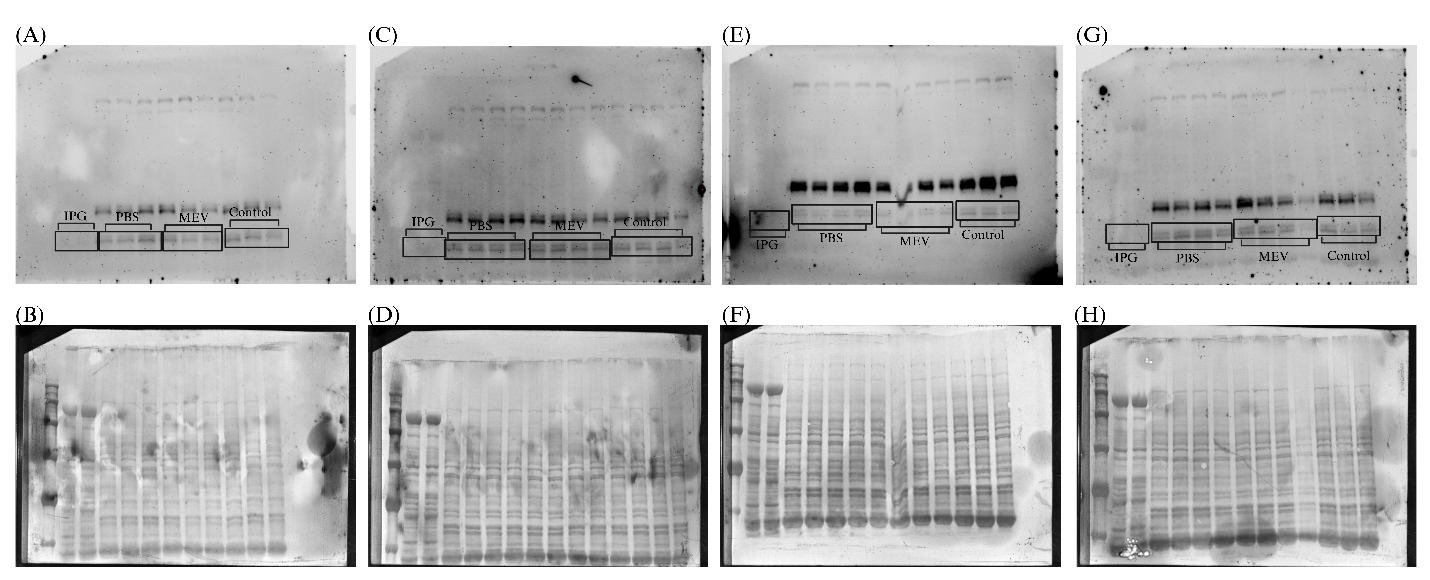
**Figure S12**: Complete western immunoblot images of NF-κB (65 kDa) at PND11 in neonatal rat hypothalamus tissues. (**A**) Male neonates with control diet (CHD). (**B**) Male CHD neonates Coomassie stained membrane. (**C**) Male neonates with high fat diet (HFD). (**D**) Male HFD neonates Coomassie stained membrane (**E**) Female neonates with CHD. (**F)** Female CHD neonates Coomassie stained membrane. (**G**) Female neonates with HFD diet. (**H**) Female HFD neonates Coomassie stained membrane. The first lane in all immunoblots represents the protein ladder (10-190 kDa). IPG = two rodent protein lysates used as the interblot converters. The same samples were included in all electroblots across sex, diets, and treatments. PBS = neonates received phosphate-buffered saline as vehicle control, MEV = neonates received milk-derived extracellular vesicles and Control = control neonates that did not receive anything.

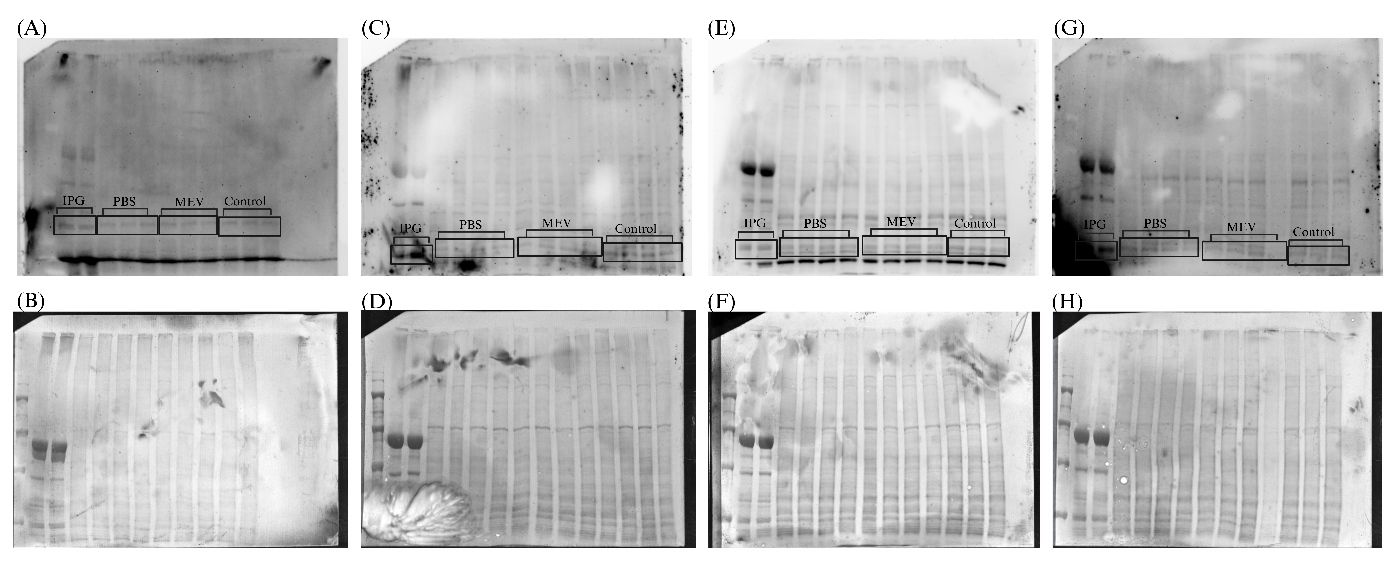
**Figure S13**: Complete western immunoblot images of phosphorylated-NF-κB at Serine 536 (65 kDa) at PND11 in neonatal rat hypothalamus tissues. (**A**) Male neonates with control diet (CHD). (**B**) Male CHD neonates Coomassie stained membrane. (**C**) Male neonates with high fat diet (HFD). (**D**) Male HFD neonates Coomassie stained membrane (**E**) Female neonates with CHD. (**F)** Female CHD neonates Coomassie stained membrane. (**G**) Female neonates with HFD diet. (**H**) Female HFD neonates Coomassie stained membrane. The first lane in all immunoblots represents the protein ladder (10-190 kDa). IPG = two rodent protein lysates used as the interblot converters. The same samples were included in all electroblots across sex, diets, and treatments. PBS = neonates received phosphate-buffered saline as vehicle control, MEV = neonates received milk-derived extracellular vesicles and Control = control neonates that did not receive anything.

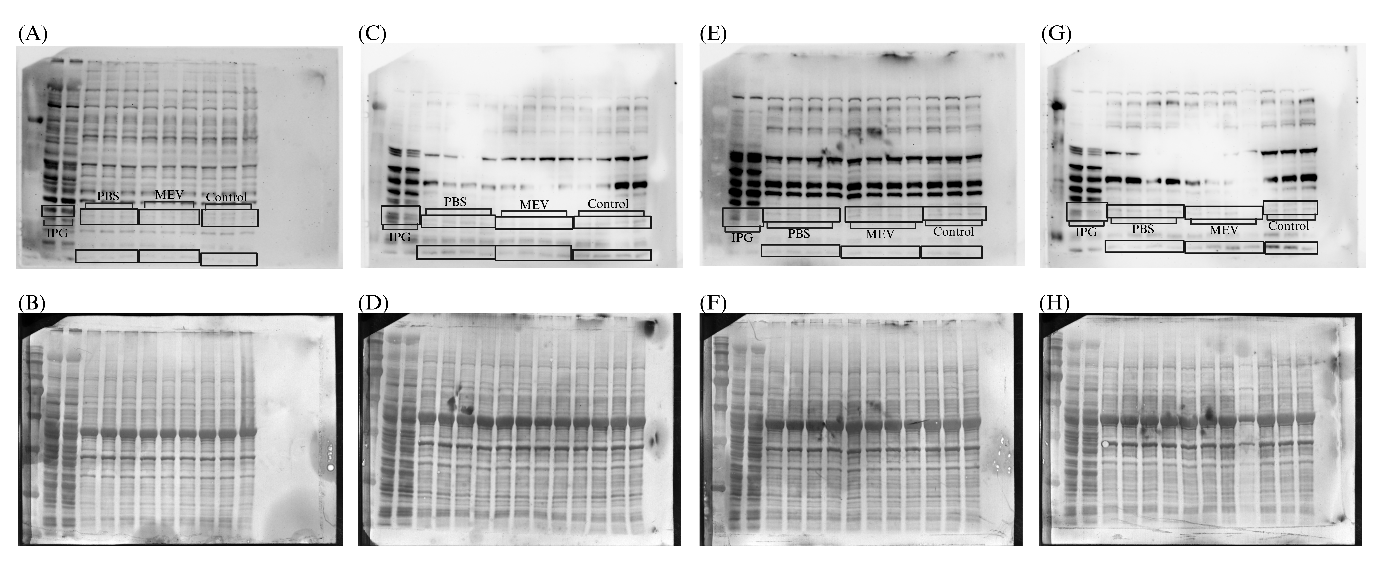
**Figure S14**: Complete western immunoblot images of NLRP3 (100kDa) at PND11 in neonatal rat hypothalamus tissues. (**A**) Male neonates with control diet (CHD). (**B**) Male CHD neonates Coomassie stained membrane. (**C**) Male neonates with high fat diet (HFD). (**D**) Male HFD neonates Coomassie stained membrane (**E**) Female neonates with CHD. (**F)** Female CHD neonates Coomassie stained membrane. (**G**) Female neonates with HFD diet. (**H**) Female HFD neonates Coomassie stained membrane. The first lane in all immunoblots represents the protein ladder (10-190 kDa). IPG = two rodent protein lysates used as the interblot converters. The same samples were included in all electroblots across sex, diets, and treatments. PBS = neonates received phosphate-buffered saline as vehicle control, MEV = neonates received milk-derived extracellular vesicles and Control = control neonates that did not receive anything.

**Figure S15**: Complete western immunoblot images of caspase-1 (25, 35 kDa) at PND11 in neonatal rat hypothalamus tissues. (**A**) Male neonates with control diet (CHD). (**B**) Male CHD neonates Coomassie stained membrane. (**C**) Male neonates with high fat diet (HFD). (**D**) Male HFD neonates Coomassie stained membrane (**E**) Female neonates with CHD. (**F)** Female CHD neonates Coomassie stained membrane. (**G**) Female neonates with HFD diet. (**H**) Female HFD neonates Coomassie stained membrane. The first lane in all immunoblots represents the protein ladder (10-190 kDa). IPG = two rodent protein lysates used as the interblot converters. The same samples were included in all electroblots across sex, diets, and treatments. PBS = neonates received phosphate-buffered saline as vehicle control, MEV = neonates received milk-derived extracellular vesicles and Control = control neonates that did not receive anything.

**Table 1**: Primary and secondary antibody information for MEV biomarkers and NF-κB pathway markers. Targets are identified by their commercial protein names, company and catalogue number, species reactivity, molecular weight (kDa) and host species.

| Type of marker | Protein | Company | Catalogue # | Reactivity | Molecular weight (kDa) | Host |
| --- | --- | --- | --- | --- | --- | --- |
| Primary antibody for  MEV characterization | CD9 | Cell signaling  Exosomal marker antibody sampler kit | 74220T | Human | 22 | Rabbit |
|  | CD81 |  |  | Human | 22, 25, 35 |  |
|  | Synthenin-1 |  |  | Human, mouse, rat | 30 |  |
| Primary antibody for  Western Immunoblotting | TLR4 | Abcam |  | Human | 90-100 | Rabbit |
|  | Phospho-Iκκ α/β | Cell signaling  NF-κB pathway antibody sampler kit | 9936T | Human, mouse, rat | 90 | Rabbit |
|  | Phospho-IκBα |  |  |  | 39 |  |
|  | NF-κB |  |  |  | 65 |  |
|  | Phospho- NF-κB |  |  |  | 65 |  |
|  | NLRP3 | ThermoFisher | PA5-115660 | Human, mouse, rat | 100 | Rabbit |
|  | Pro-caspase-1 | ProteinTech | 22915-1-AP | Human, mouse, rat | 35 | Rabbit |
|  | Cleaved-caspase-1 |  |  |  | 25 |  |

**Table 2**: Primer sequences for rat liver and hypothalamus RT-qPCR. Targets are identified by gene names, NCBI accession numbers, product size (bp), melting temperature (⁰C), GC content (%), hairpin temperature (⁰C), homodimer (%) and heterodimer (%).

| Type of marker | Gene | NCBI Accession | Forward Primer (5’ – 3’) | Reverse Primer (5’ – 3’) | Product size (bp) | Melting temperature forward (⁰C) | Melting temperature reverse (⁰C) | GC content forward (%) | GC content reverse (%) | Hairpin temperature (⁰C) | Homodimer (%) | Heterodimer (%) | Source of primer |
| --- | --- | --- | --- | --- | --- | --- | --- | --- | --- | --- | --- | --- | --- |
| Reference gene (used for normalization) | *YWAZ* | NM_013011.4 | TTGAGCAGAAGACGGAAGGT | GAAGCATTGGGGATCAAGAA | 136 | 63.3 | 60.3 | 50.0 | 45.0 | 18.7 | 10.4 | 11.60 | Designed |
|  | *GAPDH* | NM_017008.4 | ACATCAAATGGGGTGATGCT | GTGGTTCACACCCATCACAA | 161 | 62.3 | 62.6 | 45.0 | 50.0 | 49.8 | 9.50 | 25.28 | Designed |
|  | *GUSB* | NC_086030.1 | CATGACGAACCAGTCACCAC | ACGGTCTGCTTCCCATACAC | 107 | 62.6 | 63.8 | 55.0 | 55.0 | 48 | 14.96 | 11.63 | (L. Liu et al., 2020) |
| Target marker | *TLR4* | NM_019178.2 | TCCCTGGTGTTGGATTTTACGA | TCATAGATGCTTTCTCCTCTGCTG | 176 | 62.9 | 63.9 | 45.5 | 45.8 | 51.1 | 10.97 | 10.94 | (Tian et al., 2021) |
|  | *IKK-β* | NM_053355.3 | TGAAGGTCCTGGTAGAACGG | GCCTTCTGCTTACAAACCACG | 277 | 56.5 | 63.7 | 55.0 | 52.4 | 43.1 | 12.22 | 19.96 | Designed |
|  | *IκBα* | NM_001105720.2 | GCCCTACGATGACTGTGTGTT | CTGTTTCCCCAAATTTCACAA | 186 | 64.5 | 59.8 | 52.4 | 38.1 | 44.5 | 9.47 | 13.28 | (Klimas et al., 2021) |
|  | *NF-Κb* | NM_199267.2 | GAGACCTTCAACACCCCAGC | ATGTCACGCACGATTTCCC | 263 | 64.4 | 62.7 | 60.0 | 52.6 | 43.5 | 8.11 | 8.52 | (Al-Rasheed et al., 2016) |
|  | *NLRP3* | NM_001191642.1 | GACCTCAACAGACGCTACACCCA | GTCCCACATCTTAGTCCTGCCAA | 105 | 63..8 | 65.8 | 56.5 | 52.2 | 28.4 | 8.41 | 8.24 | (Tian et al., 2021) |
|  | *IL-18* | NM_019165.2 | AGAAGAAGGCTCTTGTGTCAACTT | TACAGGAGAGGGAGACATCCTTC | 112 | 59.1 | 64.2 | 41.7 | 52.2 | 46.4 | 12.28 | 19.64 | (H. Chang et al., 2014) |

**Table 3**: Parameters of primary antibodies used for MEV characterization and western immunoblotting. Targets are identified by protein names, company and catalogue numbers.

| Protein name | Company and catalogue # | Protein amount (μg) | Gel % | Transfer settings | Blocking solution | Primary antibody ratio | Secondary antibody ratio |
| --- | --- | --- | --- | --- | --- | --- | --- |
| CD9 | Cell signaling  Exosomal marker antibody sampler kit (74220T) | 20  20  30 | 15% | Turbo 7 minutes | 1% casein for 30 min  2.5% casein for 30 min  1% casein for 30 min | 1:1000 24hours at 4 ⁰C | 1:15000 45minutes at room temperature |
| CD81 |  |  | 15% | Turbo 10 minutes |  | 1:1000 24hours at 4 ⁰C | 1:15000 45minutes at room temperature |
| Synthenin-1 |  |  | 10% | Standard 20 minutes, 2.5amps |  | 1:1000 24hours at 4 ⁰C | 1:15000 45minutes at room temperature |
| TLR4 | Abcam (ab13556) | 10 | 6% | Standard 10 minutes, 2.5amps | 10% casein for 30 min | 1:1000 24hours at 4 ⁰C | 1:15000 45minutes at room temperature |
| Phospho-Iκκ α/β | Cell signaling  NF-κB pathway antibody sampler kit (9936T) | 20 | 8% | Standard 15 minutes, 2.5amps | 1% casein for 30 min | 1:1000 1 hour at room temperature | 1:5000 1 hour and 15 minutes at room temperature |
| Phospho-IκBα |  | 20 | 10% | Standard 10 minutes, 2.5amps | 2.5% casein for 1 hour | 1:1000 1 hour at room temperature | 1:10000 45minutes at room temperature |
| NF-κB |  | 10 | 10% | Standard 40 minutes, 2.5amps | 10% casein for 1 hour | 1:1000 24hours at 4 ⁰C | 1:10000 45minutes at room temperature |
| Phospho- NF-κB |  | 20 | 8% | Standard 15 minutes, 2.5amps | 1% casein for 30 min | 1:1000 1 hour at room temperature | 1:5000 1 hour and 15 minutes at room temperature |
| NLRP3 | ThermoFisher (PA5-115660) | 25 | 6% | Standard 25 minutes, 2.5amps | 1% casein for 30 min | 1:1000 24hours at 4 ⁰C | 1:10000 45minutes at room temperature |
| Pro-caspase-1 | ProteinTech (22915-1-AP) | 10 | 10% | Standard 20 minutes, 2.5amps | 10% casein for 1 hour | 1:1000 24hours at 4 ⁰C | 1:12000 45minutes at room temperature |
| Cleaved-caspase-1 |  |  |  |  |  |  |  |

**Table 4**: Normalized transcript abundance data and normalized protein abundance data for NF-kb pathway targets in rat liver

| Treatment groups | Sample IDs | Normalized transcript abundance | | | | | | | Normalized protein abundance | | | | | | | |
| --- | --- | --- | --- | --- | --- | --- | --- | --- | --- | --- | --- | --- | --- | --- | --- | --- |
|  |  | TLR4 | Iκκ β | IκBα | NF--κB | NLRP3 | Caspase-1 | IL-18 | TLR4 | P-Iκκ β (S176/180) | P-IκBα (S32) | NF-κB | P-NF-κB (S536) | NLRP3 | Pro-caspase-1 | Cleaved-caspase-1 |
| mCHD-PBS | D1M1 | 1.10E+00 | 7.68E-01 | 1.10E-01 | 8.71E-01 | 8.37E-01 | 1.04E+00 | 6.57E-01 | 0.039647893 | 0.195126701 | 0.345757228 | 0.173700556 | 0.040714828 | 0.931991315 | 0.011016371 | 0.110519765 |
| mCHD-PBS | D3M1 | 8.10E-01 | 7.75E-01 | 8.10E-01 | 1.15E+00 | 1.06E+00 | 5.20E-01 | 4.37E-01 | 0.297240877 | 0.069252692 | 0.38623644 | 0.254438687 | 0.234249245 | 1.539822768 | 0.094835085 | 0.028426756 |
| mCHD-PBS | D9M1 | 9.63E-01 | 1.06E+00 | 9.63E-01 | 1.17E+00 | 1.53E+00 | 1.32E+00 | 5.83E-01 | 0.099962213 | 0.09467904 | 0.329033849 | 0.514192236 | 0.202612675 | 1.376459983 | 0.080807386 | 0.028549857 |
| mCHD-PBS | D11M2 | 7.73E-01 | 7.23E-01 | 7.73E-01 | 1.04E+00 | 1.02E+00 | 9.66E-01 | 4.70E-01 | 0.000798307 | 0.006125399 | 0.33096478 | 0.552064676 | 0.172971653 | 1.294377997 | 0.0222807 | 0.116153965 |
| mCHD-MEV | D1M4 | 7.17E-01 | 7.28E-01 | 7.17E-01 | 1.18E+00 | 5.23E-01 | 5.60E-01 | 3.62E-01 | 0.207226365 | 0.061046339 | 0.410077954 | 0.352979156 | 0.360295847 | 1.014642144 | 0.00992458 | 0.036691758 |
| mCHD-MEV | D3M2 | 1.00E+00 | 1.01E+00 | 1.00E+00 | 1.31E+00 | 1.10E+00 | 1.10E+00 | 7.02E-01 | 0.205204943 | 0.085604076 | 0.390634014 | 0.312342555 | 0.361427534 | 1.097369165 | 0.011150031 | 0.204047974 |
| mCHD-MEV | D9M4 | 1.00E+00 | 1.03E+00 | 1.00E+00 | 1.09E+00 | 2.06E+00 | 1.65E+00 | 1.04E+00 | 0.014089806 | 0.121267661 | 0.311437323 | 0.317404648 | 0.406945986 | 0.971754196 | 0.01612292 | 0.585642308 |
| mCHD-MEV | D11M3 | 1.75E+00 | 1.24E+00 | 1.75E+00 | 1.01E+00 | 1.61E+00 | 2.02E+00 | 1.16E+00 | 0.149089455 | 0.021059601 | 0.303163864 | 0.100321846 | 0.568429791 | 0.913885561 | 0.064991851 | 0.20252509 |
| mCHD-handle | D1M6 | 1.06E+00 | 1.02E+00 | 1.06E+00 | 1.16E+00 | 9.64E-01 | 1.02E+00 | 4.90E-01 | 0.107749951 | 0.164411131 | 0.410578507 | 0.407226322 | 1.084800411 | 1.144010255 | 0.066532771 | 0.12760006 |
| mCHD-handle | D3M3 | 5.99E-01 | 8.36E-01 | 5.99E-01 | 1.46E+00 | 1.07E+00 | 6.01E-01 | 3.67E-01 | 0.085231639 | 0.032202511 | 0.47712932 | 0.085878852 | 0.552767892 | 1.08498926 | 0.058407655 | 0.084358192 |
| mCHD-handle | D9M6 | 7.77E-01 | 1.03E+00 | 7.77E-01 | 1.24E+00 | 1.61E+00 | 9.94E-01 | 4.78E-01 | 0.025426956 | 0.02174777 | 0.339933598 | 0.28603512 | 0.327050957 | 1.069514302 | 0.112299599 | 0.132354236 |
| mCHD-handle | D11M6 | 7.84E-01 | 8.05E-01 | 7.84E-01 | 1.37E+00 | 1.38E+00 | 1.18E+00 | 5.66E-01 | 0.018782268 | 0.055139556 | 0.424118659 | 0.030537765 | 0.482406809 | 1.115812225 | 0.066784368 | 0.078239498 |
| mHFD-PBS | D2M2 | 3.99E-01 | 5.98E-01 | 3.99E-01 | 7.59E-01 | 6.63E-01 | 6.25E-01 | 4.68E-01 | 0.176903978 | 0.077655306 | 0.208944988 | 0.022157467 | 0.089764124 |  | 0.255766245 | 0.015997763 |
| mHFD-PBS | D4M1 | 5.92E-01 | 8.13E-01 | 5.92E-01 | 7.19E-01 | 8.26E-01 | 8.94E-01 | 5.87E-01 | 0.247535317 | 0.204740611 | 0.438279772 | 0.016960677 | 0.023452236 | 0.131259899 | 0.158937352 | 0.041624084 |
| mHFD-PBS | D6M1 | 7.14E-01 | 8.62E-01 | 7.14E-01 | 1.21E+00 | 9.77E-01 | 7.52E-01 | 5.39E-01 | 0.059676893 | 0.04292085 | 0.31985705 | 0.035888714 | 0.009933102 | 0.085983832 | 0.30055906 | 0.154003875 |
| mHFD-PBS | D8M1 | 6.06E-01 | 6.88E-01 | 6.06E-01 | 7.14E-01 | 6.96E-01 | 8.19E-01 | 5.35E-01 | 0.063077082 | 0.224564567 | 0.056231724 | 0.078401696 | 0.068843126 | 0.186243169 | 0.274032034 | 0.030511498 |
| mHFD-MEV | D2M4 | 7.61E-01 | 8.14E-01 | 7.61E-01 | 9.32E-01 | 1.00E+00 | 1.17E+00 | 8.66E-01 | 0.208961193 | 0.112954941 | 0.022564083 | 0.001669982 | 0.040804658 | 0.075087958 | 0.222306111 | 0.002113368 |
| mHFD-MEV | D4M3 | 6.61E-01 | 9.75E-01 | 6.61E-01 | 8.25E-01 | 7.56E-01 | 8.09E-01 | 4.60E-01 | 0.156615215 | 0.214195205 | 0.308910701 | 0.029045834 | 0.081917435 | 0.108044982 | 0.096946201 | 0.046571579 |
| mHFD-MEV | D6M3 | 3.48E-01 | 8.54E-01 | 3.48E-01 | 1.03E+00 | 6.60E-01 | 4.02E-01 | 3.82E-01 | 0.124299172 | 0.179214995 | 1.145822243 | 0.033765479 | 0.058464197 | 0.212160238 | 0.149260768 | 0.216572327 |
| mHFD-MEV | D8M4 | 4.79E-01 | 1.01E+00 | 4.79E-01 | 1.06E+00 | 6.06E-01 | 5.82E-01 | 4.51E-01 | 0.112853623 | 0.206815279 | 0.106386606 | 0.093965977 | 0.114651333 | 0.063706728 | 0.271874607 | 0.230618794 |
| mHFD-handle | D2M7 | 3.73E-01 | 6.47E-01 | 3.73E-01 | 8.28E-01 | 4.04E-01 | 3.80E-01 | 2.99E-01 | 0.024660283 | 0.202836939 | 0.43666131 | 0.326203296 | 0.129713185 | 0.43264648 | 0.653858989 | 0.310062369 |
| mHFD-handle | D4M6 | 7.36E-01 | 9.66E-01 | 7.36E-01 | 1.08E+00 | 1.01E+00 | 8.36E-01 | 4.63E-01 | 0.089831491 | 0.120242325 | 0.328148018 | 0.499230899 | 0.187738525 | 0.575260542 | 0.618501452 | 0.199114616 |
| mHFD-handle | D6M5 | 5.58E-01 | 1.04E+00 | 5.58E-01 | 9.80E-01 | 7.20E-01 | 7.43E-01 | 3.72E-01 | 0.01666465 | 0.195921223 | 0.253702386 | 0.313411446 | 0.049322357 | 0.434752976 | 0.380141679 | 0.099026416 |
| mHFD-handle | D8M5 | 4.77E-01 | 5.96E-01 | 4.77E-01 | 6.43E-01 | 5.54E-01 | 1.03E+00 | 5.69E-01 | 0.153675403 | 0.193799218 | 0.095342545 | 0.445288705 | 0.016095865 | 0.643491759 | 0.777556648 | 0.375875215 |
| mCHD-PBS | D1F1 | 8.89E-01 | 9.69E-01 | 8.89E-01 | 1.11E+00 | 1.12E+00 | 7.99E-01 | 4.39E-01 | 0.388653404 | 0.259952729 | 0.298381266 | 0.240078774 | 0.131287175 | 0.257288251 | 0.117348358 | 0.058917037 |
| mCHD-PBS | D3F1 | 1.12E+00 | 8.90E-01 | 1.12E+00 | 1.27E+00 | 1.45E+00 | 1.17E+00 | 8.41E-01 | 0.323909198 | 0.3462385 | 0.348907756 | 0.103117755 | 0.006157191 | 0.03195862 | 0.16735979 | 0.152189164 |
| mCHD-PBS | D9F1 | 5.92E-01 | 5.90E-01 | 5.92E-01 | 1.21E+00 | 5.48E-02 | 4.06E-01 | 6.71E-02 | 0.220824185 | 0.221348505 | 0.276783162 | 0.1159132 | 0.092322116 | 0.330934272 | 0.155162339 | 0.06333358 |
| mCHD-PBS | D11F2 | 1.14E+00 | 1.18E+00 | 1.14E+00 | 1.09E+00 | 1.67E+00 | 1.26E+00 | 1.22E+00 | 0.126205764 | 0.250475786 | 0.289804546 | 0.10695495 | 0.095549099 | 0.403613821 | 0.096065418 | 0.107627042 |
| mCHD-MEV | D1F3 | 1.70E+00 | 9.48E-01 | 1.70E+00 | 1.05E+00 | 1.09E+00 | 7.65E-01 | 3.90E-01 | 0.040379974 | 0.080597447 | 0.282792513 | 0.203104136 | 0.219335461 | 0.400198559 | 0.195700858 | 0.179399074 |
| mCHD-MEV | D3F5 | 1.16E+00 | 8.01E-01 | 1.16E+00 | 1.24E+00 | 1.73E+00 | 1.51E+00 | 7.96E-01 | 0.010325202 | 0.086811678 | 0.28266613 | 0.475084769 | 0.105282974 | 0.365900299 | 0.177143757 | 0.35973073 |
| mCHD-MEV | D9F3 | 1.07E+00 | 1.17E+00 | 1.07E+00 | 1.30E+00 | 4.12E-01 | 1.35E+00 | 9.53E-01 | 0.115097607 | 0.351000177 | 0.29617462 | 0.419994997 | 0.205641244 | 0.372865086 | 0.128916528 | 0.201955733 |
| mCHD-MEV | D11F4 | 1.32E+00 | 9.30E-01 | 1.32E+00 | 1.31E+00 | 1.53E+00 | 1.24E+00 | 7.76E-01 | 0.047202675 | 0.241890576 | 0.31104338 | 0.265364812 | 0.178075049 | 0.313157916 | 0.204819434 | 0.087589343 |
| mCHD-handle | D1F6 | 1.12E+00 | 9.85E-01 | 1.12E+00 | 1.18E+00 | 2.13E+00 | 1.85E+00 |  | 0.207873924 | 0.335812716 | 0.323184484 | 0.223001923 | 0.220161356 | 0.360560349 | 0.134165075 | 0.157564402 |
| mCHD-handle | D3F7 | 1.22E+00 | 9.01E-01 | 1.22E+00 | 1.17E+00 | 1.54E+00 | 1.26E+00 | 6.65E-01 | 0.205580336 | 0.356463562 | 0.282364884 | 0.198812549 | 0.149539029 | 0.474753206 | 0.219208118 | 0.065831086 |
| mCHD-handle | D9F5 | 8.93E-01 | 8.60E-01 | 8.93E-01 | 1.13E+00 | 5.81E-01 | 1.11E+00 | 6.04E-01 | 0.113172554 | 0.099261593 | 0.30567742 | 0.105341039 | 0.002615125 | 0.581675472 | 0.222259638 | 0.220803392 |
| mCHD-handle | D11F6 | 1.25E+00 | 7.96E-01 | 1.25E+00 | 1.19E+00 | 6.07E-01 | 8.11E-01 | 6.92E-01 | 0.12312223 | 0.289129259 | 0.330630261 | 0.070196494 | 0.260682435 | 0.490006886 | 0.239984972 | 0.258026471 |
| mHFD-PBS | D2F2 | 4.87E-01 | 6.55E-01 | 4.87E-01 | 1.17E+00 | 5.21E-01 | 7.32E-01 | 2.74E-01 | 0.231435581 | 0.100651912 | 0.02675523 | 0.04602565 | 0.239872682 | 0.454126993 | 0.05900455 | 0.19264985 |
| mHFD-PBS | D4F1 | 1.32E+00 | 9.59E-01 | 1.32E+00 | 1.31E+00 | 1.23E+00 | 1.34E+00 | 8.32E-01 | 0.138310448 | 0.17382142 | 1.044713178 | 0.072821397 | 0.234161 | 0.5321843 | 0.040801301 | 0.234848594 |
| mHFD-PBS | D6F1 | 7.85E-01 | 8.50E-01 | 7.85E-01 | 1.19E+00 | 8.18E-01 | 9.54E-01 | 4.32E-01 | 0.125086053 | 0.097020255 | 0.735111368 | 0.111801712 | 0.332122889 | 0.742142337 | 0.081663366 | 0.316882738 |
| mHFD-PBS | D8F2 | 7.49E-01 | 8.56E-01 | 7.49E-01 | 9.45E-01 | 9.91E-01 | 6.96E-01 | 4.92E-01 | 0.124937046 | 0.122958341 | 0.195651129 | 0.150190277 | 0.366372327 | 0.810487561 | 0.090264279 | 0.243849648 |
| mHFD-MEV | D2F3 | 6.17E-01 | 6.72E-01 | 6.17E-01 | 1.12E+00 | 9.19E-01 | 5.44E-01 | 4.18E-01 | 0.306385789 | 0.270542816 | 0.161070846 | 0.144419465 | 0.244482395 | 0.948198741 | 0.100565935 | 0.149776699 |
| mHFD-MEV | D4F4 | 7.30E-01 | 8.25E-01 | 7.30E-01 | 1.30E+00 | 1.58E+00 | 9.34E-01 | 8.93E-01 | 0.064873286 | 0.343652116 | 0.64811255 | 0.037975505 | 0.29100871 | 0.862805252 | 0.132688758 | 0.053336184 |
| mHFD-MEV | D6F4 | 6.88E-01 | 7.91E-01 | 6.88E-01 | 1.04E+00 | 4.49E-01 | 7.21E-01 | 4.01E-01 | 0.122691482 | 0.163025884 | 1.285874293 | 0.018864584 | 0.365788887 | 0.657382541 | 0.125494173 | 0.174155635 |
| mHFD-MEV | D8F4 | 8.08E-01 | 8.88E-01 | 8.08E-01 | 1.12E+00 | 9.93E-01 | 8.25E-01 | 5.98E-01 | 0.424769593 | 0.400760525 | 0.047115267 | 0.092665618 | 0.286511057 | 0.854611908 | 0.184830722 | 0.106095908 |
| mHFD-handle | D2F5 | 9.78E-01 | 9.39E-01 | 9.78E-01 | 1.28E+00 | 1.16E+00 | 9.39E-01 | 6.80E-01 | 0.579695446 | 0.343692959 | 0.116451639 | 0.329831948 | 0.350595506 | 0.804461048 | 0.140373818 | 0.03104934 |
| mHFD-handle | D4F6 | 8.25E-01 | 9.60E-01 | 8.25E-01 | 1.29E+00 | 9.58E-01 | 6.30E-01 | 6.01E-01 | 0.502818455 | 0.347041062 | 0.388549513 | 0.508742674 | 0.501247734 | 0.68261642 | 0.157064656 | 0.125913622 |
| mHFD-handle | D6F6 | 6.76E-01 | 8.93E-01 | 6.76E-01 | 1.01E+00 | 8.86E-01 | 7.46E-01 | 4.93E-01 | 0.728375939 | 0.435776819 | 0.236152762 | 0.53309722 | 0.164459528 | 0.942257494 | 0.163177244 | 0.135106904 |
| mHFD-handle | D8F5 | 6.35E-01 | 9.79E-01 | 6.35E-01 | 1.48E+00 | 8.79E-01 | 8.03E-01 | 4.32E-01 | 0.791736823 | 0.536630536 | 0.01600914 | 0.7905841 | 0.376664911 | 1.201292217 | 0.127145571 | 0.501201077 |

**Table 5**: Normalized transcript abundance data and normalized protein abundance data for NF-kb pathway targets in rat hypothalamus

| Treatment groups | Sample IDs | Normalized transcript abundance | | | | | | | Sample IDs | Normalized protein abundance | | | | | | | |
| --- | --- | --- | --- | --- | --- | --- | --- | --- | --- | --- | --- | --- | --- | --- | --- | --- | --- |
|  |  | TLR4 | Iκκ β | IκBα | NF--κB | NLRP3 | Caspase-1 | IL-18 |  | TLR4 | P-Iκκ β (S176/180) | P-IκBα (S32) | NF-κB | P-NF-κB (S536) | NLRP3 | Pro-caspase-1 | Cleaved-caspase-1 |
| mCHD-PBS | D1M1 | 8.40E-01 | 1.70E+00 | 1.14E-01 | 1.58E-01 | 7.83E-01 | 1.02E+00 | 3.59E-01 | D1M2 | 0.52 | 3.75 | 0.411270737 | 1.526435156 | 1.107141579 | 0.089709267 | 0.118242227 | 0.259364073 |
| mCHD-PBS | D3M1 | 9.47E-01 | 1.18E+00 | 6.72E-02 | 9.13E-02 | 7.63E-01 | 9.78E-01 | 3.33E-01 | D9M2 | 0.24 | 3.20 | 0.12290719 | 0.976878537 | 0.759395863 | 0.098146071 | 0.076997733 | 0.1308741 |
| mCHD-PBS | D9M1 | 5.51E-01 | 1.34E+00 | 1.01E-01 | 1.37E-01 | 5.91E-01 | 7.29E-01 | 1.68E-01 | D11M1 | 0.47 | 4.19 | 0.019462694 | 1.092065555 | 1.195135235 | 0.095848465 | 0.023062134 | 0.098410449 |
| mCHD-PBS | D11M2 | 1.12E+00 | 1.60E+00 | 1.17E-01 | 1.29E-01 | 8.14E-01 | 1.02E+00 | 3.20E-01 |  |  |  |  |  |  |  |  |  |
| mCHD-MEV | D1M4 | 8.26E-01 | 1.25E+00 | 1.04E-01 | 1.29E-01 | 8.48E-01 | 8.16E-01 | 2.01E-01 | D1M3 | 2.98 | 8.64 | 0.15415478 | 1.912770726 | 1.193098609 | 0.439988858 | 0.197681661 | 0.075580633 |
| mCHD-MEV | D3M2 | 8.38E-01 | 1.81E+00 | 1.44E-01 | 1.58E-01 | 1.19E+00 | 1.01E+00 | 1.78E-01 | D9M5 | 3.51 | 5.39 | 0.152476445 | 0.912482784 | 0.767488601 | 0.241081956 | 0.124972488 | 0.044091093 |
| mCHD-MEV | D9M4 | 6.57E-01 | 1.27E+00 | 9.91E-02 | 1.20E-01 | 7.01E-01 | 8.57E-01 | 1.98E-01 | D11M4 | 4.58 | 3.35 | 0.257076608 | 0.694775076 | 0.952552119 | 0.179018253 | 0.159067853 | 0.196923141 |
| mCHD-MEV | D11M3 | 6.52E-01 | 1.22E+00 | 1.13E-01 | 1.03E-01 | 7.28E-01 | 5.81E-01 | 2.36E-01 |  |  |  |  |  |  |  |  |  |
| mCHD-handle | D1M6 | 8.77E-01 | 1.47E+00 | 1.19E-01 | 1.34E-01 | 7.40E-01 | 1.07E+00 | 3.94E-01 | D1M5 | 4.74 | 5.66 | 0.178011175 | 1.428609521 | 1.596474314 | 0.430582028 | 0.222612666 | 0.058176717 |
| mCHD-handle | D3M3 | 8.26E-01 | 1.05E+00 | 8.77E-02 | 1.17E-01 | 5.58E-01 | 7.53E-01 | 2.89E-01 | D9M7 | 5.01 | 4.19 | 0.37414283 | 0.833315663 | 1.06527368 | 1.193420987 | 0.220896816 | 0.12026756 |
| mCHD-handle | D9M6 | 8.20E-01 | 1.46E+00 | 1.12E-01 | 1.13E-01 | 8.84E-01 | 9.21E-01 | 2.37E-01 | D11M5 | 2.56 | 3.22 | 0.110093788 | 0.502014855 | 0.772242745 | 1.044995029 | 0.163990944 | 0.094806458 |
| mCHD-handle | D11M6 | 6.90E-01 | 1.17E+00 | 9.89E-02 | 1.00E-01 | 5.71E-01 | 6.50E-01 | 2.68E-01 |  |  |  |  |  |  |  |  |  |
| mHFD-PBS | D2M2 | 9.96E-01 | 9.73E-01 | 2.28E+00 |  |  |  |  | D2M1 | 1.60 | 3.79 | 0.055652434 | 0.914066552 | 1.657170849 | 0.742182342 | 0.390444889 | 0.321680935 |
| mHFD-PBS | D4M1 | 6.36E-01 | 1.37E+00 | 1.84E-01 | 2.28E-01 | 8.89E-01 | 6.26E-01 | 4.65E-01 | D4M2 | 1.36 | 3.87 | 0.02046511 | 0.561737501 | 1.729878488 | 0.444003043 | 0.303493882 | 0.313450009 |
| mHFD-PBS | D6M1 | 6.15E-01 | 1.10E+00 | 1.58E-01 | 1.96E-01 | 6.53E-01 | 8.17E-01 | 2.03E-01 | D6M2 | 0.96 | 4.59 | 0.026377111 | 0.737929149 | 2.217147148 | 0.742182342 | 0.268062585 | 0.174991805 |
| mHFD-PBS | D8M1 | 9.10E-01 | 1.24E+00 | 1.61E-01 | 1.78E-01 | 5.54E-01 | 6.68E-01 | 2.15E-01 | D8M2 | 0.10 | 4.49 | 0.03476541 | 0.858870234 | 1.974534334 | 0.444003043 | 0.240376185 | 0.19418227 |
| mHFD-MEV | D2M4 | 7.39E-01 | 1.42E+00 | 1.63E-01 | 2.03E-01 | 7.75E-01 | 9.16E-01 | 2.66E-01 | D2M3 | 0.18 | 3.87 | 0.089898437 | 0.687018657 | 1.827224428 | 1.004080039 | 0.021027572 | 0.345438077 |
| mHFD-MEV | D4M3 | 5.46E-01 | 1.36E+00 | 2.32E-01 | 1.48E-01 | 7.82E-01 | 8.03E-01 | 2.06E-01 | D4M4 | 0.37 | 3.29 | 0.062598474 | 0.664941941 | 1.899449616 | 0.230845392 | 0.011245855 | 0.37960924 |
| mHFD-MEV | D6M3 | 7.32E-01 | 1.41E+00 | 1.94E-01 | 2.41E-01 | 7.87E-01 | 8.63E-01 | 2.11E-01 | D6M4 | 0.83 | 1.99 | 0.106910468 | 0.325593986 | 1.163518658 | 0.287213559 | 0.02544551 | 0.411778361 |
| mHFD-MEV | D8M4 | 5.34E-01 | 1.14E+00 | 1.51E-01 | 2.20E-01 | 7.97E-01 | 8.55E-01 | 2.72E-01 | D8M3 | 1.43 | 2.26 | 0.068987874 | 0.486900821 | 1.004623198 | 0.621781792 | 0.022303811 | 0.538119158 |
| mHFD-handle | D2M7 | 8.43E-01 | 1.47E+00 | 1.62E-01 | 2.07E-01 | 7.18E-01 | 7.39E-01 | 4.87E-01 | D2M6 | 1.15 | 3.85 | 0.011946607 | 1.54429068 | 1.401159018 | 1.330600809 | 0.054691746 | 0.637084495 |
| mHFD-handle | D4M6 | 7.97E-01 | 1.21E+00 | 1.27E-01 | 1.10E-01 | 9.68E-01 | 7.48E-01 | 3.07E-01 | D4M6 | 0.99 | 6.08 | 0.176751259 | 2.929836387 | 1.35229075 | 2.646554997 | 0.145985469 | 0.568844641 |
| mHFD-handle | D6M5 | 7.21E-01 | 1.24E+00 | 1.33E-01 | 2.08E-01 | 8.00E-01 | 7.69E-01 | 3.30E-01 | D6M6 | 1.49 | 3.32 | 0.214682337 | 1.130749822 | 0.714920203 | 2.090246686 | 0.161735817 | 0.421990911 |
| mHFD-handle | D8M5 | 5.88E-01 | 1.25E+00 | 1.18E-01 | 1.30E-01 | 6.13E-01 | 4.74E-01 | 4.42E-01 | D8M6 |  | 2.91 | 0.098886538 | 0.944015636 | 0.481762347 | 1.591497148 | 0.459815319 | 0.388557282 |
| mCHD-PBS | D1F1 | 6.79E-01 | 1.08E+00 | 5.72E-02 | 1.04E-01 | 7.07E-01 | 7.95E-01 | 1.95E-01 | D1F2 | 2.51 | 1.12 | 0.337225265 | 0.27514391 | 0.905070188 | 0.836892417 | 0.319860664 | 0.112198425 |
| mCHD-PBS | D3F1 | 8.93E-01 | 1.17E+00 | 1.06E-01 | 1.03E-01 | 1.00E+00 | 7.25E-01 | 2.07E-01 | D3F2 | 2.41 | 0.68 | 0.364036806 | 0.171870276 | 0.394002674 | 0.731149056 | 0.249647816 | 0.091156254 |
| mCHD-PBS | D9F1 | 7.08E-01 | 1.35E+00 | 1.37E-01 | 1.35E-01 | 8.05E-01 | 8.17E-01 | 1.94E-01 | D9F2 | 3.62 | 0.85 | 0.276100251 | 0.133863395 | 0.352168387 | 1.145345617 | 0.198528969 | 0.078934663 |
| mCHD-PBS | D11F2 | 5.27E-01 | 1.14E+00 | 1.43E-01 | 1.11E-01 | 9.36E-01 | 8.22E-01 | 2.77E-01 | D11F1 | 2.71 | 1.02 | 0.453739846 | 0.120029512 | 0.281765969 | 0.75838333 | 0.18842709 | 0.075482939 |
| mCHD-MEV | D1F3 | 7.81E-01 | 1.47E+00 | 1.43E-01 | 1.14E-01 | 8.08E-01 | 9.62E-01 | 2.13E-01 | D1F4 | 1.85 | 0.65 | 0.412740541 | 0.249864749 | 0.008800055 | 0.322291493 | 0.366769159 | 0.571258417 |
| mCHD-MEV | D3F5 | 8.30E-01 | 1.22E+00 | 1.09E-01 | 1.23E-01 | 7.22E-01 | 8.66E-01 | 1.99E-01 | D3F4 | 4.28 |  | 0.222484122 |  | 0.144019013 | 1.002256203 | 0.476337821 | 0.137430898 |
| mCHD-MEV | D9F3 | 7.52E-01 | 1.30E+00 | 9.34E-02 | 1.24E-01 | 1.09E+00 | 7.60E-01 | 2.65E-01 | D9F4 | 3.93 | 1.20 | 0.162669803 | 0.336797757 | 0.018958779 | 1.030671872 | 0.371959999 | 0.237296173 |
| mCHD-MEV | D11F4 | 6.89E-01 | 1.32E+00 | 1.11E-01 | 1.13E-01 | 7.82E-01 | 8.74E-01 | 1.83E-01 | D11F3 | 4.54 | 1.16 | 0.038342901 | 0.318985722 | 0.218134892 | 1.235802588 | 0.386648983 | 0.394142395 |
| mCHD-handle | D1F6 | 7.33E-01 | 1.31E+00 | 6.42E-02 | 8.72E-02 | 8.39E-01 | 1.01E+00 | 1.45E-01 | D1F5 | 3.22 | 1.24 | 0.123459192 | 0.40970194 | 0.319380701 | 0.98793834 | 0.365261132 | 0.69408167 |
| mCHD-handle | D3F7 | 9.03E-01 | 1.51E+00 | 1.19E-01 | 1.18E-01 | 8.37E-01 | 7.47E-01 | 2.36E-01 | D3F6 | 4.01 | 1.39 | 0.027540932 | 0.40475337 | 0.349123313 | 1.193624982 | 0.514880358 | 0.738372103 |
| mCHD-handle | D9F5 | 8.82E-01 | 1.51E+00 | 1.05E-01 | 1.49E-01 | 9.64E-01 | 9.67E-01 | 2.72E-01 | D11F5 | 4.92 | 1.31 | 0.016109774 | 0.381303634 | 0.441893718 | 1.569658462 | 0.571598136 | 0.31503209 |
| mCHD-handle | D11F6 | 9.91E-01 | 1.44E+00 | 8.76E-02 | 1.08E-01 | 8.14E-01 | 9.67E-01 | 2.24E-01 |  |  |  |  |  |  |  |  |  |
| mHFD-PBS | D2F2 | 6.82E-01 | 1.32E+00 | 5.66E-02 | 1.57E-01 | 7.63E-01 | 7.05E-01 | 2.99E-01 | D2F1 | 1.22 | 5.69 | 2.007778566 | 0.823582544 | 1.459626881 | 0.985420546 | 0.767576619 | 0.315206782 |
| mHFD-PBS | D4F1 | 9.79E-01 | 1.43E+00 | 2.66E-01 | 3.04E-01 | 1.04E+00 | 1.08E+00 | 6.11E-01 | D4F2 | 1.57 | 2.78 | 2.152306838 | 0.644227247 | 1.111750618 | 0.56689065 | 0.34126754 | 0.320276718 |
| mHFD-PBS | D6F1 | 7.77E-01 | 1.46E+00 | 1.82E-01 | 2.13E-01 | 1.05E+00 | 8.18E-01 | 3.90E-01 | D6F2 | 1.12 | 3.42 | 1.868713409 | 0.899231021 | 1.143417508 | 0.051037058 | 0.347561716 | 0.399218475 |
| mHFD-PBS | D8F2 | 4.78E-01 | 1.15E+00 | 6.18E-02 | 1.83E-01 | 7.30E-01 | 5.80E-01 | 3.62E-01 | D8F1 | 0.26 | 4.08 | 1.847879011 | 0.84336647 | 1.189332903 | 0.462105613 | 0.302757457 | 0.475752366 |
| mHFD-MEV | D2F3 | 1.04E+00 | 1.88E+00 | 2.65E-01 | 2.70E-01 | 8.87E-01 | 9.92E-01 | 6.31E-01 | D2F4 | 0.64 | 3.88 | 2.530764574 | 0.709153037 | 1.118012006 | 0.945089003 | 0.520926346 | 0.857473731 |
| mHFD-MEV | D4F4 | 9.56E-01 | 1.94E+00 | 1.31E-01 | 2.10E-01 | 1.07E+00 | 9.29E-01 | 3.93E-01 | D4F3 | 1.00 | 3.68 | 2.213291908 | 0.50851156 | 1.332772567 | 0.915906963 | 0.229010492 | 0.572540482 |
| mHFD-MEV | D6F4 | 3.97E-02 | 1.70E+00 | 1.85E-01 | 2.80E-01 | 9.81E-01 | 6.59E-01 | 7.15E-01 | D6F3 | 1.31 | 1.85 | 2.454114152 | 0.139335282 | 0.917921 | 1.104618248 | 0.468800354 | 0.698458033 |
| mHFD-MEV | D8F4 | 6.81E-01 | 1.41E+00 | 1.48E-01 | 1.75E-01 | 7.84E-01 | 7.33E-01 | 5.38E-01 | D8F3 | 1.00 | 3.79 | 0.558094909 | 0.536255189 |  | 0.366296696 | 0.456978844 | 0.993915698 |
| mHFD-handle | D2F5 | 7.75E-01 | 1.87E+00 | 1.07E-01 | 1.44E-01 | 1.23E+00 | 1.09E+00 | 4.97E-01 | D4F5 | 1.47 | 3.14 | 2.519484444 | 1.108271066 | 1.074682445 | 1.651239981 | 0.302093975 | 0.274929449 |
| mHFD-handle | D4F6 | 4.87E-02 | 1.79E+00 | 7.71E-02 | 1.50E-01 | 9.95E-01 | 8.33E-01 | 3.55E-01 | D6F5 | 1.90 | 4.22 | 1.40819767 | 1.073914409 | 1.097283667 | 1.894177453 | 0.318082585 | 0.143065187 |
| mHFD-handle | D6F6 | 7.90E-01 | 1.80E+00 | 8.35E-02 | 1.93E-01 | 1.06E+00 | 7.57E-01 | 9.19E-01 | D8F6 | 1.19 | 1.81 | 1.738654872 | 0.418033557 | 0.821109894 | 1.548771426 | 0.279258569 | 0.111544376 |
| mHFD-handle | D8F5 | 6.76E-01 | 1.68E+00 | 1.68E-01 | 2.40E-01 | 9.53E-01 | 7.04E-01 | 8.90E-01 |  |  |  |  |  |  |  |  |  |
